# Contextualized phenovectors reveal a Drosophila serotonergic circuit controlling satiety

**DOI:** 10.64898/2026.07.01.735755

**Authors:** Sangyu Xu, Xianyuan Zhang, James C. Stewart, Joses Ho, Deepak Choudhury, Zhiping Wang, Adam Claridge-Chang

**Affiliations:** Institute for Molecular and Cell Biology (IMCB), Agency for Science, Technology and Research (A*STAR), Singapore, Singapore; Program in Neuroscience and Behavioral Disorders, Duke-NUS Medical School, Singapore; Singapore Institute of Manufacturing Technology, Agency for Science, Technology and Research (A*STAR), Singapore, Singapore; Department of Physiology, Yong Loo Lin School of Medicine, National University of Singapore, Singapore

## Abstract

Distinguishing whether a neural circuit genuinely controls a holistic behavioral state, rather than merely producing isolated effects, is a central challenge in neuroscience. Feeding research illustrates the problem: decades of work have reached conflicting conclusions partly because individual behavioral features are assessed in isolation. Here we establish a contextualized ethomics approach that benchmarks optogenetic circuit manipulations against natural hunger–satiety transitions, using high-dimensional behavioral tracking in *Drosophila* and starvation as a ground-truth intervention. Applying this framework, we find that serotonergic neurons marked by the *Tryptophan hydroxylase neuronal* (Trhn) enhancer are both required for and instructive of a satiety-like state, whereas other feeding-related circuits produce only fragmentary behavioral changes. Intersectional dissection localized this control to Trhn neurons of the ventral nerve cord, which act through the sugar transporter Sut2 to sense nutrient state. This work strengthens our understanding of serotonergic feeding control and provides a generalizable framework for validating circuit–state relationships.

## Introduction

Understanding neural circuit control of internal states is a central goal in neuroscience. Feeding behavior provides an interesting and tractable model for studying state transitions, as hunger-satiety alternation represents one of the most ancient and fundamental behavioral states (*1*, *2*) and can be easily manipulated. To transit the hunger–satiety axis, the nervous system requires signals that can translate nutrient status into orchestrated circuit and behavioral changes. While neural manipulations can alter feeding behavior, measuring single behavioral phenotypes risks mistaking isolated effects for state control. Identifying which neural populations authentically control these broadly coordinated state transitions, as opposed to producing dissociated behavioral effects, remains a core challenge. However, no framework exists to systematically evaluate whether circuit manipulations produce authentic state changes or merely isolated behavioral effects.

Various neuromodulatory systems have been identified as drivers of feeding behavior, yet their roles in hunger state control remain unclear. Given their broad projections and diverse effects, these systems are likely to be key mediators of the hunger–satiety state transition (*3–6*). Serotonin (5-hydroxytryptamine, 5-HT) is an ancient biomolecule and a highly conserved neuromodulator (*7–9*) that regulates nearly every aspect of animal physiology and behavior (*10*, *11*). The broad projection patterns of serotonergic neurons are suggestive of a system that is involved in signal broadcasting (*12*). Mammalian data suggest that serotonin predominantly serves as a satiation signal, suppressing food consumption and promoting satiety-related behaviors (*13–16*). The function of serotonin in invertebrates, however, is divergent, promoting behaviors connected to hunger, consumption, or satiety in different circuits across species (*17–24*).

This functional diversity of serotonin, even within a single species, is evident in the vinegar fly, *Drosophila melanogaster*, where both appetizing and satiating effects of serotonergic signaling have been observed. One study showed that the serotonergic neurons captured by the *Tryptophan hydroxylase neuronal* (*Trhn*) enhancer reduce feeding (*20*), while other studies have shown that neurons specified by the *Serotonin transporter* (*SerT*) upstream region increase feeding (*21*, *22*). This phenomenon raises the question as to whether such discrepant effects of serotonin might reflect distinct modules operating at different feeding phases, or if they can be reconciled into a common framework of how serotonin regulates feeding state.

Flies provide an advantageous system for dissecting serotonergic feeding control, with sophisticated genetic tools, large sample sizes, and conserved molecular pathways. As flies eat nanoliter-scale meals, weighing the feed is not possible, so surrogate measures, such as proboscis extension, reporter dyes, radioactive tracers, olfactory responses, feeding latency, and food contact, are typically used (*25*). The capillary feeder (CAFE) assay and recent advances in automated tracking of food intake in flies measured feed volume more directly (*26*, *27*). However, measuring food consumption or adjacent metrics in isolation cannot distinguish interventions that specifically impact appetite from those that affect feeding as an epiphenomenon of other changes (*28*).

Since authentic internal states like hunger and satiety govern multiple, coordinated behavioral patterns(*2*, *29*), validating circuit function requires measuring multiple behavioral features simultaneously. Automation and machine vision can facilitate this integrative approach being applied to numerous neurogenetic interventions, in an approach termed *ethomics* (*30*), which has been developed in several species (*31*), including *Drosophila* circuit analysis (*32*, *33*). Building on automated feed-tracking approaches, we developed a system that tracks food consumption volume and locomotion with high temporal resolution and facilitates optogenetic interventions in neuronal function during a feeding experiment.

Here we aimed to understand how serotonergic neurons encode and regulate the hunger–satiety axis using a contextualized ethomics approach that benchmarks circuit manipulations against natural state transitions. We developed and tested an automated feed-tracking system to analyze multiple behavioral features during *Drosophila* hunger and satiety, using starvation as the ground truth of hunger to evaluate whether optogenetic neuronal manipulations produce merely dissociated behavioral effects or authentic state changes. We term this approach “**d**elta **e**thomic **s**tate-**t**ransition **r**ecapitulation **a**ssessment,” DESTRA. This method allowed us to identify serotonergic populations that are both instructive to and required for satiety while revealing that many other purported “feeding circuits” fail to recapitulate authentic hunger states, operationally defined as authentic-hunger-like state (AHLS). Neurogenetics revealed how a specific circuit in the nerve cord links internal nutritional state to organized behavioral output, demonstrating a fundamental mechanism by which a serotonergic circuit translates metabolic signals into coherent hunger and satiety states. The neurogenetic DESTRA approach provides a systematic method for evaluating which neural systems have authentic control of internal states.

## Results

### Espresso measures hunger-dependent changes in feeding and locomotion

To understand feeding and associated functions of serotonin neuronal activities, we developed a high-throughput multimodal feeding assay, referred to as *Espresso*. We first aimed to test the functionality of the Espresso system (**Figure 1A**) during a 2-h refeeding experiment, to establish feeding and locomotion parameters in single wild-type flies (**Figure 1B**). A glass capillary positioned above each arena provided liquid food containing sugar and yeast extract; when the centroid of a fly was present in an enabled food port, the Espresso software successfully recorded a feeding event and quantified the capillary liquid level (**Figure 1C**, **Methods**). To test whether Espresso could detect the difference between various hunger states, we starved flies for 0, 24, and 48 hours (**Figure 1D**) and then tracked their food consumption and locomotion over 2 hours of refeeding (**Figure 1 E-G**). Here, we saw increased total feed volumes by 86 nL/fly and 270 nL/fly after 24- and 48-h starvation (**Figure 1H**), respectively. We also detected robust effects of starvation on other behavioral features, namely increased meal size, meal duration, and feeding speed (**Figure 1I**). After a meal, average walking speed decreased by approximately 0.37 mm/s (**Figure 1J - L**), verifying previous reports (*27*, *34*). In the post-meal epoch (red), the fly’s trajectory sometimes includes a rapid excursion toward the control (empty) port (**Figure 1J**), suggesting that flies may revisit potential food locations after feeding in a behavior reminiscent of spatial memory or conditioned place preference. Starved flies typically spent more time near or inside ports with food-containing capillaries (**Figure 1M, N**), and less time near empty-capillary control ports (**Figure 1O**). We thus conclude that Espresso provided adequate sensitivity for detecting starvation-induced feeding changes.

**Figure 1.**
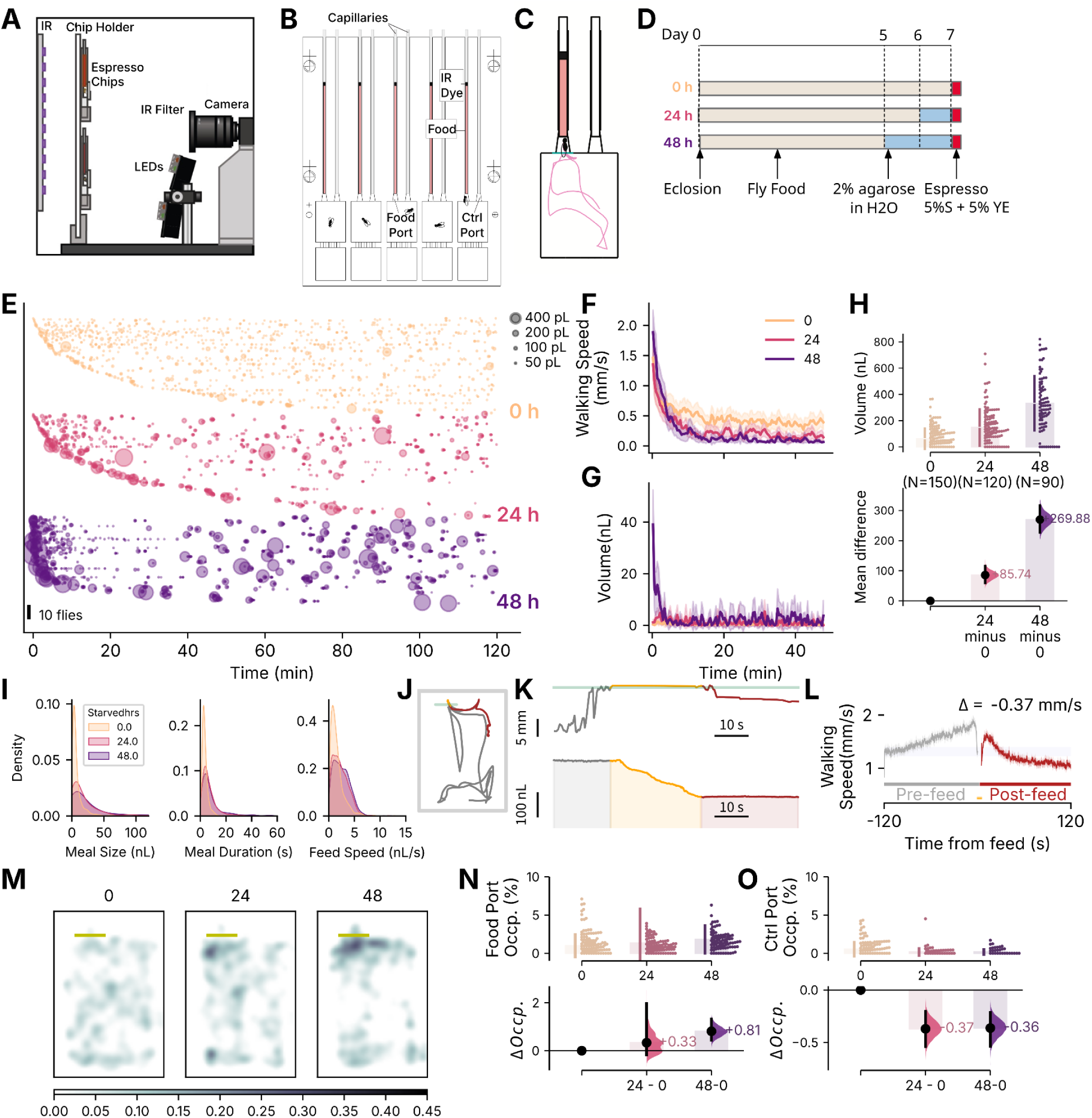
Espresso enables tracking of concurrent feeding and locomotion. **A,** Schematic of the Espresso assembly designed to track feeding and locomotion, and provide illumination for optogenetic experiments. The assembly comprises the following components: an infrared (IR) illuminator, acrylic chips integrating capillaries and behavior arena, a chip holder, red and green optogenetic LEDs, and a camera with an infrared filter. **B,** Schematic of an Espresso chip, with five behavior chambers, ten channels with ten capillaries capable of holding liquid food mounted, and one fly per chamber. At the top of each capillary, a drop of IR-opaque dye adds contrast for the IR imaging of food level. Each arena has two ports (Food port and Ctrl port), which allows analysis of the effect of food presence. **C,** In a single rectangular behavioral chamber, a representative trajectory (pink line) of a fly leading up to a feed event. The port’s threshold (green line) is used to gate when the software tracks feeding. The liquid food (red) is topped with a layer of IR-opaque dye (black). **D,** Timeline of the three feeding regimes, including fed flies (‘0-h starvation’), 24-h, and 48-h starvation. From eclosion from the pupal case at Day 0, the flies are raised on standard rich food. Flies are assayed in Espresso chips on Day 7, with the starved flies having spent one or two days on non-nutritional media (water gelled with 2% agarose). The assay’s liquid food was 5% sucrose (S) and 5% yeast extract (YE) in water. **E,** Bubble plot showing feeding events, at different starvation levels (0-h/fed, 24-h, 48-h, orange, pink, and purple, respectively). Each bubble represents one meal, with its area proportional to the volume consumed (scale bubbles at 50, 100, 200, and 400 picoliters). **F,** Line plot showing mean walking speeds of the three nutritional states over the assay’s first 50 min. Solid lines indicate the mean volume consumed; the shaded ribbon indicates the 95% confidence interval (95CI). **G,** Line plot showing feed volume over time. Lines indicate the mean volume consumed; ribbons indicate 95CIs. **H,** A Cumming estimation plot showing total volume consumed in each chamber over the duration of the assay. In the swarm plot (upper), each dot denotes measurements from an individual fly, each vertical bar represents the group’s standard deviation, and the bar gap denotes the mean. The contrast plot (lower) shows the effect sizes: each black dot represents the mean difference; the black bar represents the 95CI; each half-violin curve represents the difference distribution; and numerals are the mean effect sizes. **I,** Kernel-density estimate plots showing the distributions of meal size (nL), meal duration (s), and feeding speed (nL/s) for individual meals in unstarved (0h, orange), 24-h starved (pink), or 48-h starved (purple) flies. **J,** Representative walking trajectory of a fly in two dimensions in a 90-s interval before (gray), during (yellow), and after (red) a meal. The gray box is the chamber bounds; the green line is the threshold to the feed port. **K**, Following the same trajectory as panel **J**, a fly’s elevation (y position in mm, top sub-panel) during the peri-feed interval and the corresponding capillary liquid level (nL, bottom sub-panel). The epochs are color-coded: gray = before feed; yellow = during feed; red = after feed. **L,** Line plot of the mean walking speed (mm/s) of flies in the 120 s before and 120 s after feeding. The numerals are the pre- and post-feeding speed difference (Δ). The shaded ribbon around each line indicates the 95CI. **M,** Heat maps showing the distribution of flies from the three nutrition regimes. Color-scale units are Z-scores of time spent (units of standard deviation). Yellow horizontal bars indicate positions of the virtual feed port. White scale bar = 2 mm. **N,** Estimation plot showing food-port occupancy for flies of the three nutritional states. The swarm plot (upper) shows observed values and the contrast plot (lower) summarizes starvation effect sizes relative to the fed state. Numerals next to the half violins indicate effect sizes. **O,** Estimation plot showing control-port occupancy; graphical elements as above. Adult male flies (5–10 days post-eclosion) were used throughout.

### *Trhn-Gal4* neurons modulate food consumption with bidirectional control of meal size

Having established that Espresso can reliably detect starvation-induced behavioral changes, we next investigated whether serotonergic neurons contribute to these natural hunger-satiety transitions. The role of serotonin in fly feeding is unclear. It was previously found that serotonin neurons identified by *Trhn-Gal4* driver (the *Trhn* enhancer fused to the Gal4 transcriptional activator; **Figure 2A**) suppress feeding (*20*). To further validate the Espresso system, we undertook (conceptual) replication of these experiments using optogenetics to activate or inhibit neurons driven by this driver and monitor feeding behavior. First, immunohistochemical stains confirmed that *Trhn*-*Gal4* drives expression in both the brain and VNC (**Figure 2A**). Next, to test the hypothesis that activity in *Trhn-Gal4* neurons suppresses feeding (*20*), we conducted reciprocal optogenetic manipulations: activating these neurons with Chrimson (Chr) in starved flies (**Figure 2C**) and inhibiting them with anion channelrhodopsin-1 (ACR1) in fed flies (**Figure 2 D, E**). As expected, in starved *Trhn>Chr* flies, activating the neurons reduced food intake (**Figure 2F**). This effect was not due to fewer meals (feed count; **Figure 2O**); rather, it was a result of markedly reduced meal sizes (ΔΔ = –75 nl; **Figure 2N**). By contrast, in fed *Trhn>ACR1* flies, inhibiting the neurons increased food consumption (**Figure 2G**), as evidenced by increased feed volume, feeding duration, and meal size (**Figure 2 L - O**). These feeding effects were consistent between male and female flies (**Figure S 1 A, B**). Together, these results establish that *Trhn-Gal4* activity has bidirectional control over consumption, which is achieved through changes in meal size.

### *R50H05-Gal4* activity can promote meal frequency, but does not mediate hunger drive

Other studies have shown that the serotonergic neurons captured by *R50H05-Gal4* (an enhancer fragment of the *SerT* gene fused to *Gal4*; hereafter ‘*R50H05’*; **Figure 2B**) do the opposite to *Trhn-Gal4*: they drive hunger and promote feeding (*21*, *22*). We aimed to verify the role of R50H05 activation on fly feeding. Immunohistochemical analyses confirmed that *R50H05* drives expression in a population in the brain that overlaps with *Trhn-Gal4* expression but is absent in the VNC (**Figure 2B**). To verify the hunger-promoting effects of *R50H05* activity (*21*, *22*), we tested their activation in fed flies (**Figure 2H**) and inhibition in starved, hungry flies (**Figure 2I**). Activation of *R50H05*>*Chr* with red light resulted in fed, sated flies eating more (**Figure 2J**), arising from an increase in feed count (**Figure 2O**) rather than meal size (**Figure 2N**). These effects were observed in both male and female flies (**Figure S 1C**). *R50H05* activation thus drives a general increase in food-seeking behavior rather than consumption. Inhibition of starved *R50H05*>*ACR1* flies with green light had no effect on the total feed volume (**Figure 2K**), or the other three features (duration, meal size and count; **Figure 2 L–O**). We thus conclude that while *R50H05* activation can increase feeding frequency in sated flies, the activity of these neurons is not essential for hunger-driven feeding behaviors.

These results confirm the opposing feeding effects of activated *Trhn-Gal4* versus *R50H05* neurons (*20*, *21*), and highlight that the two neural groups elicit opposing effects when inhibited. Interestingly, the changes in consumption result from different behavioral features: actuated *Trhn-Gal4* flies consumed smaller or larger meals (with *Chr* and *ACR1*, respectively), while actuated *R50H05>Chr* flies ate more frequently with no change in meal size.

**Figure 2.**
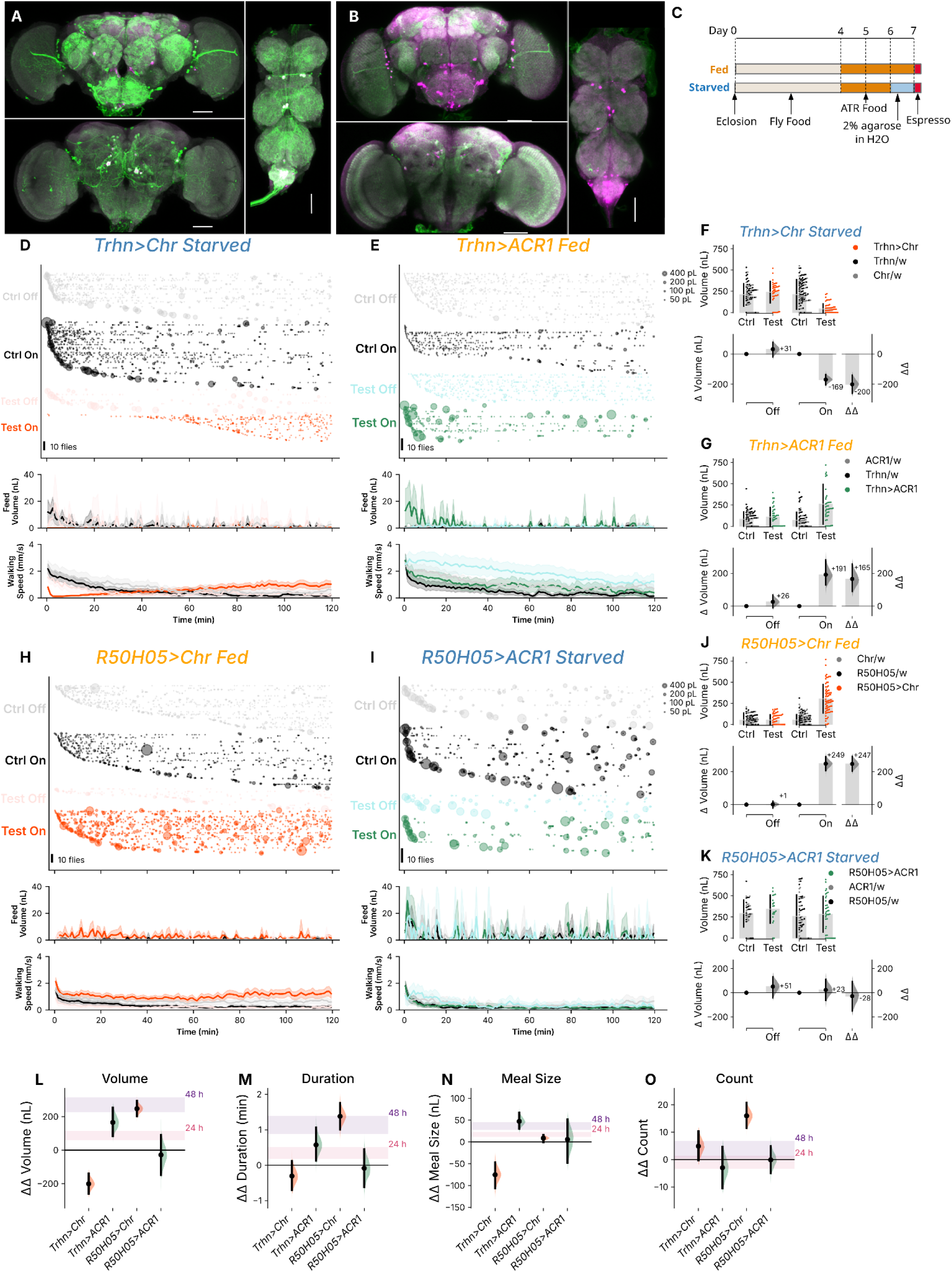
*Trhn-Gal4* neurons suppress feeding; *R50H05* neurons promote food seeking. **A**, Immunohistochemical staining of fly brains identifying the expression pattern of *Trhn-Gal4* driving *UAS*-*Chr*. Green, magenta, and gray indicate stains from antibodies α-GFP (Chr-YFP fusion), α-5-HT (neurotransmitter), and α-Brp (synaptic protein), respectively. Scale bar: 50 µm. **B,** Staining of *R50H05*-*Gal4* driving *UAS*-*Chr* (B). Graphical elements as above. **C,** Starvation protocol for optogenetic experiments: newly eclosed flies were kept on standard food for 4 days, then transferred to ATR food for two or three days. On day 6, starved flies were maintained on nutrient-free gel for 24 h, then assayed in Espresso for 2 h on day 7. **D,** Bubble plots (**top sub-panel**) for feeds over two hours of illumination of starved *Trhn*>*Chr* (red bubbles) and control (*UAS-Chr/+*, gray and *Trhn-Gal4*/+, black) flies. Each row of the bubble plot shows the timing of individual meals from a single fly: each bubble denotes one meal, and the area of the bubble is proportional to the volume of the meal. **Middle and lower sub-panels** show two line plots of mean feed volume (nL, middle) and mean walking speed (mm/s, bottom) over time; ribbons indicate 95CI. Sample sizes: N_Control light off_ = 81, _NTest light off_ = 39, N_Control light on_ = 121, N_Test light on_ = 59. **E**, Bubble plots and line charts for meals, feed volume, and speed in fed *Trhn*>*ACR1* and control flies (*UAS-ACR1/+*, gray and *Trhn-Gal4*/+, black). Graphical elements as above. Sample sizes: N_Control light off_ = 81, N_Test light off_ = 39, N_Control light on_ = 80, N_Test light on_ = 40) experiments. **F,** Estimation ΔΔ plot of starved *Trhn*>*Chr* flies and the control groups. The data are shown as a swarm plot (top) and a contrast plot (bottom) with the ΔΔ effect size (bottom right). The four conditions are, from left to right: controls (Ctrl) with lights off; test *Trhn*>*Chr* flies with lights off; controls with lights on; and test flies with lights on. In the contrast panel the Δ comparisons are for the lights-off groups, and lights-on groups, from left to right, and the ΔΔ effect that quantifies the [genotype × light] effect, i.e. the ‘opto-genetic’ effect. Numerals next to the half-violin curves are the effect-size means. See methods for full explanation of ΔΔ analysis. Sample sizes as above. **G**, Estimation plot for an optogenetic experiment with *Trhn*>*ACR1* flies (green dots) and relevant controls (*UAS-ACR1/+*, gray dots and *Trhn-Gal4*/+, black dots). Same graphical elements as above; see methods for details of ΔΔ analysis. Sample sizes as above. **H,** Time-course bubble plots and line plots of fed*R50H05*>*Chr* in a 2-h Espresso assay. Graphical elements as above. Sample sizes: N_Control light off_ = 120, N_Test light off_ = 60, N_Control light on_ = 140, N_Test light on_ = 70. **I**, Bubble plots and time lines of fed *R50H05*>*ACR1* flies and controls. Graphical elements as above. Sample sizes: N_Control light off_ = 40, N_Test light off_ = 20, N_Control light on_ = 77, N_Test light on_ = 38 **J,** Estimation graphic of the quantification of the optogenetic effect size of activating neurons in starved *R50H05*>*Chr* flies. See above and methods for full explanation of ΔΔ analysis. **K,** Quantification of the optogenetic effect size of inhibiting neurons in starved *R50H05*>*ACR1* flies. See above for graphical elements and sample sizes. **L**, Forest plot of volume effect sizes (ΔΔ values, in SD) in (left to right) activated *Trhn-Gal4* cells (starved), inhibited *Trhn-Gal4* cells (fed), activated *R50H05* cells (fed), and inhibited *R50H05* cells (starved). Units are nL of liquid. Color code for half-violin plots is Red = Chr activation; Green = ACR1 inhibition. The horizontal shaded ribbons show the confidence intervals of the feeding-volume effects of 24-h and 48-h starvation (pink and purple, respectively). **M,** Forest plot of ΔΔ values of changes in feeding duration (min) for four optogenetic experiments. The pink and purple ribbons indicate the effects of starvation on feeding duration. **N,** Forest plot of optogenetic ΔΔ changes in meal size (nL) in four experiments. Graphical elements as above. **O,** Forest plot of optogenetic ΔΔ changes in meal counts in four experiments. Graphical elements as above. All flies used were adult males (5–10 days post-eclosion). Optogenetic effects with controls separately plotted are shown in **Figure S2**.

**Figure S1.**
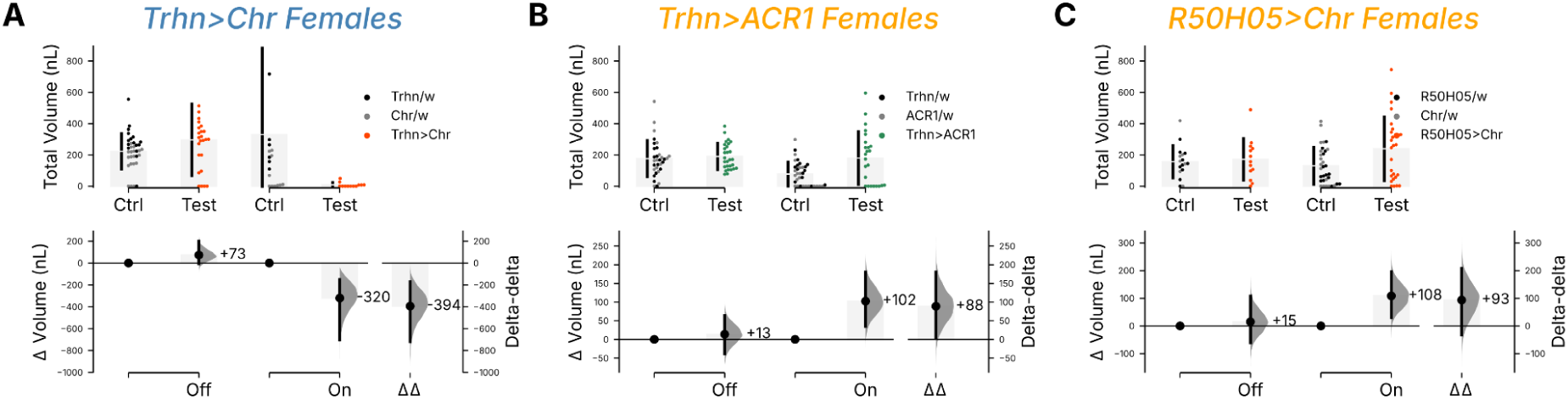
Optogenetic effects on food intake with serotonergic neurons in females. The effects of optogenetic manipulation of serotonergic neurons on feeding behavior in mated adult females (5–10 days post-eclosion), with three optogenetic interventions. In all panels, data are visualized using Cumming plots, which use swarm plots to show observed values (top) and contrast plots (bottom) to display the effect sizes. **A,** Volume effects in starved female flies with and without *Trhn>Chr* activation. Top panel displays total volume (nL) consumed in control (Ctrl) and Test groups. The bottom panel is a contrast plot with the Δ (left and center) and ΔΔ (right) effect sizes, indicating the changes in feeding volume. Numeric effect sizes are noted next to the half-violin curves. Sample sizes: N_Control light off_ = 35, N_Test light off_ = 25, N_Control light on_ = 18, N_Test light on_ = 12. **B,** Volume effects from neuronal inhibition in fed female controls and *Trhn>ACR1* flies. The ΔΔ effect size shows the overall optogenetic effect, with effect sizes annotating the half violins. Sample sizes: N_Control light off_ = 32, N_Test light off_ = 28, N_Control light on_ = 36, N_Test light on_ = 24. **C,** Volume effects in a *R50H05>Chr* activation experiment using fed females. Sample sizes: N_Control light off_ = 17, N_Test light off_ = 13, N_Control light on_ = 33, N_Test light on_ = 27.

**Figure S2.**
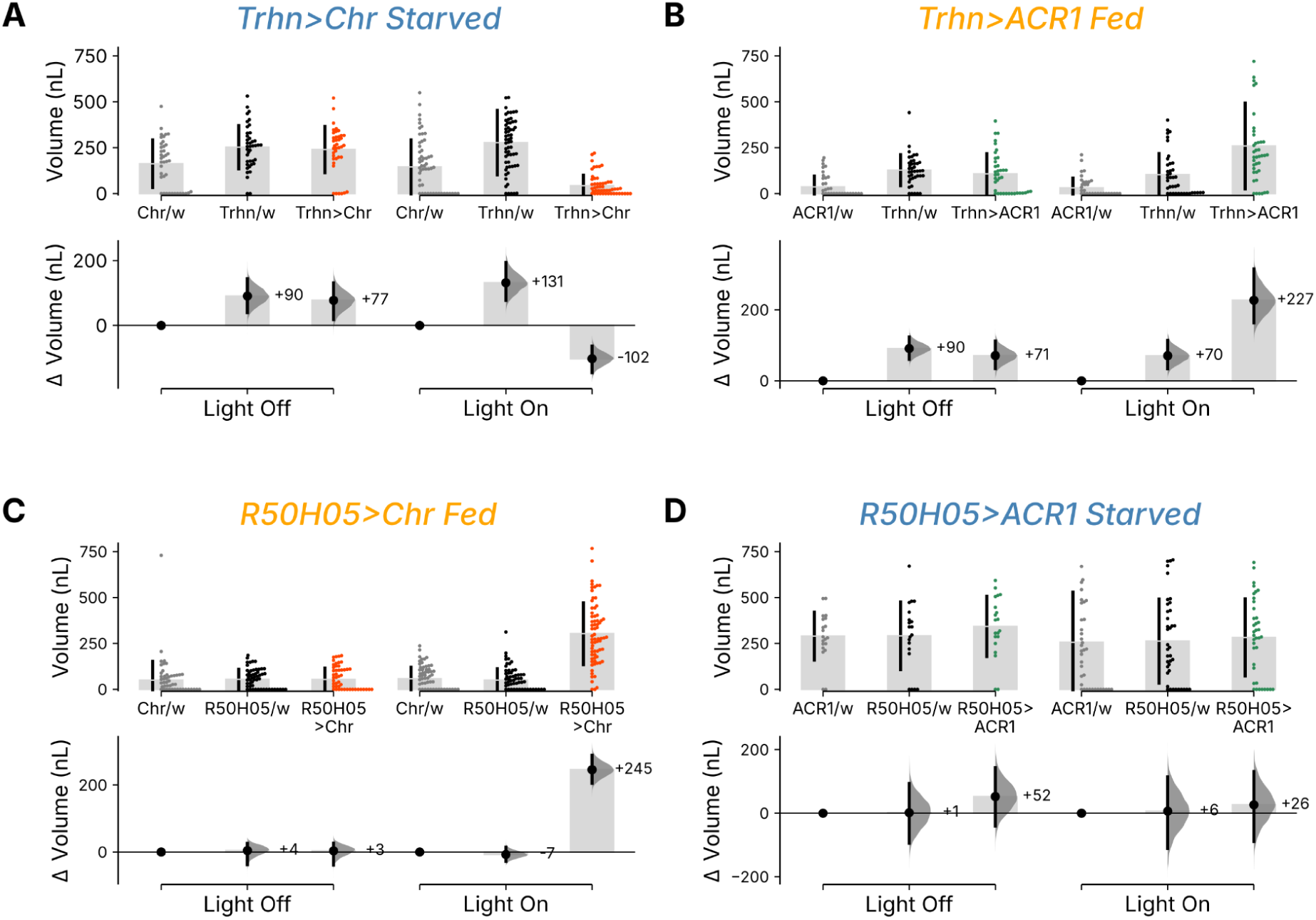
Optogenetic effects on main-figure experiments with separated genetic controls. Estimation plots showing the effects of optogenetic manipulations on feeding volume, with Driver Control and Responder Control displayed as separate columns to allow assessment of genetic background differences. Each panel includes a swarm plot showing individual measurements (top) and a contrast plot depicting the effect sizes (bottom). In the swarm plots, vertical bars represent the standard deviation, with the central gap indicating the mean value. In the lower axes (contrast plots), black dots denote mean differences, black bars represent the 95CI, and half-violins show the bootstrapped difference distributions. Numbers immediately adjacent to the half-violin curves are the mean effect sizes. This display is complementary to the pooled ΔΔ format shown in the main figures. **A,** Estimation plot of feed volume in starved *Trhn*>*Chr* flies with Driver and Responder Controls shown separately. **B,** Estimation plot of feed volume in fed *Trhn*>*ACR1* flies with separated controls. **C,** Estimation plot of feed volume in fed *R50H05*>*Chr* flies with separated controls. **D,** Estimation plot of feed volume in starved *R50H05*>*ACR1* flies with separated controls. Adult male flies (5–10 days post-eclosion) were used throughout.

### *Trhn-Gal4* and *R50H05* neurons regulate distinct locomotor patterns

The three main epochs of the feeding cycle—food seeking, eating, and satiety—are associated with distinct patterns of locomotion and quiescence. As natural feeding states involve coordinated changes in both feeding *per se* and movement, we next examined how optogenetic manipulations of serotonergic cells affected locomotor features. We reasoned that walking speed could both control for motor deficits and reflect foraging drive; elevation and latency could reflect motivation to feed; food port occupancy could indicate whether flies found or were interested in food; and peri-feed locomotion could indicate whether internal state was updated by nutrient intake.

Each Espresso chamber is vertically oriented, with the food port located at the top. As such, hungry flies will spend a lot of time at the top, near to or within the food port (**Figure 1M**). When the serotonergic neurons of hungry *Trhn>Chr* flies were activated, however, we saw that the flies dwelled at the chamber floor as indicated by reduced elevation (**Figure 3A, E**), suggesting either reduced food-seeking motivation or a locomotor deficit, although the flies had no problem finding food as indicated by their unaltered food port occupancy (**Figure 3F**). To exclude a motor impairment as an alternative explanation for the lower elevation, we analyzed walking speed and number of falls and found no gross deficit (**Figure 3H, S3E**). We do note that except for a brief dip at the beginning of the assay, walking speed was normal for the majority of the time and enhanced towards the later part of the assay (**S3A, B)**. This speed enhancement occurred while feeding was suppressed in both epochs (**Figure S3C, D**), although the suppression was weaker in the second epoch by 83 nL. By contrast, fed flies normally avoid the food port (**Figure 1 M, N**). When the serotonergic neurons of fed/sated *Trhn>ACR1* flies were inhibited, we saw increased time spent around the food port (**Figure 3B**) and thus a concomitant increase in elevation (**Figure 3E**) and food port occupancy (**Figure 3F**). Indeed, light-inhibited fed *Trhn>ACR1* flies moved more like hungry flies: relative to fed controls, they had a lower latency to first feed (**Figure 3G**), slower walking (**Figure 3H**), and a similar peri-feed speed ratio, i.e. postprandial slowing (**Figure 3I**). Interestingly, *Trhn* neurons had bidirectional control over five locomotion metrics, such that their inhibition mimicked hungry movement while their activation elicited the opposite effects.

We next performed optogenetic experiments to manipulate *R50H05* neuron activity. When fed *R50H05>Chr* flies were actuated, they showed reduced feeding latency, similar to hungry flies (**Figure 3G**), but also increased their dispersal in the chamber (**Figure 3C, S3F)** and walked much faster (**Figure 3G, H**). These latter effects contrast with naturally hungry flies, which also eat large volumes of food, but focus their activity around the food-port and walk a bit slower than sated flies in the presence of food (**Figure S3G**). In addition, the increase in locomotor speed is independent of starvation state and even in the absence of food in the port this increase was observed (**Figure S3I**). Although *R50H05>Chr* activation drives a large increase in food consumption (Δ = +247 nL; **Figure 2J, L**), the aberrant locomotor features imply that *R50H05* activity does not recreate natural hunger-like movement but instead promotes generalized locomotion and exploration. *R50H05>ACR1* inhibition had no discernible effect on any of the locomotor metrics (**Figure 3D, E-I**).

From these experiments, we conclude that these two serotonergic populations have fundamentally different effects on feeding-associated movement patterns. *Trhn-Gal4* neurons coordinate feeding-related locomotor patterns that are concordant with natural hunger and satiety states, while *R50H05* neurons promote locomotion that appears dissimilar to natural hunger.

**Figure 3.**
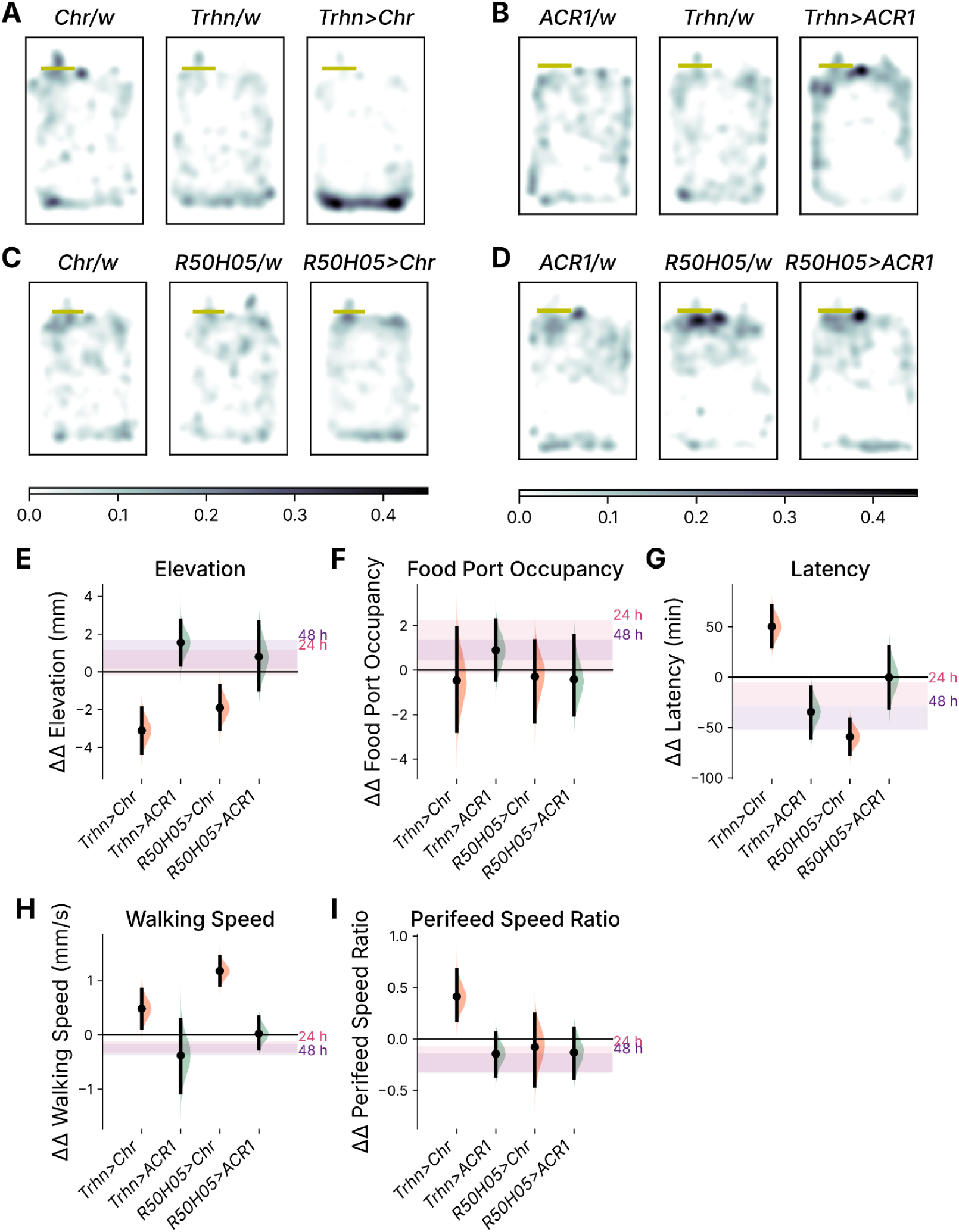
Effects of *Trhn*-*Gal4* and *R50H05* manipulations on locomotion. The impact of optogenetic manipulations on locomotor behaviors associated with feeding, using heatmaps to show spatial distribution and forest plots to summarize behavioral quantification. **A,** Heatmaps of mean Z-scores for time spent in chamber locations during illumination of starved flies, including *Trhn>Chr* flies and two control lines (*UAS-Chr/+*, and *Trhn-Gal4/*+). The heatmaps use a color gradient to indicate the density of fly presence. The heatmap color key is shown below panel **D**, with units in standard deviations. Yellow horizontal bars mark the spatial threshold for feed detection, i.e. the entrance to the feeding port. The white scale bar represents 2 mm. **B,** Heatmaps of mean Z-scores for time spent in chamber locations during illumination of fed flies in a *Trhn>ACR1* inhibition experiment including (*UAS-ACR1/+*, and *Trhn-Gal4/*+ controls. Heatmap scale, feed-detection threshold, and scale bar as above. **C,** Heatmaps of mean Z-scores for time spent in chamber locations during illumination of fed flies in a *R50H05>Chr* activation experiment including (*UAS-Chr/+*, and *R50H05/*+ controls. Heatmap scale, feed-detection threshold, and scale bar as above. **D,** Heatmaps of mean Z-scores for time spent in chamber locations during illumination of starved flies in a *R50H05>ACR1* inhibition experiment including (*UAS-Chr/+*, and *R50H05/*+ controls. Heatmap scale, feed-detection threshold, and scale bar as above. **E,** Forest plot of ΔΔ effect sizes for chamber elevations values from the quartet of optogenetic experiments. Each half-violin curve shows the distribution of bootstrapped ΔΔ differences in elevation; each black dot represents the mean value, each vertical bar indicates the 95CI. Pink and purple ribbons show the 95CIs of starvation effect sizes (24 h and 48-h starvation). Color code for half-violin plots: Red, Chr activation; Green, ACR1 inhibition. **F,** Forest plot of ΔΔ effect sizes for food port occupancy. Curves, dots, and bars indicate the ΔΔ distributions, means, and confidence intervals as above. The pink and purple ribbons show the confidence intervals of starvation latency effect-size intervals as above. **G,** Forest plot of ΔΔ effect sizes for latency to first feed. Curves, dots, and bars indicate the ΔΔ distributions, means, and confidence intervals as above. The pink and purple ribbons show the confidence intervals of starvation latency effect-size intervals as above. **H,** Forest plot of ΔΔ effect sizes for mean walking speed. Markers used as above. **I,** Forest plot of ΔΔ effect sizes for the peri-feed speed ratio, the ratio of walking speed (after-feed speed divided by before-feed speed). Markers used as above. Adult male flies (5–10 days post-eclosion) were used throughout.

**Figure S3.**
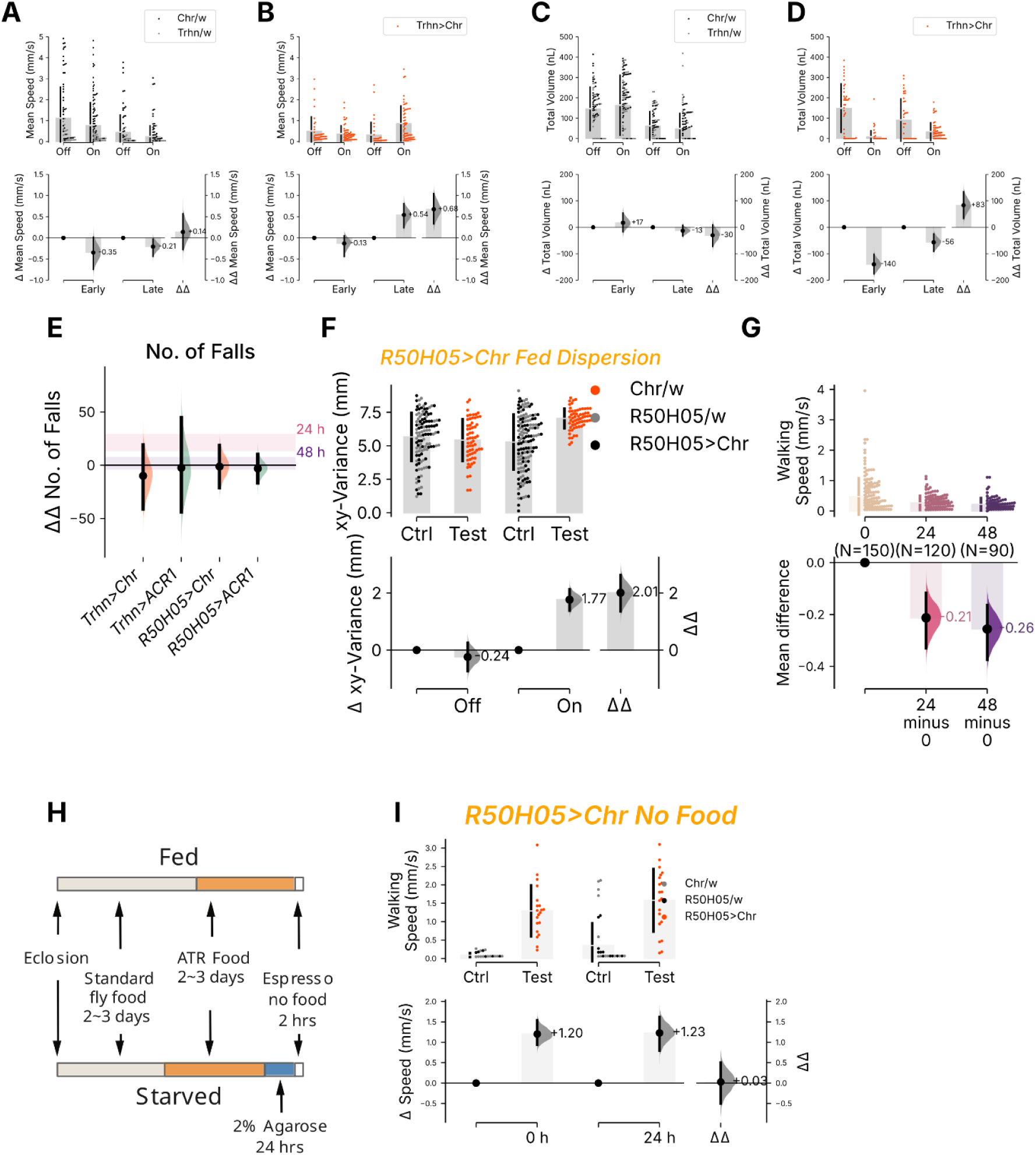
Additional optogenetic serotonergic effects on locomotion. **A**, The ΔΔ effect sizes of optogenetic activation on speed between early (first 60 min) and late (second 60 min) epochs for control animals. **B,** The ΔΔ effect sizes of optogenetic activation on speed between early and late epochs for *Trh>Chr* animals. **C,** The ΔΔ effect sizes of optogenetic activation on feed volume between early and late epochs for control animals. **D,** The ΔΔ effect sizes of optogenetic activation on feed volume between early and late epochs for *Trh>Chr* animals. **E,** The ΔΔ effect sizes of optogenetic activation and inhibition on the number of falls for different genotypes. The y-axis shows the changes in the number of falls (ΔΔ number of falls), comparing *Trhn>Chr*, *Trhn>ACR1*, *R50H05>Chr*, and *R50H05>ACR1*. Error bars indicate confidence intervals. Ribbons show the change in falls (Δ) for starved flies (24-h in pink, 48-h in purple). **F,** The effect of optogenetic activation of *R50H05>Chr* flies on their dispersion within feeding chambers. The top axis displays the location (xy) variance values for control (Ctrl) and experimental (Test) groups, with individual data points and distributions (standard deviation) shown. The bottom graph presents the change in dispersion (Δ variance in units) under light-off and light-on conditions, with ΔΔ showing the 2-way overall difference. Sample sizes: N_Control light off_ = 120, N_Test light off_ = 60, N_Control light on_ = 140, N_Test light on_ = 70. **G,** Walking speed of *w^1118^* flies under fed and starved conditions. The top y-axis shows speed in mm/s, while the lower y-axis indicates the mean difference in speed. Data compares walking speeds in fed flies (0-h starvation) and after 24- and 48-h of starvation. Sample sizes (N) are provided for each starvation regime. Sample sizes: N_0h_ = 150, N_24h_ = 120, N_48h_ = 90. **H,** Schematic outlining experimental regimes for studying the effects of starvation and optogenetic manipulation on fly behavior. Flies were first raised on standard fly food, before transfer to all-*trans* retinal food; if starved, they were placed onto nutrient-free 2% agarose for 24 h, prior to testing for 2 h in Espresso. **I,** Effects of optogenetic Chr activation of *R50H05* neurons on the walking speed of fed and starved flies. The top graph shows speed (mm/s) for control and test groups under fed (0 h) and starved (24 h) conditions. The bottom graph presents the changes in speed (Δ Speed in mm/s) and the difference between these changes (ΔΔ), illustrating the impact of starvation on optogenetic effects on walking. Sample sizes: N_Control light off_ = 40, N_Test light off_ = 20, N_Control light on_ = 39, N_Test light on_ = 20. Adult male flies (5–10 days post-eclosion) were used throughout.

### Subtractive genetics localizes serotonergic satiation to the *Trhn-Gal4* NOT *R50H05* subset

Despite their considerable expression overlap, given that the *Trhn-Gal4* and *R50H05* drivers elicit strikingly different effects on food consumption and its associated behaviors, we assumed that these drivers capture different neurons with different functions (*20*, *21*, *35*). We hypothesized that a subset of neurons captured by *Trhn-Gal4*, but not *R50H05*, was responsible for the naturalistic satiety-like activation effects. To investigate this hypothesis, we refined our serotonergic targeting approach by using a genetic NOT operation, using *Gal80* repression of driver *Gal4* activity and therefore restricting the transcription of the responder gene to cells that lack the Gal80 repressor. First, we subtracted the *R50H05* pattern from *Trhn-Gal4* expression. In starved *Trhn*-NOT-*R50H05>Chr* flies (**Figure S4A**), activation resulted in reduced consumption (Δvolume = –120 nL), demonstrating that the non-intersecting neurons are not required for optogenetic feeding suppression (**Figure 4, Figure S5A**). We then subtracted the *Trhn* pattern from *R50H05* expression. In sated *R50H05*-NOT-*Trhn>Chr* flies (**Figure S4B**), activation elicited only a minor increase in consumption (Δvolume = +25 nL; **Figure 4, Figure S5B**), indicating that the neural activity responsible for the large *R50H05* consumption increase occurs in neurons captured by both drivers. The neurons responsible for the feeding suppression seen upon *Trhn>Chr* activation are thus located in a *Trhn*-subset that is not captured by *R50H05*, while the feeding increase from *R50H05>Chr* activation is mediated by cells located within the *Trhn* ∩ *R50H05* intersection.

### Ventral-cord serotonergic neurons instruct feeding suppression

These intersectional analyses pointed to a specific anatomical locus for serotonergic feeding control. One conspicuous group of *Trhn*-NOT-*R50H05* neurons are 5-HT^+^ cells in the VNC. This led us to investigate whether VNC neurons mediate the observed feeding phenotypes. We hypothesized that these VNC-serotonin neurons mediated the *Trhn>Chr* phenotype. To test this, we used another NOT operator driven by the *teashirt* enhancer (*tsh,* (*36*)) that expresses the Gal80 repressor in the VNC (*37*), generating a *Trhn*-NOT-*tsh* expression pattern, thereby expressing Chr almost exclusively in *Trhn-Gal4* brain cells (a *Trhn*-Brain pattern; **Figure S4C**). With nearly all VNC expression repressed, *Trhn*-NOT-*tsh>Chr* activation in starved flies had a negligible effect on suppressing food consumption (Δvolume = –36 nL; **Figure 4, Figure S5C**). We interpret this cautiously: tsh-Gal80 repression of the abdominal ganglion is incomplete, and within-group variability suggests VNC repression differs across individuals (Figure S4C). We therefore treat –36 nL as an upper bound on the brain contribution.

We then established a reciprocal experiment in order to express only in the VNC, using the *tsh* to restrict expression. To do so, we used a quaternary genetic scheme (α*Tub84B-frt-Gal80-stop-frt; tsh-LexA; LexAop2-FLPL*), which allowed us to express the Gal80 repressor everywhere except the VNC (*38*). This genetic AND operator allowed Chr expression only in *Trhn-Gal4* cells in the *tsh^+^*VNC (**Figure S4D**). Consequently, activation in starved *Trhn*-AND-*tsh>Chr* flies drove substantial feeding suppression (Δvolume = –227 nL; **Figure 4, Figure S5D**) that was comparable with the *Trhn>Chr* effect size (Δvolume = –200 nL; **Figure 2F**). These findings suggest that the neurons responsible for *Trhn>Chr* feeding suppression are located in the VNC.

Next, we explored the serotonergic VNC cells responsible for suppressing food consumption by developing genetic tools that could capture 5-HT+ VNC neurons. An *in silico* search of a single-cell RNA data from the Drosophila VNC (*39*) identified candidate genes whose enhancers seemed suitable for generating split-Gal4 lines (*40*) for narrow neuronal targeting. An anatomical screen identified two split-*Gal4* lines that showed selective expression in the VNC: serotonin × VNC 1 and 2 (SXVNC1 and SXVNC2; **Figure S4E, F**). SXVNC1 expresses Gal4 in thoracic 5-HT cells, whereas SXVNC2 appears to express Gal4 in a cluster of neurons in the tip of the VNC. SXVNC1*>Chr* and SXVNC2*>Chr* activation in hungry flies led to reduced food consumption (Δvolume = –250 nL, –181 nL; **Figure 4, S5 G, H**). These results confirm that the *Trhn-*VNC neurons are responsible for essentially all the feeding suppression observed in starved, activated *Trhn-Gal4*>*Chr* flies.

Having shown that serotonergic VNC neurons can potently suppress feeding when activated, we next asked whether they are required for natural satiety. Inhibition of each VNC-specific serotonergic driver in sated flies produced modest increases in consumption (Δvolume = +33 nL, +50 nL, and +35 nL for Trhn-AND-tsh, SXVNC1, and SXVNC2, respectively; **Figure 4, S5 E, F, I, J**). The effect sizes are small and the confidence intervals include negative values, so do not individually constitute strong evidence for each population’s contribution. Nonetheless, the three mean differences are increases and directionally concordant with the large effect seen upon broad *Trhn>ACR1* silencing (+165 nL; **Figure 2G**). We cautiously interpret this as a partial, distributed contribution of VNC serotonergic neurons to satiety maintenance, with the total VNC contribution likely in the range of +30–50 nL—roughly a quarter of the broad-driver effect. However, these estimates’ precision ranges are also consistent with each VNC subset making negligible contributions, i.e. are not required for satiety maintenance.

**Figure 4.**
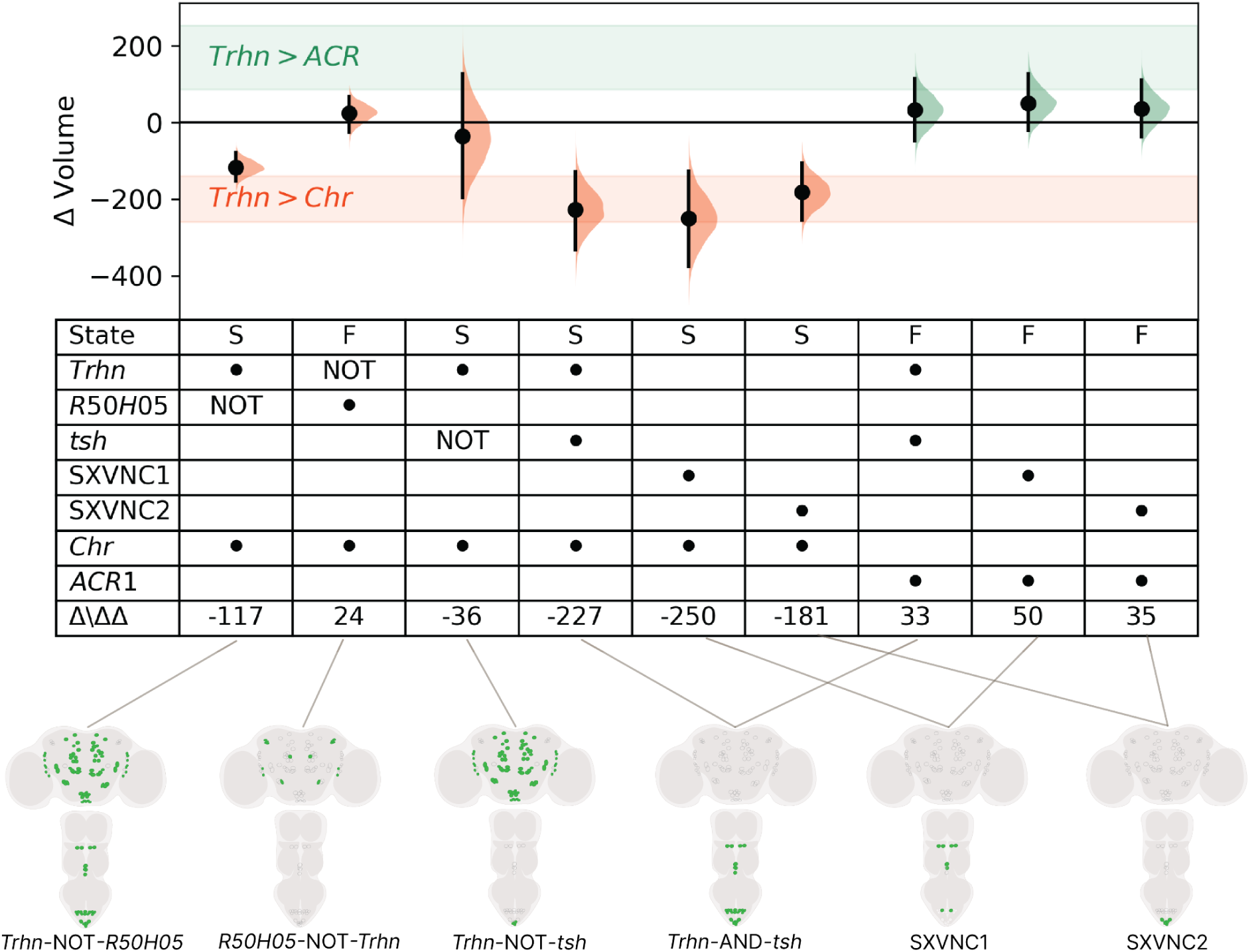
Activity in the *Trhn* VNC neurons suppresses feeding. A forest plot illustrating the optogenetic effects on feeding volume in starved and fed flies, using various Boolean composite driver lines. The figure shows how specific manipulations of serotonergic groups in the brain and VNC influence feeding behavior under different physiological states. **Top Panel.** The y-axis represents the change in feed volume (Δ Volume), with positive values indicating an increase and negative values indicating a decrease in feeding volume. Each half-violin curve shows the distribution of each effect size, with the black dot denoting the mean effect, and the black vertical bar representing the 95CI. The interventions include optogenetic activation (*Chr*, red) and inhibition (*ACR*, green). The shaded regions highlight the CI of effects of the broad driver, *Trhn>Chr* (red) and *Trhn>ACR* (green). **Grid key.** Below the plot, a table key summarizes the experimental conditions and outcome: nutritional state (S for starved, F for fed), transgenes in each line, and the corresponding ΔΔ effect sizes. For AND drivers, two black dots mark the two constituent drivers; for NOT lines, the repressing line is marked ‘NOT’. **Schematic.** The schematic provides representations of the anatomical expression patterns for each line used. Green indicates serotonergic cells expressing Gal4, while gray indicates serotonergic cells not expressing Gal4. Adult male flies (5–10 days post-eclosion) were used throughout.

**Figure S4.**
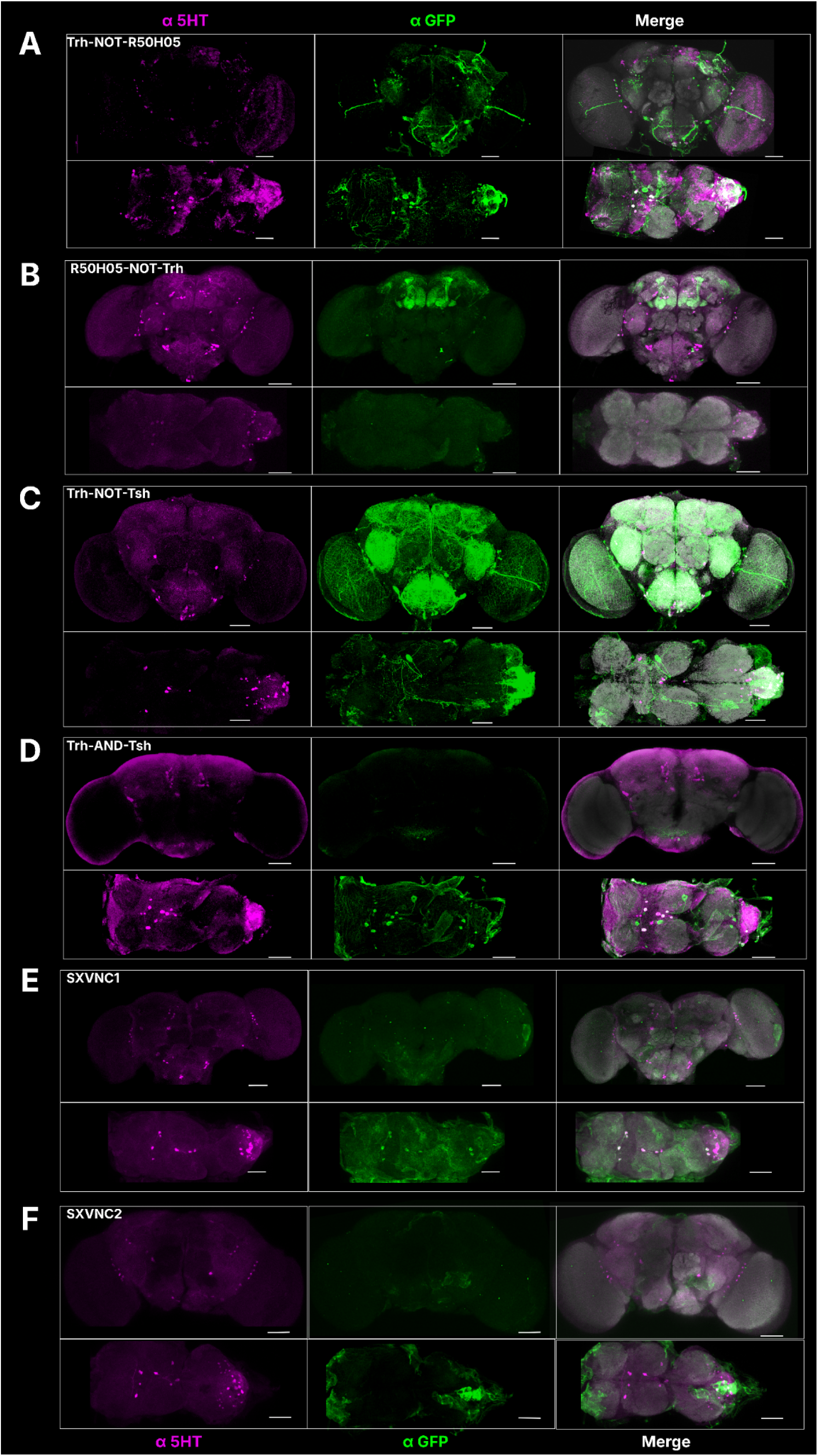
Expression patterns of various serotonin neuronal driver lines. Maximum intensity projections of confocal images illustrating the expression patterns of various intersectional driver lines targeting serotonin neurons. Each panel (**A** through **F**) is a representative staining of a specific Boolean transgenic line, showing 5-HT^+^ cells in the brain and ventral nerve cord (VNC). In all panels, blue indicates stain from the nc82 antibody for neuropil (α-BRP); magenta is α-5-HT signal; and green is α-GFP signal from binding to yellow fluorescent protein (YFP) that is fused to Chr or ACR1. Scale bars = 50 µm. Refer to Table S1 for all complete genotypes. **A.** *Trhn*-NOT-*R50H05*>*Chr*. Green Chr^+^ cells are present in both the brain and VNC, with a slightly higher number in the brain, indicating a broad expression pattern. **B.** *R50H05*-NOT-*Trhn*>*Chr*. Green Chr^+^ cells are observed in the brain but not in the VNC; prominent mushroom body expression is visible, consistent with R50H05 expression in the mushroom bodies shown in Figure 2B, and reflecting the R50H05 population that does not overlap with Trhn-LexA-expressing cells. **C.** *Trhn*-NOT-*tsh*>*Chr*. Green Chr^+^ cells are predominantly in the brain, with fewer in the VNC, indicating a stronger expression in the brain. Transgenes are *Trhn-Gal4 > tsh-Gal80; UAS-Chr*. **D.** *Trhn*-AND-*tsh*>*Chr*. Green Chr⁺ cells are more concentrated in the VNC compared to the brain, suggesting focused VNC expression. **E.** *SXVNC1>ACR1.* Green ACR1⁺ cells are exclusively located in the VNC, with no visible expression in the brain, indicating VNC-specific expression. Yellow arrowheads indicate cells expressing ACR1. **F.** *SXVNC2*>*ACR1*. Green ACR1⁺ cells are found only in the VNC, with no expression in the brain, suggesting targeted VNC expression. Adult male flies (5–10 days post-eclosion) were used throughout. Ectopic labelling is visible in some panels although the nature of the cells is uncertain.

**Figure S5.**
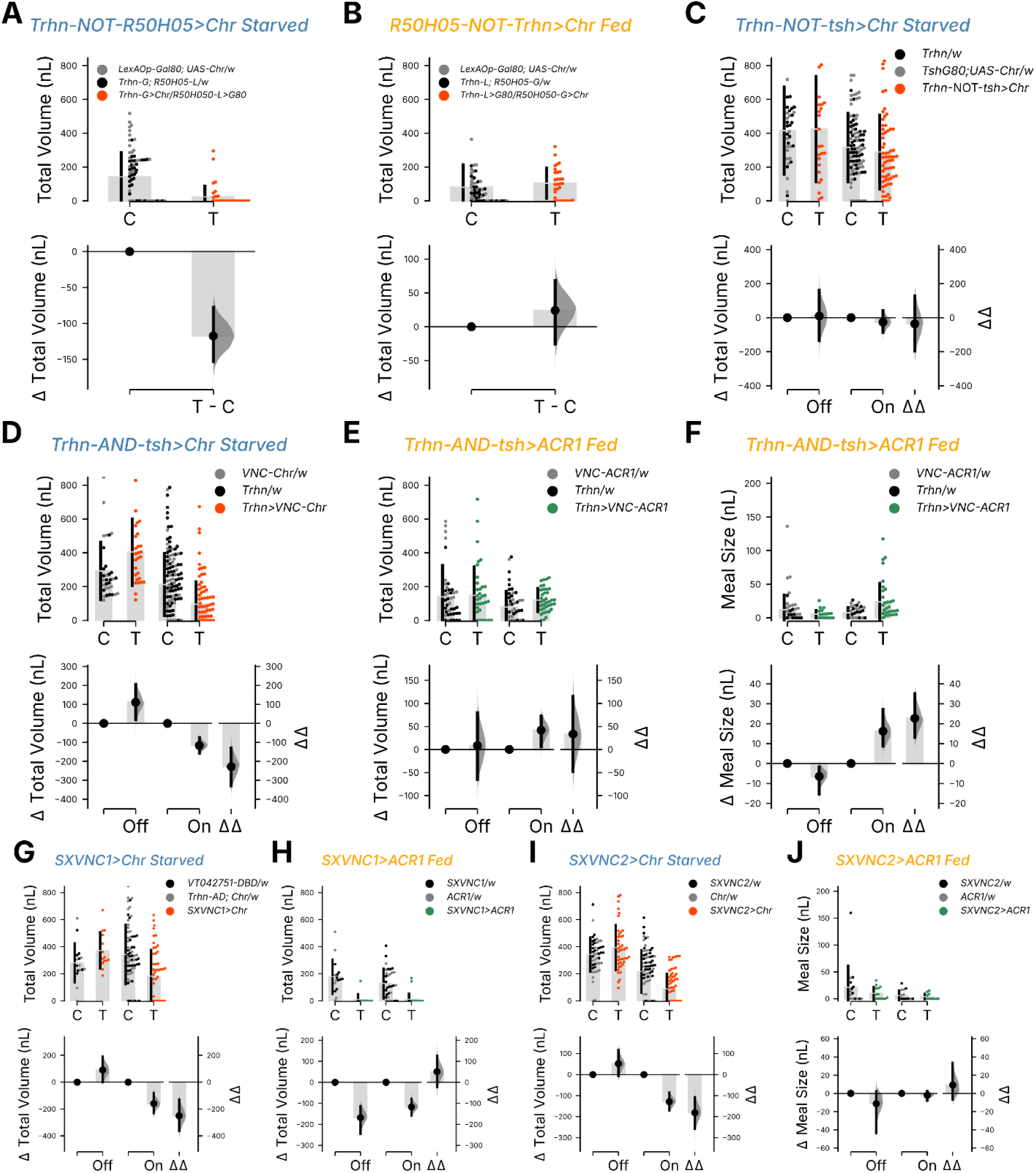
Effects of optogenetic manipulations on feeding intake. Estimation plots showing the effects of optogenetic manipulations on feeding intake. Each panel includes a swarm plot showing individual measurements (top) and a contrast plot depicting the effect sizes (bottom). In the swarm plots, vertical bars represent the standard deviation, with the central gap indicating the mean value. In the lower axes (contrast plots), black dots denote mean differences, black bars represent the 95CI, and half-violins show the bootstrapped distribution differences. Numbers immediately adjacent to the half-violins are the mean effect sizes. **A,** A Cumming estimation plot showing the effect on feed volume when the *Trhn-NOT-R50H05>Chr* population was activated by red light compared to control flies. The plot includes observed values (top) and a contrast plot (bottom). Sample sizes: N_Control light on_ = 80, N_Test light on_ = 40. **B,** A Cumming estimation plot of the effect on feed volume when the *R50H05-NOT-Trhn>Chr* population was activated by red light compared to control flies. Sample sizes: N_Control light on_ = 60, N_Test light on_ = 30. **C,** A ΔΔ plot showing the effects of activating brain serotonin neurons (*Trhn-NOT-tsh>Chr*) on feed volume in starved flies. The plot includes a swarm plot of individual measurements and a contrast plot for effect size. Sample sizes: N_Control light off_ = 35, N_Test light off_ = 25, N_Control light on_ = 113, N_Test light on_ = 67. **D,** A ΔΔ plot depicting the effects of activating ventral nerve cord (VNC) serotonin neurons (*Trhn-AND-tsh>Chr*) on feed volume in starved flies. The plot follows the same format as panel **C**, with individual measurements and a contrast plot. Sample sizes: N_Control light off_ = 32, N_Test light off_ = 28, N_Control light on_ = 139, N_Test light on_ = 88. **E,** A ΔΔ plot showing the effects of inhibiting VNC serotonin neurons with *ACR1* (*Trhn-AND-tsh>ACR1*) on feed volume in fed flies. The plot includes a swarm plot of individual measurements and a contrast plot for effect size. Sample sizes: N_Control light off_ = 51, N_Test light off_ = 38, N_Control light on_ = 52, N_Test light on_ = 38. **F,** A ΔΔ plot of meal size illustrating the effects of inhibiting VNC serotonergic neurons in fed flies with *ACR1* (*Trhn-AND-tsh>ACR1*). The plot follows the same format as panel **E**, with individual measurements and a contrast plot. Sample sizes: N_Control light off_ = 51, N_Test light off_ = 38, N_Control light on_ = 52, N_Test light on_ = 38. **G,** A ΔΔ plot of meal size illustrating the effects of activating SXVNC1 neurons in fed flies with *Chr* (*SXVNC1>Chr*). The plot follows the same format as panel **E**, with individual measurements and a contrast plot. Sample sizes: N_Control light off_ = 16, N_Test light off_ = 14, N_Control light on_ = 68, N_Test light on_ = 52. **H,** A ΔΔ plot of meal size illustrating the effects of inhibiting SXVNC1 neurons in fed flies with *ACR1* (*SXVNC1>ACR1*). The plot follows the same format as panel **E**, with individual measurements and a contrast plot. Sample sizes: N_Control light off_ = 16, N_Test light off_ = 14, N_Control light on_ = 32, N_Test light on_ = 28. **I,** A ΔΔ plot of meal size illustrating the effects of activating SXVNC2 neurons in fed flies with *Chr* (*SXVNC2>Chr*). The plot follows the same format as panel **E**, with individual measurements and a ^contrast plot. Sample sizes:^ N_Control light off_ = 48, N_Test light off_ = 42, N_Control light on_ = 82, N_Test light on_ = 68. **J,** A ΔΔ plot of meal size illustrating the effects of inhibiting SXVNC2 neurons in fed flies with *ACR1* (*SXVNC2>ACR1*). The plot follows the same format as panel **E**, with individual measurements and a contrast plot. Sample sizes: N_Control light off_ = 16, N_Test light off_ = 14, N_Control light on_ = 16, N_Test light on_ = 14. Adult male flies (5–10 days post-eclosion) were used throughout. Optogenetic effects with controls separately plotted are shown in **Figure S6**.

**Figure S6.**
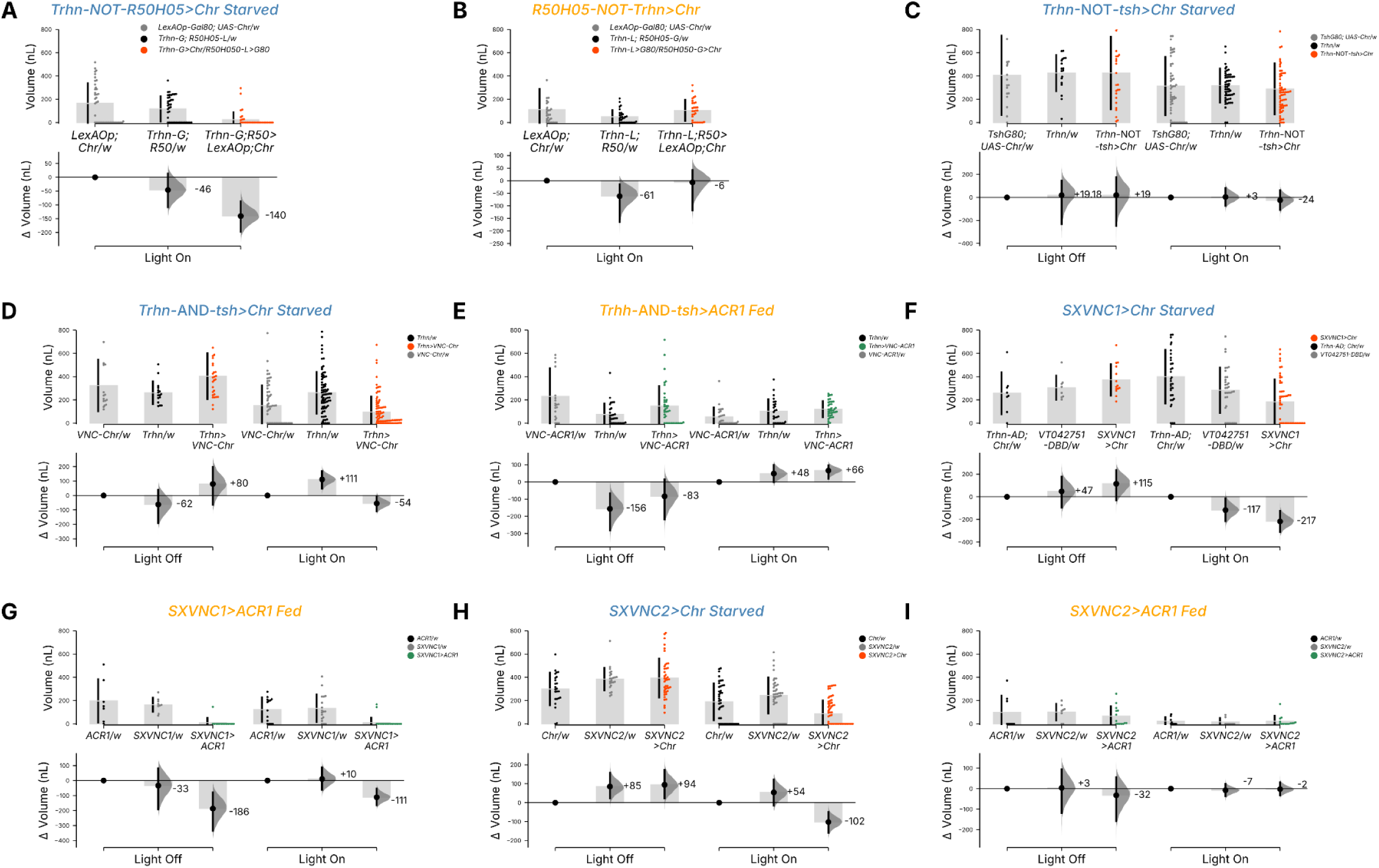
Optogenetic effects on main-figure experiments with separated genetic controls. Estimation plots showing the effects of optogenetic manipulations on feeding volume, with Driver Control and Responder Control displayed as separate columns to allow assessment of genetic background differences. Each panel includes a swarm plot showing individual measurements (top) and a contrast plot depicting the effect sizes (bottom). In the swarm plots, vertical bars represent the standard deviation, with the central gap indicating the mean value. In the lower axes (contrast plots), black dots denote mean differences, black bars represent the 95CI, and half-violins show the bootstrapped difference distributions. Numbers immediately adjacent to the half-violin curves are the mean effect sizes. This separated display allows direct comparison of the two genetic backgrounds and is complementary to the pooled ΔΔ format shown in the main figures. **A,** Estimation plot of feed volume in starved *Trhn-*NOT*-R50H05*>*Chr* flies with Driver and Responder Controls shown separately. **B,** Estimation plot of feed volume in fed *R50H05-*NOT*-Trhn*>*Chr* flies with separated controls. **C,** Estimation plot of feed volume in starved *Trhn-*NOT-*tsh*>*Chr* flies with separated controls. **D,** Estimation plot of feed volume in starved *Trhn-*AND-*tsh*>*Chr* flies with separated controls. **E,** Estimation plot of feed volume in fed *Trhn-*AND-*tsh*>*ACR1* flies with separated controls. **F,** Estimation plot of feed volume in starved *SXVNC1*>*Chr* flies with separated controls. **G,** Estimation plot of feed volume in fed *SXVNC1*>*ACR1* flies with separated controls. **H,** Estimation plot of feed volume in starved *SXVNC2*>*Chr* flies with separated controls. **I,** Estimation plot of feed volume in fed *SXVNC2*>*ACR1* flies with separated controls. Adult male flies (5–10 days post-eclosion) were used throughout.

### Feeding-related behavioral features are largely independent

Having established that Espresso reliably captures starvation-induced changes in food intake (**Figure 1**) and that Trhn-Gal4 neurons bidirectionally modulate consumption (**Figures 2–4**), we next asked whether the additional behavioral dimensions tracked by the system such as locomotion, port occupancy, and feeding microstructure provided information beyond consumption alone. Consumption is a reliable and sensitive measure of hunger drive, but a manipulation that changes consumption could reflect not only satiety signaling *per se*, but also altered upstream functions such as motor function, motivation, all of which are indistinguishable from a consumption-only read-out (*13*, *28*). Our analysis therefore proceeded in two stages: first, we evaluated optogenetic interventions against consumption metrics (**Figures 2–4**); second, we built a multi-metric analytical framework, using phenotypic vectors (‘phenovectors’), to determine how well the overall behavioral profile of each manipulation aligned with natural state transitions. From the food capillary-imaging and fly-tracking data, we calculated 17 summary metrics (**Methods**) that describe different aspects of foraging, consumption, and locomotion, to comprehensively characterize feeding-related behaviors. To examine the extent of redundancy across these 17 metrics, we inspected the intra-individual correlogram for all nutrition states. This analysis showed highly diverse correlation values (mean *r* = 0.25, standard deviation = 0.22, **Figure S7A**), indicating that most of the metrics are poor approximations of each other (**Figure S7 B-G**). Correlograms of feeding states similarly showed low average inter-metric correlations for fed, and 24-h- and 48-h starved flies (*r =* 0.35, 0.26, 0.29, respectively, **Figure S7 H - K**). The weaker correlograms in starved flies indicate that behavioral features dissociate across states (**Figure S7I**). In a complementary analysis, simple linear dimension reduction with principal component analysis failed to describe >90% of the metrics with fewer than ten principal components, further demonstrating that the multiple metrics were in fact mostly not redundant (**Figure S7L**). While metrics were largely non-redundant overall, the correlogram revealed two broadly correlated clusters corresponding to feeding microstructure and locomotion (**Figure S7A**). These results indicate that our behavioral metrics capture informative aspects of feeding-related behavior, validating their use as independent dimensions for understanding hunger–satiety states.

### Starvation effects can be summarized as a state-transition effect-size vector

Having established that the metrics capture distinct aspects, we next sought to use them to comprehensively characterize how hunger states manifest across these multiple dimensions. As hunger and satiety are continuous states rather than discrete categories, we therefore characterized the nutritional axis as a quantitative spectrum set by controlled starvation duration. We computed state-transition vectors between three reproducible positions along this axis to concisely summarise how the behavioral system shifts gradually toward hunger. Our experimental design is between-subject: each group of flies was assigned a defined position on the nutritional axis at experiment onset, allowing us to establish controlled, reproducible states.

Our early Espresso analyses showed that starved flies showed substantial consumption increases during refeeding (Δ = +86 nl and +270 nl, for 24- and 48-h, respectively; **Figure 1H**). Though consumption is a critical measure of hunger, we aimed to analyze all changes in this complex behavior, and to describe starvation-induced behavior changes with an integrated, ethomic approach (*33*). As the 17 features (and their differences) were necessarily measured on different scales, we standardized the mean differences (Δ values) between sated and hungry (24-h and 48-h starvation) flies by transformation into Hedges’ *g* (*41*) and used these effect sizes to represent a *state-transition vector* (**Figure 5A**). The starvation vectors represent substantial overall displacements; for example, for all behavioral features, the 48-h flies underwent a mean absolute change of |*g*| = 0.76 SD. Along with large increases in consumption, flies starved for 24-or 48-hours showed longer feeding duration, larger meal sizes, and faster feeding, but, interestingly, did not increase their feed count (meal frequency) (**Figure 5A**). The list of effect sizes of the differences in behavioral features between fed and starved animals, the fed→starved state-transition vector, is a concise summary of starvation’s effects; it can serve as a reference to quantitatively evaluate how closely artificial neural manipulations align with naturalistic hunger effects.

### Contextualized screen reveals *Trhn-Gal4* neurons as holistic regulators of hunger–satiety

Having established a quantitative framework for characterizing the natural hunger–satiety axis, we next sought to identify circuits that regulate changes similar to authentic state transitions. Our earlier findings that the *Trhn-Gal4* neurons could bidirectionally modulate food intake made them candidate satiety regulators. We hypothesized that if these neurons genuinely control state, rather than just affecting isolated feeding behaviors, their manipulation should recreate the coordinated behavioral changes we observed during natural hunger–satiety transitions. Our analysis framework comprised four steps. In step 1, we calculated vectors of standardized effect sizes to describe the holistic phenotypic profile of each pleiotropic intervention, referred to as *phenotypic effect vectors* (hereafter ‘phenovectors’). During step 2, we benchmarked the optogenetic phenovectors against the naturalistic state-transition vectors derived from the starvation experiments (**Figure 5A - C, Figure S8 C-H**). Along with the satiety→hunger vectors described above e.g. [24 h minus 0 h], we also computed the reverse vectors, e.g. [0 h minus 24 h], to represent hunger→satiety transitions. In step 3, we compared the serotonergic effects with other candidate feeding-regulation circuits by performing activating and inhibiting optogenetic analyses of three circuits known to be involved in feeding behavior and metabolism in adult flies: *NPF-Gal4* derived from the *neuropeptide F* gene shown to promote feeding and wakefulness (*42*, *43*); *Akh-Gal4* derived from *Adipokinetic hormone* shown to suppress feeding (*44*); and *Ilp2-Gal4* derived from *Insulin-like peptide 2* shown to promote feeding (*45*). Finally, in step 4, we used hierarchical clustering and regression to assess how closely each manipulation’s phenovector matched those of the satiety⇄hunger state transitions (**Figure 5D**).

The hierarchical clustermap showed that the behavioral features were grouped into three types: food-seeking, food consumption, and locomotion (**Figure 5D**). The experimental dimension segmented into three clusters: (1) one that contained the 24-h and 48-h hunger→satiety state transitions with the optogenetic phenovectors of fed-fly experiments; (2) an intermediate cluster with little similarity to state transitions; and (3) a cluster comprising the two satiety→hunger transition vectors and the phenovectors of starved optogenetic flies. Strikingly, *Trhn>Chr* and *Trhn>ACR1* clustered the nearest to the four natural state transitions (Clusters 1 and 3, respectively), suggesting that, among the lines tested, the broad *Trhn* network’s activity most faithfully controls hunger–satiety transitions. In Cluster 3, *Trhn>ACR1* inhibition was the nearest neighbor of the satiety→hunger state-transition vector derived from the 24-h and 48-h starvation experiments (**Figure 5D**). In Cluster 1, the nearest neighbors of the ‘reverse-starvation’ hunger→satiety vectors were *Akh>Chr*, *Trhn>Chr*, and *Trhn-*NOT*-R50H05>Chr* activation (**Figure 5D**). The rest of Cluster 1 comprised phenovectors for VNC-specific serotonergic driver activation. Cluster 2 comprised *R50H05>Chr*; and *Trhn-NOT-tsh>Chr* (both capturing serotonergic neurons in the brain) and other interventions with effects profiles that were distinct from the natural metabolic transition vectors. Interventions in this cluster induced disjointed phenovectors; for example, animals ate more while also showing heightened food-seeking, reflected in elevated walking speed and increased feed counts. (**Figure 5D**).

To quantify the similarity between the optogenetic interventions and a natural transition, we applied linear regression to each phenovector compared with 48-h starvation. When we ranked the experiments by the slope and correlation of their phenovectors against starvation, *Trhn* manipulations consistently emerged as close matches to natural hunger–satiety transitions. These measures showed that both *Trhn>Chr* (β = –0.83, *r* = –0.86) and *Trhn>ACR1* (β = –0.82, *r* = –0.86) were among the most similar to 48-h starvation, confirming that these manipulations largely recapitulate natural hunger ⇄ satiety shifts (**Figure S8 A, B, C, I, J**). Several VNC serotonergic neuron drivers also showed partial recapitulation of natural hunger ⇄ satiety shifts, including *SXVNC1>Chr* (β = –0.52, *r* = –0.76) and *SXVNC2*>*ACR1* (β = –0.38, *r* = –0.59). These results confirm that *Trhn-Gal4* cells, particularly in the VNC, are regulators of AHLS as assessed by the seventeen feeding and locomotor metrics captured in the Espresso framework. We note that this characterisation is necessarily scoped to the metrics collected here; the possibility remains that Trhn neuronal manipulations could dissociate from natural hunger or satiety along behavioral dimensions not measured by Espresso (such as olfactory responses, proboscis kinematics, or social interactions). Within the Espresso metric space, however, no other circuit we tested matched the breadth and fidelity of *Trhn* state recapitulation. With the possible exception of *Akh* neurons (**Figure S8 A, B, F, I, J)**, *Trhn-Gal4* cell activation and inhibition recapitulate the complex behavioral patterns of natural satiety and hunger more faithfully than other feeding-related neuromodulatory circuits. The strong alignment between activation of Akh, an independently shown satiety signal (*44*), with hunger -> satiety transition, provides a reassuring second DESTRA phenocopy. In contrast, *R50H05>Chr* neurons did not fully capture the starved state (**Figure S8 A, B, D, K)**, though were highly correlated with *NPF>Chr* flies (β = –0.73, r = –0.81, **Figure S8 L**), the meaning of which remains to be investigated. Increased walking speed and chamber dispersal in *R50H05*>*Chr* flies could in principle reflect enhanced foraging drive; however, this profile differs measurably from both naturally starved *w^1118^* flies and *Trhn*>*ACR1* flies within Espresso, where hunger-driven locomotion is directed toward and concentrated around the food port rather than generalised across the chamber. Lastly, we examined whether our findings were sensitive to analytical choices in the phenovector framework. To assess the contribution of ratio metrics, we recomputed rankings excluding all ratio metrics; the relative ordering of manipulations shifted **(Figure S9A)**, indicating that ratio metrics capture biologically distinct information not redundant with the raw metrics alone. To assess whether our findings were sensitive to the choice of similarity metric, we recomputed the clustermap using Spearman’s rank correlation **(Figure S9B)** and cosine distance **(Figure S9C)**. In both cases, the clustering structure was mostly similar.

**Figure 5.**
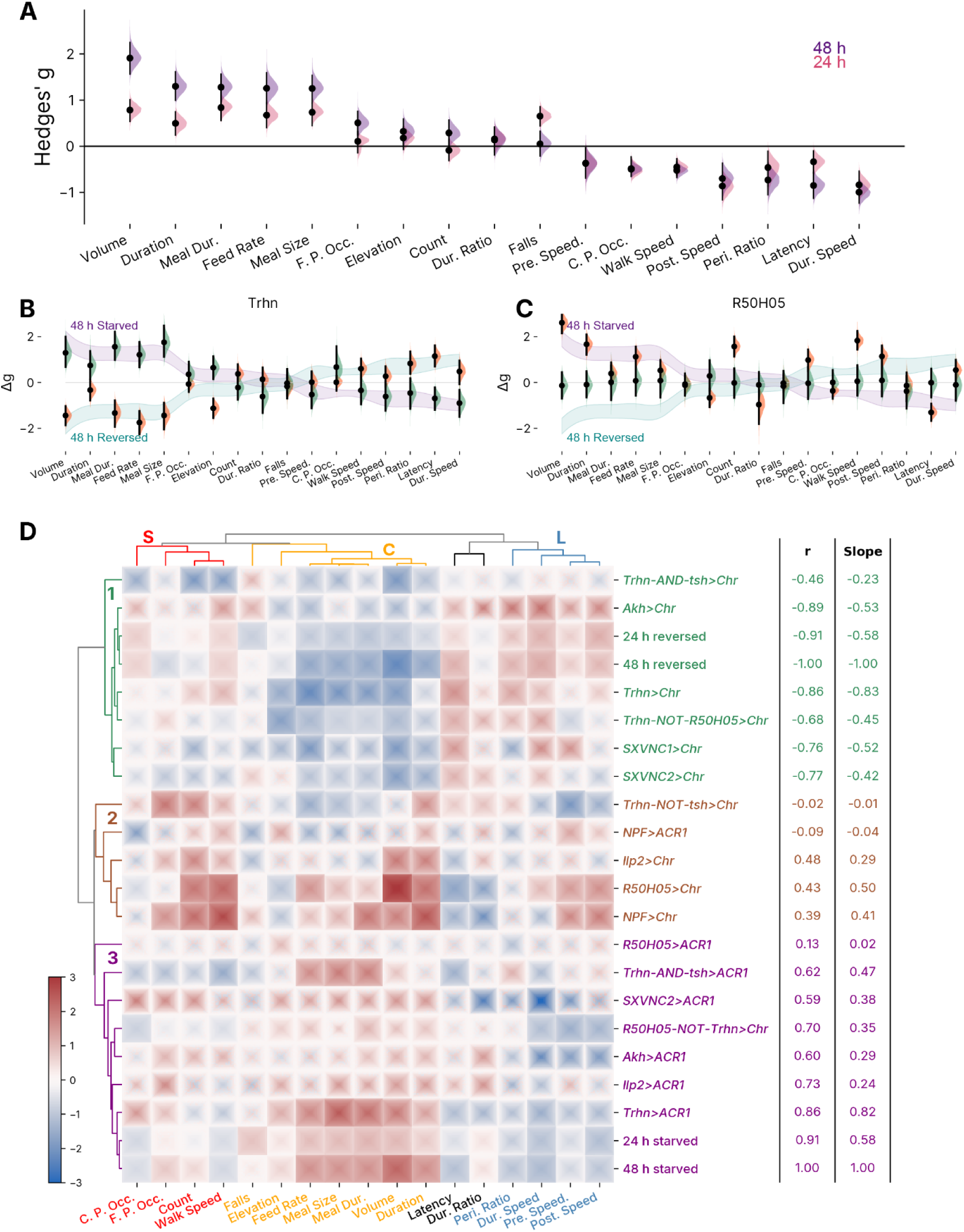
Ethograms reveal that *Trhn-Gal4* circuit activity mimics natural satiety. Ethogram analyses of feeding-behavior changes arising from optogenetic manipulations and starvation. **A,** Forest plot of state transition vectors using Hedges’ g of 17 behavioral changes after 24-h and 48-h starvation. Each half-violin curve represents the effect size distribution for that metric. Black dots indicate the mean effect size; vertical bars show the confidence interval. The metrics include volume, duration, meal size, and locomotor attributes, which are ranked by effect magnitude in 48-h starvation, from largest increase to largest decrease. **B,** Overlaid forest plots for *Trhn-Gal4* optogenetic effects, showing the effects of activation (fed, *Chr*, orange) and inhibition (starved, *ACR1*, green), ordered by the 48-h effect ranking. The 48-h starved phenovector ribbon (purple) and ‘reversed-starvation’ ribbon (teal) are drawn to show the natural-hunger profile. **C,** Overlaid forest plots of *R50H05* optogenetic effects, following the same format as panel **B** with *Chr* effects (orange) and *ACR1* effects (green). The purple and teal ribbons indicate the 48-h effects profile for starvation and its inverse, respectively. **D,** Correlation hierarchical clustermap of Hedges’ g effect sizes from both natural and optogenetic experiments. The colors indicate effect sizes, with red and blue indicating the positive and negative values (respectively) of the bootstrapped effect-size (Δg) distribution. Dendrograms illustrate the proximity between behavioral metrics (top) and experimental conditions (left). Metrics are grouped into three clusters: food-seeking (S, red), food consumption (C, yellow), and locomotor attributes (L, blue and black). Experimental conditions are segmented into three clusters, highlighting similarities between hunger–satiety transitions and optogenetic manipulations: cluster 1 includes the reverse-starvation 24-h and 48-h hunger→satiety transition vectors (24 h reversed, 48 h reversed); cluster 2 contains no state vectors; and cluster 3 contains the two satiety→hunger transition vectors (24 h starved, 48 h starved) and the phenovectors of starved optogenetic flies. Several behavioral metrics are abbreviated: Meal Dur. = Meal duration, Pre. Speed = Prefeed speed, Dur. Speed = Duringfeed speed, Post. Speed = Postfeed speed, Dur. Ratio = Duringfeed speed ratio, Peri. Ratio = Peri-feed speed ratio, F. P. Occ. = Food port occupancy, C. P. Occ. = Control port occupancy. Adult male flies (5–10 days post-eclosion) were used throughout.

**Figure S7.**
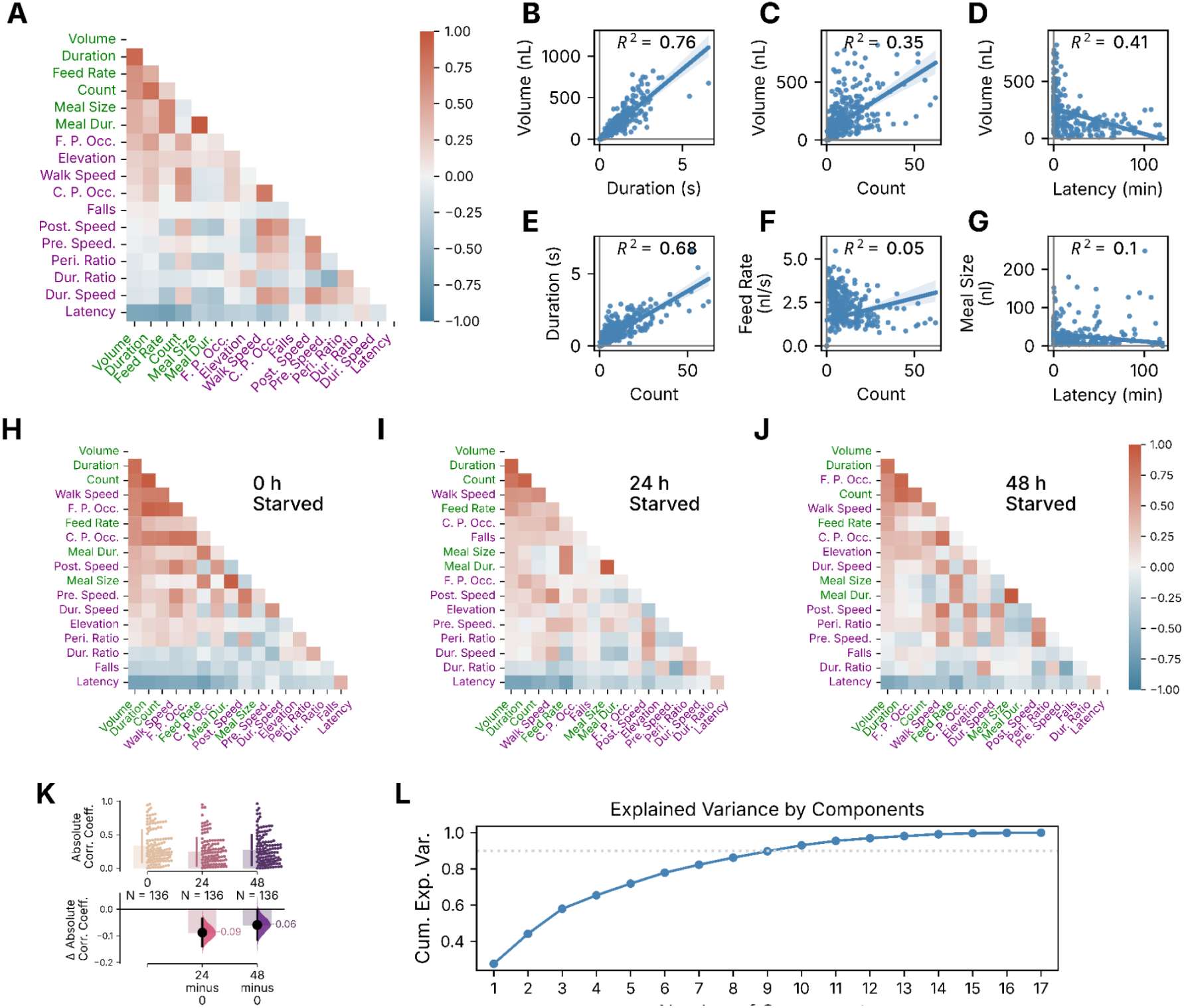
Correlation analysis of feeding-related behaviors indicates limited redundancy. **A**, Correlogram displaying the correlations between various feeding and behavioral metrics in fed flies. The color intensity and hue (red for positive, blue for negative) indicate the strength and sign of the correlations between different metrics, such as volume, duration, feed rate, and latency. Sample size N = 310 flies. Feeding-related metrics are colored in green in axes and locomotion related metrics are colored in purple. **B,** Scatter plot showing that volume and duration are highly correlated. Each dot represents the mean value from an individual fly. The line indicates the regression fit and the shaded region represents the 95CI for the regression line. **C,** Scatter plot illustrating that volume and count are moderately correlated. **D,** Volume and feeding latency are moderately correlated. **E,** Duration and feed count are strongly correlated. **F,** Feed rate and count are correlated very weakly. **G,** Meal size and latency are weakly correlated. **H,** Correlogram in fed flies (starved 0 h). Sample size N = 150 flies. The color scale and arrangement are as above. **I,** Correlogram of metrics in flies that were starved for 24 h. Sample size N= 120. **J,** Correlogram of metrics in 48-h starved flies. Sample size N = 90. **K,** An estimation plot showing the absolute correlation coefficient values between pairs of metrics in different nutritional conditions (0-, 24-, and 48-h of starvation). The upper axis shows correlation values between 136 pairs of metrics, with standard deviations (lines), and mean values (bars). The lower axis shows the mean differences (black dots), 95CIs (black lines), and effect distribution (half-violin curves); numerals are effect sizes (ΔR^2^). **L,** Line plot of cumulative variance explained by principal components. The x-axis represents the number of components; the y-axis shows the cumulative explained variance. The horizontal dashed line shows 90% variance explained. Adult male flies (5–10 days post-eclosion) were used for feeding experiments (panels E–F).

**Figure S8.**
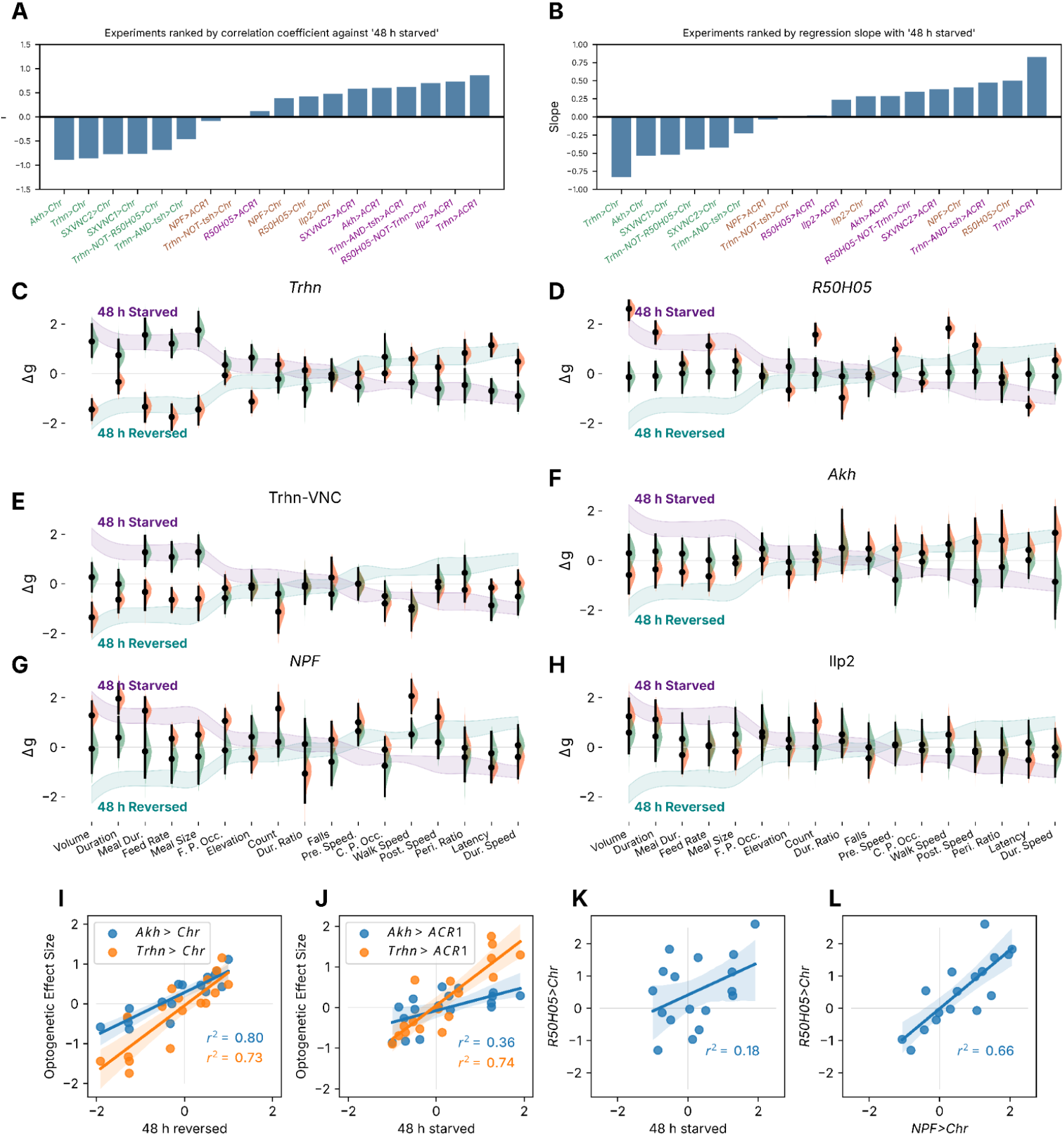
Comparing optogenetic behavioral profiles with hunger–satiety transitions. **A**, Correlation coefficients from linear regression of optogenetic phenovectors against 48-h starvation effects. Coefficients are ranked, showing the strength and sign of the relationship between each experimental condition and hunger. Text labels on x-axis are colored green, brown or purple, for cluster 1, 2, and 3, respectively. **B,** Slopes (β) from the same linear regressions, indicating the magnitude of alignment with the starvation response. Slope values are ranked; text labels on x-axis are colored by cluster. **C,** Forest plot of effect sizes for *Trhn*>Chr under starved conditions (orange) and *Trhn*>ACR1 under fed conditions (green). The ribbons outline the 95CIs for hunger–satiety transitions between 48-h starved (purple) and the inverse effects (teal). This panel is shown in Figure 5B, it is re-plotted here for easy comparison. **D,** Forest plot of effect sizes for *R50H05*>*Chr* (fed, red) and *R50H05*>*ACR1* (starved, green). This panel is shown in Figure 5C, it is re-plotted here for easy comparison. Effect sizes are co-plotted with the 48-h starvation and reverse CI ribbons. **E,** Forest plot of effect sizes for *Trhn-*AND*-tsh*>*Chr* (starved, orange) and *Trhn-*AND*-tsh*>*ACR1* (fed, green), co-plotted with the hunger-transition ribbons. **F,** Forest plot of phenovectors for *Akh*>*Chr* (starved, orange) and *Akh*>*ACR1* (fed, green). **G,** Forest plot of phenovectors for *NPF*>*Chr* (fed, orange) and *NPF*>*ACR1* (starved, green). **H,** Forest plot of phenovectors for *Ilp2*>*Chr* (fed, orange) and *Ilp2*>*ACR1* (starved, green). **I,** Scatter plot with a regression analysis that compares effect sizes of *Akh*>*Chr versus* 48-h reversed (blue) and *Trhn*>*Chr versus* 48-h reversed (orange). Dots represent individual effect sizes of individual behavioral features, lines indicate the regression fit line, and shaded regions represent the 95CI of the fit. **J,** Scatter plot comparing effect sizes of *Akh>ACR1* fed *versus* 48-h starved (blue) and *Trhn>ACR1* fed *versus* 48-h starved (orange). **K,** Scatter plot comparing effect sizes of *R50H05>Chr* fed *versus* 48-h starved. **L,** Scatter plot comparing effect sizes of *R50H05>Chr* fed *versus NPF>Chr* fed. Adult male flies (5–10 days post-eclosion) were used throughout.

**Figure S9.**
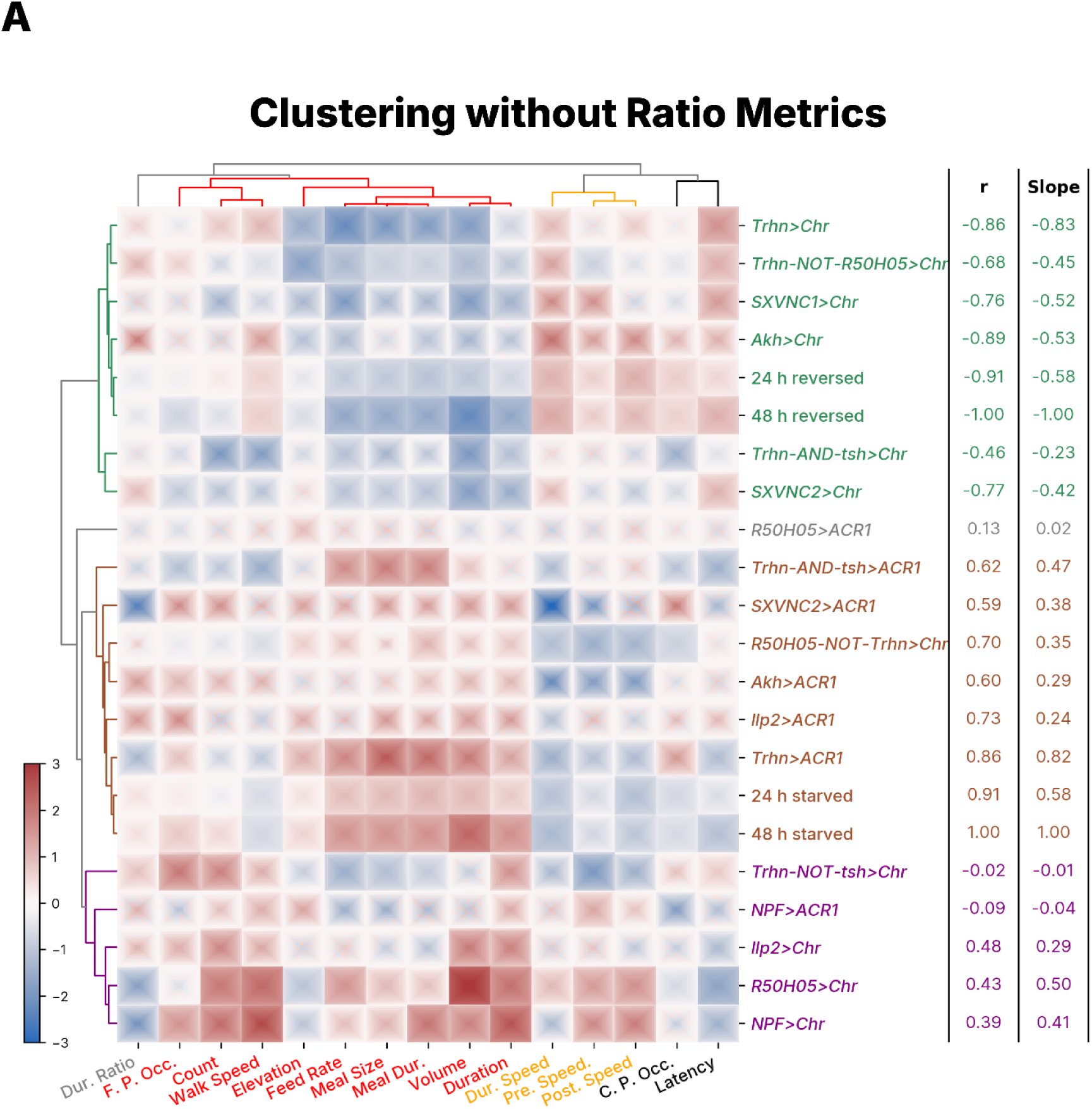
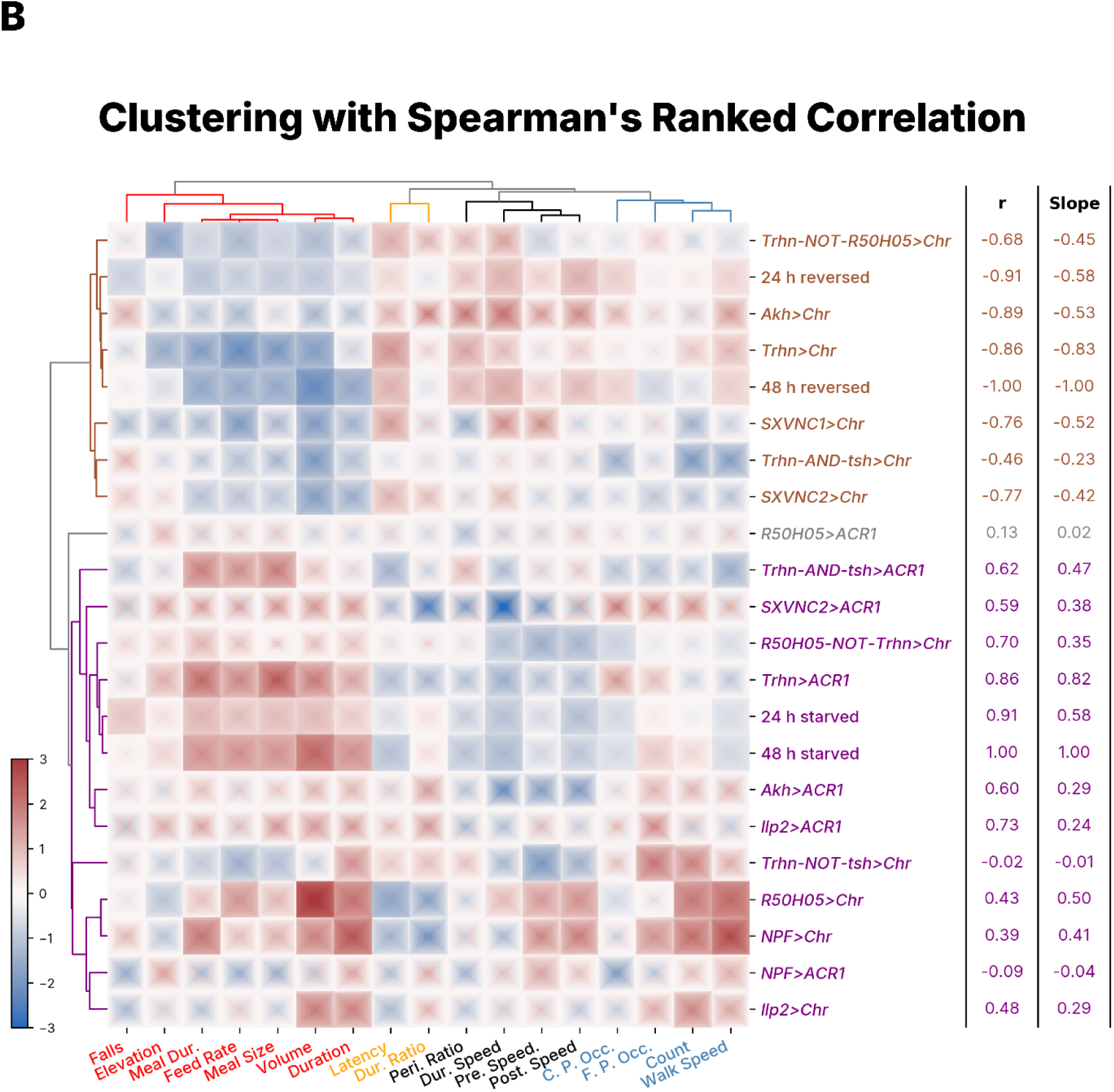
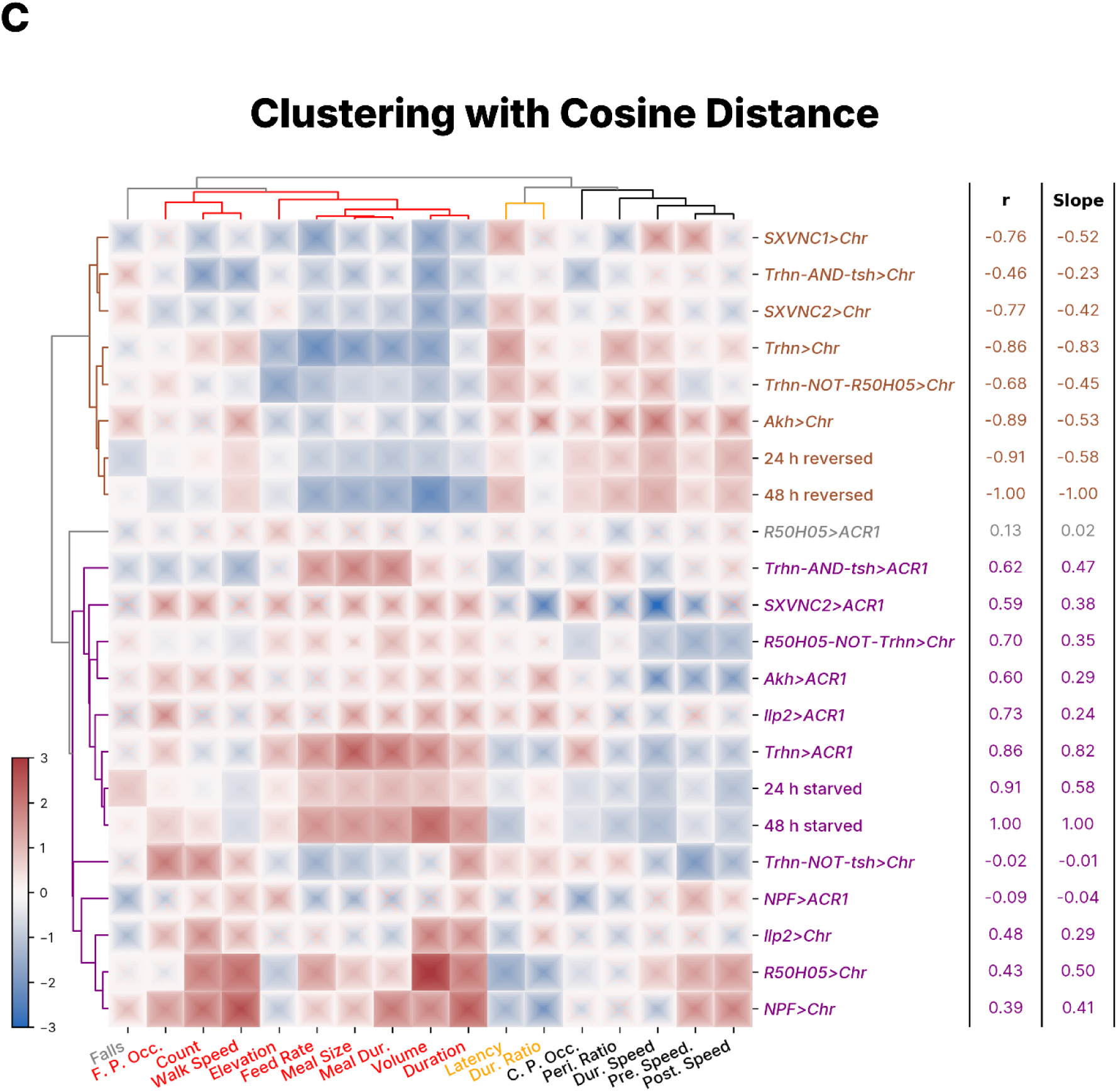
Robustness of phenovector analysis. Ethogram analyses of the same data as Figure 5D using different metrics and clustering methods. Correlation hierarchical clustermap of Hedges’ g effect sizes from both natural and optogenetic experiments. The colors indicate effect sizes, with red and blue indicating the positive and negative values (respectively) of the bootstrapped effect-size (Δg) distribution. Dendrograms illustrate the proximity between behavioral metrics (top) and experimental conditions (left). Metrics are grouped into three clusters: food-seeking (S, red), food consumption (C, yellow), and locomotor attributes (L, blue and black). Experimental conditions are segmented into three clusters, highlighting similarities between hunger–satiety transitions and optogenetic manipulations: cluster 1 includes the reverse-starvation 24-h and 48-h hunger→satiety transition vectors (24 h reversed, 48 h reversed); cluster 2 contains no state vectors; and cluster 3 contains the two satiety→hunger transition vectors (24 h starved, 48 h starved) and the phenovectors of starved optogenetic flies. Several behavioral metrics are abbreviated: Meal Dur. = Meal duration, Pre. Speed = Prefeed speed, Dur. Speed = Duringfeed speed, Post. Speed = Postfeed speed, Dur. Ratio = Duringfeed speed ratio, Peri. Ratio = Peri-feed speed ratio, F. P. Occ. = Food port occupancy, C. P. Occ. = Control port occupancy. **A,** Clustering using all metrics except Dur. Ratio and Peri. Ratio to test for redundancy of these two metrics. **B,** Clustering using Spearman’s Ranked Correlation. **C,** Clustering using cosine distance.

### Serotonin is a satiety factor that accumulates during starvation

Having established that *Trhn*^+^ serotonergic cells regulate the hunger–satiety axis, we next investigated their mechanism of action. We reasoned that serotonin levels could be a molecular correlate of nutritional state and we hypothesized that serotonin levels would be higher in sated flies. Surprisingly, immunostaining revealed that compared to fed controls, 24-h starvation elevated serotonin immunoreactivity by 74% in the brain and 63% in the VNC (**Figure 6 A - D**), in other words, in the CNS as a whole. Starving flies for 24 hours and refeeding them for 1 hour abolished this effect (**Figure S10 A - C**).

Because many neuromodulatory neurons employ co-transmitters, we wanted to check whether serotonin *per se* is required for feeding suppression. To do so, we knocked down endogenous *Trhn* mRNA in *Trhn-Gal4* neurons using an RNA-interference (RNAi) transgene. In an Espresso assay, we saw that this knockdown increased the feeding of fed flies when compared to controls (Δ = +85 nL; **Figure 6E**). This increase was smaller than the ACR1 inhibitory effect (85 < 165 nL; **Figure 2G**) but nevertheless demonstrates that serotonin is a key neurotransmitter for feeding suppression during the sated state. We also used RNA interference to knock down Trhn-dependent serotonin synthesis in *Trhn* VNC neurons in fed *Trhn*-AND-*tsh*>*Trhn-RNAi* flies. The resulting knockdown flies ate +69 nL more food, showing that the serotonergic VNC neurons use 5-HT during satiety suppression.

These results show that serotonin is an important satiety effector of *Trhn^+^* VNC neurons. The high level of serotonin in starved flies (**Figure 6A, B**) suggests that, during hunger, 5-HT synthesis continues while serotonin release (and/or degradation) is inhibited due to a lack of neuronal activity (*46*).

### A sugar-sensing mechanism for serotonergic feeding control

In our final analyses, we investigated how satiety-inducing serotonergic neurons sense metabolic state. Prior work has shown that cell-autonomous nutrient sensing has a role in other feeding-related cells, including SLC5A11 in the ellipsoid body (*47*), sugar transport in Dh44 cells (*48*), and enteroendocrine cells (*44*). We therefore made *ad hoc* selections of two sugar transporters, *Glucose transporter 1* (*Glut1* (*49*)) and *sugar transporter 2* (*sut2* (*50*)), and tested the effects of knocking them down in *Trhn*^+^ neurons. Knocking down *Glut1* (*51*) in all *Trhn-Gal4* neurons had a negligible effect on feeding (Δ = –23 nL [95CI –50, +10]), while its knockdown in VNC cells had a modest effect (though the estimate lacks precision; Δ = +41 nL [95CI –10, +90]; **Figure 6F**). Knocking down *sut2* in *Trhn-Gal4* cells also yielded a modest effect (Δ = –58 nL [95CI –160, +30]), yet knockdown in VNC 5-HT^+^ neurons drove a clear increase in feeding (Δ = +57 nL [95CI +30, +89]; **Figure 6F**). These results indicate that the satiety-inducing function of *Trhn* VNC neurons likely depends on sugar transport *via* the Sut2 transporter. Based on these results, we propose that Sut2 performs a sugar-sensing role: when nutrient levels are high, intracellular sugar contributes to the activation of *Trhn^+^* VNC neurons, which suppresses feeding. Comparing the *Glut1* knockdown phenovectors against the 48-h starvation vector revealed a moderate alignment for *Trhn*-AND-*tsh*>*Gluti* (R² = 0.14) but near-zero alignment for broad *Trhn*>*Gluti* (R² = 0.00), suggesting that *Glut1* knockdown in VNC-restricted neurons produces a partial hunger-like profile whereas broad knockdown does not (**Figure 6G**). By contrast, *sut2* knockdown in *Trhn*-AND-*tsh* neurons showed stronger alignment with the starvation vector (R² = 0.42), compared with negligible alignment for broad *Trhn*>*sut2i* (R² = 0.06), consistent with *sut2* acting specifically within VNC serotonergic neurons to mediate sugar sensing (**Figure 6H**).

**Figure 6.**
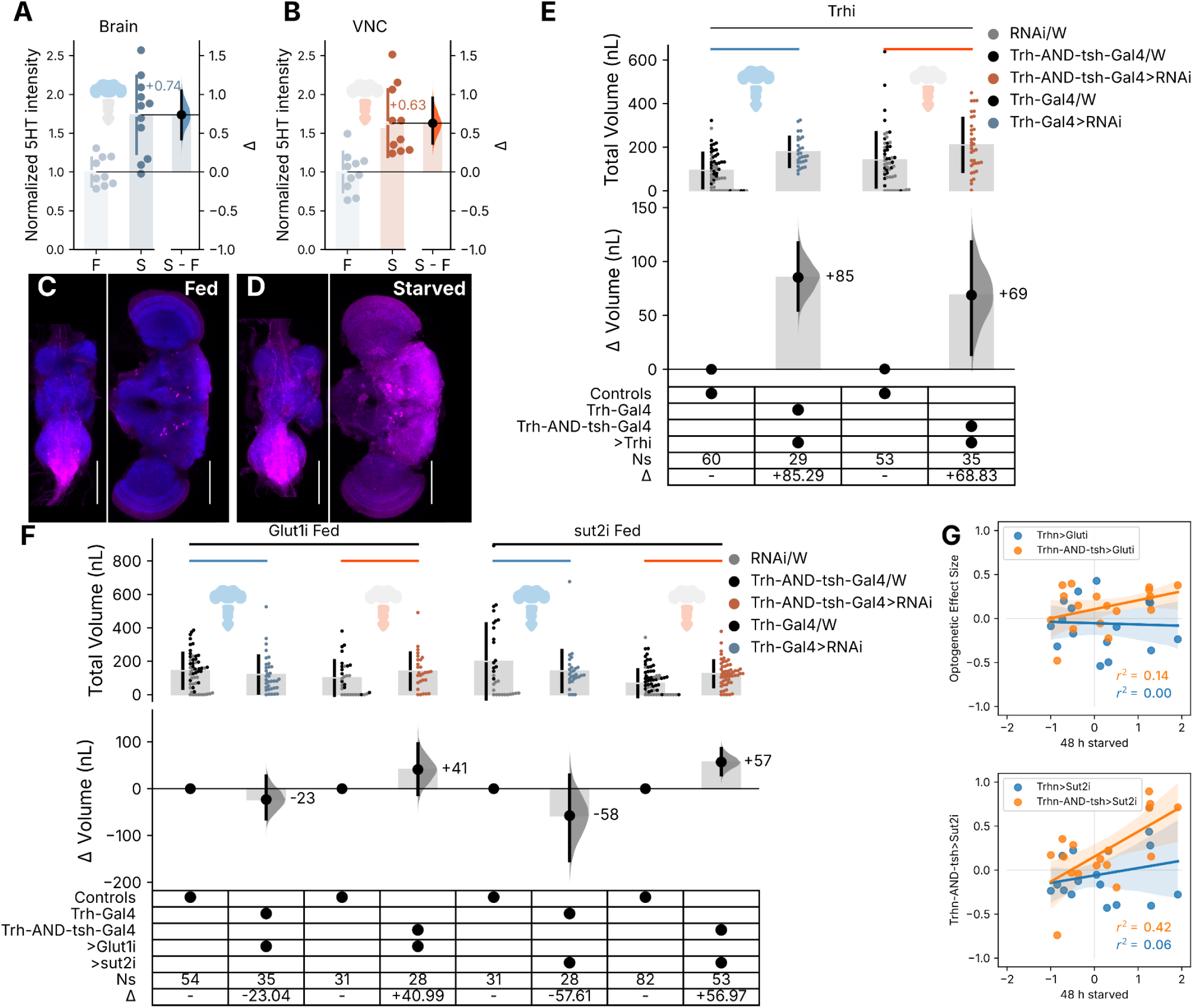
Molecular mechanism of *Trhn* neuronal suppression of feeding. **A**, A Gardner-Altman estimation plot showing the normalized intensity of 5-HT staining in the brain of fed and starved *w^1118^* males. The left axis shows a swarm plot of raw data, and the right axis shows a half violin plot of the estimated effect size. Numbers immediately left to the half violin indicate the mean effect size. Sample sizes: N_Fed_ = 10, N_Starved_ = 11. **B**, Estimation plot of 5-HT staining intensity in the VNCs of the same fed and starved *w^1118^* male flies. **C**, Representative images of stained VNC and brain from a fed male fly. Magenta is α-5-HT signal; blue is nc82 α-BRP signal. Scale bar = 100 µm. **D**, Representative images of VNC and brain from starved flies. Visual elements as per panel **C**. **E**, A Cumming estimation plot showing the effects of feed volume in controls and RNAi knockdown of *Trhn* in *Trhn-Gal4* (left, blue) and *Trhn*-AND-*tsh* (right, red) neurons. The top panel shows a swarm plot of mean feed volumes consumed (nL), and the bottom panel shows a contrast plot of the effect sizes. Numbers left of each half-violin curve indicate the mean effect size. The key table below the plot shows the presence of the driver and responder alleles, as well as sample sizes and effect sizes. ‘*Trhi*’ = *UAS-Trhn-RNAi*. **F**, Estimation plot showing the effects of feed volume as a result of *Glut* and *sut2* knockdowns (*UAS-Glut-RNAi* and *UAS-sut2-RNAi*, respectively) in *Trhn* and *Trhn*-AND-*tsh* neurons. The top panel shows a swarm plot of raw data, and the bottom panel shows a contrast plot of the effect sizes. Numbers left of half violin indicate the mean effect size. The table below the plot shows the presence of the driver and responder alleles, as well as sample sizes and effect sizes. The transgenes in the key table are abbreviated: *‘Trhn’* = *Trhn-Gal4*; *‘tsh’* = [*tub-frt-Gal80-frt; tsh-LexA>lexAop-FLP*]; ‘*Gluti’* = *UAS-Glut-RNAi*; and ‘*sut2i’ = UAS-sut2-RNAi*. For **E** and **F**, data with controls separately plotted are shown in **Figure S10**. **G,** Scatter plot of *Trhn>Gluti versus* 48-h starved (blue), as well as *Trhn*-AND-*tsh>Gluti versus* 48-h starved (orange). **H,** Scatter plot of *Trhn>sut2i* and 48-h starved (blue), as well as a comparison of *Trhn*-AND-*tsh>sut2i* and 48-h starved (orange). Dots denote individual effect sizes; straight lines denote regression lines; shaded regions denote 95CI of the regression lines. Adult male flies (5–10 days post-eclosion) were used throughout. Optogenetic effects with controls separately plotted are shown in **Figure S10**.

**Figure S10.**
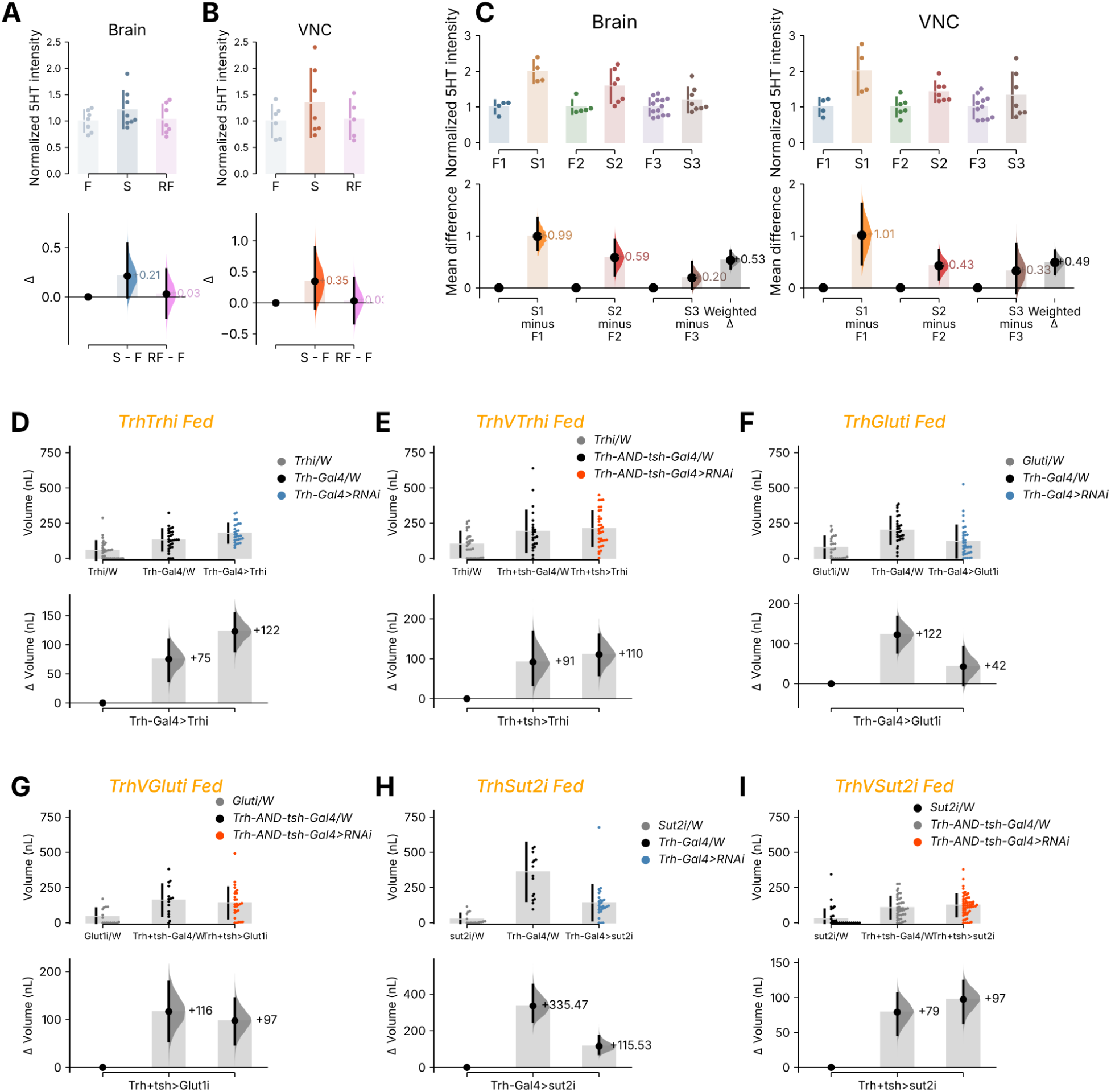
5-HT immunoreactivity refed experiment and Separated genetic controls for RNAi knockdown experiments. (**A - B**) Estimation plots showing a modified replicate of the experiment shown in Figure 6A**, B**. In addition to the fed and starved groups, an additional group of flies were starved and refed liquid food as that was used in Espresso. The refed flies showed similar immunoreactivity for 5-HT to the fed group, presumably as a return to baseline. **C,** A meta-analysis of the three replicates of 5-HT starvation immunoreactivity experiments as presented in Figure 6A**, B** and **Figure S10 A, B**. Upper panels show individual sample values for normalized 5-HT intensity. Bottom panels show the mean (black dots), the bootstrap distribution (half violins) and 95CI (black vertical bars) of the mean difference effect size for each pair of conditions within a replicate. The gray bar on the far right shows the weighted delta calculated from all replicates (0.53 for brain samples and 0.49 for VNC samples). (**D - I**) Estimation plots showing the effects of RNAi-mediated knockdown on feeding volume, with Driver Control and Responder Control displayed as separate columns to allow assessment of genetic background differences. Each panel includes a swarm plot showing individual measurements (top) and a contrast plot depicting the effect sizes (bottom). In the swarm plots, vertical bars represent the standard deviation, with the central gap indicating the mean value. In the lower axes (contrast plots), black dots denote mean differences, black bars represent the 95CI, and half-violins show the bootstrapped difference distributions. Numbers immediately adjacent to the half-violin curves are the mean effect sizes. **D,** Estimation plot of feed volume in fed *Trhn*>*Trhn-RNAi* flies with Driver and Responder Controls shown separately. **E,** Estimation plot of feed volume in fed *Trhn*-AND-*tsh*>*Trhn-RNAi* flies with separated controls. **F,** Estimation plot of feed volume in fed *Trhn*>*Glut1-RNAi* flies with separated controls. **G,** Estimation plot of feed volume in fed *Trhn*-AND-*tsh*>*Glut1-RNAi* flies with separated controls. **H,** Estimation plot of feed volume in fed *Trhn*>*sut2-RNAi* flies with separated controls. **I,** Estimation plot of feed volume in fed *Trhn*-AND-*tsh*>*sut2-RNAi* flies with separated controls. Adult male flies (5–10 days post-eclosion) were used throughout.

## Discussion

Single-metric assays risk misidentifying isolated effects as genuine state changes. Using DESTRA, we show that while several feeding-related circuits produced phenovectors discordant with natural hunger–satiety transitions, Trhn neurons bidirectionally and faithfully recapitulated the coordinated behavioral signature of natural satiety.

### Ethomics detect coordinated behavioral changes in authentic state transitions

Internal states like hunger and satiety produce coordinated behavioral changes across multiple dimensions (*2*, *29*, *52*). Early rodent studies showed that satiety involves not only reduced feeding but also postprandial behaviors, such as grooming and rest (*13*, *53*). Machine vision and deep learning now enable automated, high-dimensional behavioral quantification. This approach, termed computational ethology or ethomics, has facilitated integrative behavioral analysis (*30*, *31*, *54*, *55*). Such systematic analysis of behavioral pleiotropy is part of the broader program of phenomics, the large-scale study of phenotypes (*56–58*). In flies, ethomics using unsupervised-defined activity categories (*32*) or human-identified motor and social-interaction features (*33*) has facilitated large-scale behavioral screens to study circuit interventions.

In our analysis of hunger-satiety pleiotropy, specific feeding-related features merit discussion. First, hungry flies walked more slowly after a meal than before (**Figure 1L**), supporting previous findings on postprandial sleep (*27*, *34*) and confirming that postprandial quiescence begins immediately after feeding. Such meal-related speed reduction is enhanced in hungry flies (**Figure 5A**). Second, feed count remained largely unchanged in starved flies, suggesting it is not a key marker of the satiety→hunger state transition. Notably, an increased feed count might betray a fragmented feeding state, as both *R50H05>Chr* and *NPF>Chr* flies exhibited higher feed counts despite overall phenovector discordance with state vectors (**Figure S8 D, G**). Third, as in humans (*59*), hunger has a robust effect on meal size rather than frequency. Previous *Drosophila* studies linked meal size to food choice (*60*) and the leucokinin pathway (*61*) but our findings provide the first evidence that hunger regulates meal size. Rather than increasing meal frequency, hungry flies consume larger meals, which are typically followed by rest. Importantly, although the 17 metrics fall into two categories—feeding and locomotion—these metrics do not necessarily move in lockstep under optogenetic perturbation: *R50H05*>*Chr* activation simultaneously drove large increases in walking speed and dispersal yet increased food port occupancy and feeding frequency, demonstrating that even strongly correlated metrics within a block can be independently perturbed. A manipulation recapitulating the full coordinated profile is more likely to reflect a genuine state shift than one that moves only a subset.

Consumption is the most salient readout of hunger and a reasonable starting point for assay development (*25*). The value of the additional metrics emerged precisely because consumption alone proved inadequate to distinguish authentic state changes from isolated feeding effects. This is most clearly illustrated by *R50H05* and *NPF* activation, which both drive consumption increases that are, in isolation, indistinguishable from hunger, yet whose phenovectors reveal a divergent pattern from starvation states.

### Using natural state transitions as ground truth for neurogenetic ethomics

Our study extends the ethomics framework by introducing DESTRA: using natural state-transition vectors as ground truth for evaluating phenovectors elicited by neurogenetic manipulations. There are three enabling aspects: First, from automated tracking data, we calculated a large number of behavior metrics that aimed to capture hunger’s pleiotropic nature. Second, we used graded starvation to produce shifts along the hunger–satiety axis, giving references against which to benchmark perturbations. Third, by utilizing standardized effect sizes, we placed optogenetic phenotypes and natural states on the same scale of hunger–satiety state-transition reference vectors, enabling direct assessment of how faithfully different neural interventions recapitulate natural behavioral programs. Instead of comparing single metrics in isolation, we use phenovectors to estimate a ‘state-likeness’ (**Figure 5A**).

### Activity in the *Trhn* neurons phenocopies natural satiety

Two prior studies identified fly serotonergic neuron populations with diverse effects on memory, activity, quiescence, and courtship, including diametrically opposite effects on feeding (*20*, *21*). Our results not only reproduce the opposing feeding effects of the two lines but also reconcile this apparent discrepancy by showing that feeding changes arise from non-overlapping serotonergic cell groups, providing new insight into serotonergic locomotor control.

Previous research had suggested that feeding reductions due to serotonergic activation in *Trhn*>*TrpA1* flies derived from a generalized state of activity quiescence (*20*). DESTRA analysis challenges this interpretation, finding that *Trhn-Gal4* activation induces coordinated behavioral changes that closely mimic natural satiety and postprandial quiescence without inducing extended generalized quiescence, while inhibition of these neurons shifts flies into a hunger-like state (**Figures 5 and S8**). This finding supports a causal sequence: *Trhn*-cell activity → satiety brain state → locomotor effects. *Trhn* activity is both required for and instructive of the natural satiety state. This remarkable bi-directionality is consistent with the hypothesis that these neurons occupy a hub position in the state-control network.

Mechanistically, serotonin immunoreactivity was elevated in starved flies (**Figure 6A, B**), leading us to speculate that 5-HT was accumulated in *Trhn⁺*neurons during food deprivation. We could not further investigate this mechanism, although prior research suggests accumulation with reduced release, reduced degradation, increased re-uptake, and/or increased synthesis (*46*), which point to possible future experiments. Regardless, a reservoir of 5-HT in starved *Trhn⁺* neurons is consistent with the finding that optogenetic activation was effective in the starved state: forcibly opening *Chrimson* channels in neurons that have accumulated serotonin would release that reservoir, producing the robust feeding suppression we observe (**Figure 2F**). The converse logic applies to *ACR1* inhibition in fed flies: when *Trhn* neurons are otherwise active and releasing 5-HT tonically, inhibition of these neurons increases feeding (**Figure 2G**), consistent with removal of a persistent serotonergic satiety brake. On the other hand, synthase knockdown mimics the ACR1 experiment by reducing the source of 5-HT releasable. Neurochemical and behavioral data thus corroborate: the former establishes that serotonin is available in the starved state, and the latter demonstrates that releasing it is sufficient to suppress feeding.

Our findings on the fly serotonergic satiety system align with vertebrate data on serotonin’s role in satiety signaling. For example, in rodents, serotonergic circuits suppress feeding and promote postprandial quiescence (*15*, *28*), mirroring our *Trhn>opsin* results. Similarly, serotonergic neurons in the zebrafish caudal hypothalamus mediate satiety (*62*). While serotonergic regulation of feeding varies across invertebrate taxa and circuits (*23*, *24*), our findings suggest that, as in vertebrates, fly *Trhn* neurons orchestrate hunger–satiety state transitions.

### Other feeding circuits produce dissociated phenovectors

Activation of the other serotonergic driver we tested, *R50H05*, was previously interpreted as driving a hunger-like state (*21*). In contrast, our analysis revealed two dissociations: (1) *R50H05*>*Chr* flies exhibited heightened locomotion when food was available, whereas genuinely hungry flies focused their activity around the food port (**Figures 5 and S8**); and (2) *R50H05>ACR1* inhibition had no impact on feeding behavior, suggesting these neurons are not required for natural hunger. *R50H05>Chr* activation increased food consumption and feeding frequency, raising the possibility that these neurons drive a form of homeostatic hunger where energy balance is perturbed and feeding is upregulated as a compensatory response; the accompanying locomotor profile including generalised dispersal and elevated walking speed rather than food-port-directed movement is inconsistent with AHLS in the presence of food, and the *R50H05>Chr* phenovector clusters separately from authentic state transitions. We therefore favor the interpretation that R50H05 neurons modulate locomotion and incidentally increase feeding, rather than acting as nodes in the hunger–satiety axis *per se*.

Other known feeding neurons exhibited surprising optogenetic phenovectors (Figure 5D), and sometimes cluster quite far away from natural state transitions. *Ilp2* activation in fed flies increased feeding but also generated erratic locomotor patterns, while *Ilp2* inhibition produced hunger-like effects with a phenovector clustered with natural satiety→hunger transitions. These results indicate that *Ilp2* neuronal activity is required for the satiety state but their activation produces a dissociated behavioral profile inconsistent with natural hunger. Notably, *Akh>Chr* had a phenovector most similar to both *Trhn>Chr* and the natural hunger→satiety transition; this is consistent with *Akh’s* established role as a glucagon-like peptide that suppresses sugar appetite. However, *Akh* inhibition in starved flies, had minimal effects on feeding behavior, suggesting these neurons’ activity is not a major requirement for maintaining satiety. This is in contrast to the stark bidirectional effects from *Trhn* neurons. *NPF* manipulations revealed perhaps the most surprising results: activation increased feeding frequency but also elevated locomotion and dispersal, creating a behavioral profile inconsistent with an energy-conserving hunger state. Notably, the *NPF-Gal4* effects closely resembled those of *R50H05-Gal4* (R^2^ = 0.66), suggesting both populations drive locomotion-dominant states distinct from AHLS. *NPF* inhibition likewise failed to reduce feeding in hungry flies, indicating these neurons are not required for the natural hunger drive. Lastly, that we observe several lines where feeding frequency and volume were dissociated provides a starting point for investigating possible diverse modes of feeding increases, which could map onto distinct circuit architectures.

### The role of VNC serotonergic neurons and sugar sensing in feeding suppression

The intersectional optogenetic and knockdown data suggest that VNC serotonergic neurons act as nutrient sensors. When sugar levels are high, they contribute to the satiety state, and, in starved flies, their silence permits hunger-drive behavior. Three key observations support this interpretation. First, activating these neurons using multiple targeted drivers strongly suppressed feeding (**Figure 4**). Second, the phenovectors of fly lines with VNC-specific serotonergic activation were similar to hunger→satiety state transitions (e.g., *SXVNC2>Chr vs.* 48-h reversal; R^2^ = 0.59; **Figure 5D**). Third, neuronal inhibition, 5-HT synthesis knockdown, and targeted *sut2* knockdown in the VNC of fed flies all increased feeding, indicating that these neurons normally help to maintain satiety (**Figure 4 and Figure 6E, F**). The broad-driver *Trhn>sut2i* effect (Δ = –58 nL, 95CI spanning zero, **Figure 6F**) should be interpreted with caution as it possibly reflects simultaneous targeting of two functionally divergent populations: VNC *Trhn* cells, whose *sut2* knockdown increases feeding, and *R50H05*-positive brain cells, whose *sut2* disruption may have opposing effects—potentially cancelling the VNC-specific signal and producing the near-null net result we observe. The intermediate and heterogeneous phenotype of Trhn-NOT-tsh>Chr likely reflects a combination of incomplete VNC repression and opposing behavioral contributions from R50H05-positive brain Trhn neurons, consistent with the near-null net effect observed in broad *Trhn>sut2i* knockdown, where simultaneous targeting of functionally divergent populations may cancel opposing signals. It also suggests that thoracic VNC serotonergic neurons are the primary contributors to the Trhn-AND-tsh phenotype.

Several lines of evidence from our study, however, indicate that serotonergic VNC neuron control of satiety is incomplete. VNC-specific inhibition increased food consumption by only about a quarter of the effect seen with broad *Trhn>ACR1* inhibition (**Figure 4**). Similarly, our comparisons of phenovectors suggested only partial VNC serotonergic control of satiety state (e.g., *SXVNC2>ACR1 vs.* 48-h starvation, *R^2^* = 0.35; **Figure 5D**). Also notable, *SXVNC2*-*Gal4* may be labeling ejaculatory VNC neurons and therefore may not regulate AHLS, but may induce a phenotype through secondary effects. We thus conclude that serotonergic VNC cells are an important, though not unique, component of a distributed serotonergic system that has control over hunger–satiety state transitions.

These findings align with prior research on brain serotonergic circuits, particularly in the subesophageal zone, that regulate nutrient sensing, control feeding, and limit sugar ingestion (*63–65*). Additionally, brain serotonergic neurons suppress feeding-related behaviors (*66*). The modest effects we observed with brain-only expression (*Trhn*-NOT-*tsh*>*Chr*) suggest additional complexity in how different serotonergic populations coordinate feeding, highlighting the need for further investigation.

### Study Limitations

At least seven methodological limitations warrant consideration when interpreting our findings. First, the vertical orientation of Espresso chambers could affect feeding measurements if interventions impair climbing ability. Our analysis of port visits helped control for this, showing no obvious deficits in the lines used; future horizontal chamber designs may further mitigate the issue. Second, while we did measure walking, we cannot rule out fine motor deficits, such as proboscis extension impairments, that could influence feeding behavior. Third, although liquid food allows precise measurements, it differs from flies’ natural diet of fermenting fruit, potentially masking texture- or limb-gustation effects. Fourth, continuous optogenetic activation might oversimplify serotonin’s natural dynamics, as phasic or rhythmic activity patterns could be essential for modulating behavior. Fifth, although our 17-metric ethomics approach is more comprehensive than prior methods, it might still miss relevant behavioral features; additional methods such as key-point tracking (*67*), expanded time-series analyses, and machine-learning classification could reveal potentially relevant features (*2*, *31*, *55*, *68*). Sixth, the DESTRA reference vector is derived from a single wild-type strain under specific starvation conditions; alternative durations, food compositions, or environmental conditions could shift it, and metric selection and clustering method affect driver ordering. Seventh, most intersectional manipulations were tested with Chrimson activation rather than ACR1 inhibition, reflecting an initial screen for sufficiency across a combinatorially large driver space; inhibition counterparts, which would establish necessity for each subset, remain an important direction for future work. Despite these limitations, multiple factors strengthen our conclusions: consistency with prior studies showing that different serotonin drivers have opposite feeding effects (*20–22*), replication across different Boolean genetic tools, and the striking observation that both *Chr* and *ACR1* phenovectors for *Trhn-Gal4* recapitulate natural hunger–satiety state transitions. Together, they establish a key serotonergic circuit as an authentic controller of feeding.

### A minimal architecture for behavioral state control

Our findings demonstrate that a small set of neuromodulatory cells (*Trhn^+^* cells) bidirectionally regulate the hunger–satiety axis, showing that fundamental states can emerge from the activity (or quiescence) of a compact neural population. The partial recapitulation of satiety by VNC serotonergic neurons suggests a distributed architecture, where distinct serotonergic groups mediate different aspects of the satiety program. Among the circuits tested, only *Akh-Gal4* activation showed similar state-matching to *Trhn-Gal4*, suggesting a potential interaction between these two systems in coordinating hunger–satiety transitions. Nonetheless, given that *Akh* inhibition lacked strong effects, this interaction is likely not a simple linear one, with *Trhn* appearing to be the more dominant state regulator. Our experimental paradigm provides an opportunity to address a fundamental question in state control (*2*): to what extent do different neuromodulatory systems have distinct versus overlapping roles in satiety? Future research could use this approach to examine interactions between serotonergic and Akh systems, as well as other feeding-related neuromodulators. More broadly, our findings illustrate how studying circuit manipulations in the context of natural state transitions enhances our understanding of how small populations of neuromodulatory cells generate complex brain states.

The contextualized phenovector approach offers a powerful tool for investigating other complex internal states with tractable behavioral ground truths, including thirst, sleep, stress responses, aggression, and courtship. Comparing manipulation-induced phenovectors to naturalistic state transitions allows researchers to better evaluate whether genetic or physiological perturbations induce fragmentary behavioral changes or authentically drive internal states.

## Materials and Methods

### Fly stocks and husbandry

All flies were maintained on standard cornmeal-based medium (1.25 % w/v agar, 10.5 % w/v dextrose, 10.5 % w/v maize and 2.1 % w/v yeast (*69*). Stocks were kept at room temperature unless otherwise specified. Crosses were performed at 25 °C in a day-night cycling incubator (Mir-253, Sanyo). Unless otherwise specified, adult male flies, aged 5–10 days, were used for behavioral experiments and immunohistochemistry. Experiments with females used mated adult females aged 5–10 days. The *R50H05* driver line is a 1365 base pair fragment of the *SerT* enhancer fused to *Gal4* (*70*). The *Trhn-Gal4* driver line uses a regulatory sequence from the *Trhn* gene (*35*). Other lines included *tsh-Gal80* (*37*), and *αTub84B-frt-Gal80-frt; tsh-LexA*, *8*×*-LexAop2-FLPL/CyO-RFP-tb; UAS-10X-IVS-myr:GFP* (*38*, *71*).

**Table S1.** Genotype abbreviations. Detailed genotype information of fly strains and their short names as referred to in this paper.

| Name | Genotype |
| --- | --- |
| *Trhn>Chr* | *w1118; 20XUAS-IVS-CsChrimson.mVenus/+; Trhn-Gal4/+* |
| *R50H05>Chr* | *w1118; 20XUAS-IVS-CsChrimson.mVenus/+; R50H05-Gal4/+* |
| *Trhn>ACR1* | *w1118; +/+; Trhn-Gal4/UAS-GtACR1.d.EYFP* |
| *R50H05>ACR1* | *w1118; +/+; R50H05-Gal4/UAS-GtACR1.d.EYFP* |
| *Trhn-AND-tsh>Chr* | *w1118, αTub84B-frt-Gal80-frt; tsh-LexA, 8X-LexAop2-FLPL/+; 20XUAS-IVS-CsChrimson.mVenus/Trhn-Gal4* |
| *Trhn-AND-tsh>ACR1* | *w1118, αTub84B-frt-Gal80-frt; tsh-LexA, 8X-LexAop2-FLPL/+; UAS-GtACR1.d.EYFP/Trhn-Gal4* |
| *SXVNC1>Chr* | *w1118; Trhn-p65.AD/+; VT042751-GAL4.DBD/20XUAS-IVS-CsChrimson.mVenus* |
| *SXVNC2>Chr* | *w1118; Trhn-p65.AD/+; R75E01-GAL4.DBD/20XUAS-IVS-CsChrimson.mVenus* |
| *SXVNC1>ACR1* | *w1118; Trhn-p65.AD/+; VT042751-GAL4.DBD/UAS-GtACR1.d.EYFP* |
| *SXVNC2>ACR1* | *w1118; Trhn-p65.AD/+; R75E01-GAL4.DBD/UAS-GtACR1.d.EYFP* |
| *Trhn-NOT-R50H05>Chr* | *w1118; R50H05-lexA:p65/20XUAS-IVS-CsChrimson.mVenus; Trhn-Gal4/LexAop-Gal80* |
| *R50H05-NOT-Trhn>Chr* | *w1118; Trhn-lexA:p65/20XUAS-IVS-CsChrimson.mVenus; R50H05-Gal4/LexAop-Gal80* |
| *Trhn-NOT-tsh>Chr* | *w1118; 20XUAS-IVS-CsChrimson.mVenus/+; tsh-Gal80/Trhn-Gal4* |
| *NPF>Chr* | *w1118; 20XUAS-IVS-CsChrimson.mVenus/NPF-GAL4.1* |
| *NPF>ACR1* | *w1118; NPF-GAL4.1/+; UAS-GtACR1.d.EYFP* |
| *AKH3>Chr* | *w1118; 20XUAS-IVS-CsChrimson.mVenus/+; Akh-gal4.L/+* |
| *AKH2>ACR1* | *w1118; Akh-gal4.L/+; UAS-GtACR1.d.EYFP/+* |
| *Dilp2>Chr* | *w1118; Ilp2-GAL4.R/20XUAS-IVS-CsChrimson.mVenus* |
| *Dilp2>ACR1* | *w1118; Ilp2-GAL4.R/+; UAS-GtACR1.d.EYFP* |

**Table S2.** Fly stocks.

| Name | Source |
| --- | --- |
| 20XUAS-IVS-CsChrimson.mVenus | BDSC #55135 |
| Trhn-Gal4 | Kravitz Lab |
| R50H05-Gal4 | BDSC #38764 |
| UAS-GtACR1.d.EYFP | BDSC #92983 |
| αTub84B-frt-Gal80-frt; tsh-LexA, 8X-LexAop2-FLPL; | Grueber Lab |
| Trhn-p65.AD | BDSC #75650 |
| VT042751-GAL4.DBD | BDSC #74891 |
| R75E01-GAL4.DBD | BDSC #75650 |
| LexAop-Gal80 | Aw Lab |
| R50H05-lexA:p65 | BDSC #52856 |
| tsh-Gal80 | Aw Lab |
| Trhn-lexA:p65 | BDSC #38764 |
| NPF-GAL4.1 | BDSC #25681 |
| Akh-gal4.L (3) | BDSC #25684 |
| Akh-gal4.L (2) | BDSC #25683 |
| Ilp2-GAL4.R | BDSC #37516 |
| UAS-TrhRNAi | VDRC TID105414 |
| UAS-Glut1RNAi | BDSC #40904 |
| UAS-sut2RNAi | BDSC #64861 |

To establish defined nutritional states, flies were assigned to starvation durations (0, 24, or 48 h) as separate between-group cohorts; within-subject tracking of the state transition during refeeding was not performed, as the precise moment of individual state change is uncontrolled and variable.

### Optogenetic control design

For each optogenetic experiment, two genetic controls were used: a “Driver Control” (*Gal4* × *w^1118^*), carrying the *Gal4* driver without the *UAS*-effector, and a “Responder Control” (*UAS* × *w^1118^*), carrying the *UAS*-effector without the *Gal4*. These controls account for any behavioral effects of the *Gal4* or *UAS* transgenes alone. The genotype × light interaction effect size (ΔΔ) was computed relative to the pooled standard deviation across all four groups (lights-off Driver Control, lights-off Responder Control, lights-on Driver Control, lights-on Responder Control), following (*72*).

### Screen for novel VNC 5-HT split-Gal4 lines

We first conducted an *in silico* search of a single-cell transcriptomic sequencing dataset on *Drosophila* VNC gene expression(*39*) using SCope (*73*). SCope was used to search single-cell RNAs from the adult *Drosophila* ventral nerve cord for [*Trhn* ∩ *SerT*] cells (*39*). The search identified a cluster of 67 presumptive serotonergic neurons, which was exported as a .loom file for analysis with [loomxpy](https://github.com/aertslab/LoomXpy). Expression levels for each gene in these 67 cells were computed as Z-scores and ranked. The following criteria were applied to increase signal to noise ratio: (1) Few cells with high Z-scores (fewer than 10 cells with Z-scores above 3 with the maximum Z-score above 5); (2) more than 5 counts were sequenced per cell. By these criteria, 68 genes were selected. The gene list was further trimmed by cross-referencing FlyLight expression first generation Gal-4 database. The genes were then visually inspected in FlyLight (*74*) for proximity/colocation with the *Trhn* neurons in the VNC. Based on this analysis, we drafted a list of candidate genes whose enhancers could be suitable for generating split-*Gal4* lines (*40*), which enable narrow neuronal targeting. For hits from the Flylight screen, GMR fly lines for either *AD* or *DBD* constructs were purchased depending on availability. The driver lines in the table were screened for behavior effects either as straight Gal4 drivers or in combination with either *Trh-p65.AD* (BDSC #70975)*; UAS-Chrimson* or *UAS-Chrimson; Trh-Gal4.DBD* (BDSC #70371).

**Table 1.** Summary of Gal4 and split-Gal4 lines used to make new serotonergic driver lines. Gene names, their derivative promoter fragments, and corresponding Bloomington *Drosophila* Stock Center (BDSC) stock numbers for transgenic lines carrying *Gal4* and/or split-*Gal4* driver lines. The promoters are identified by their Janelia GMR or Vienna Tile codes, and each transgene expresses either whole Gal4; or a transcription-factor moiety with the DNA-binding domain (*DBD*, fused with a dimerization leucine zipper) of *Gal4*; or a zipper fusion with the *Gal4* activation-domain (*AD*).

| Gene | Promoter fragment | BDSC stock numbers |
| --- | --- | --- |
| *Pvf3* | GMR77D01 | 39960 (Gal4) |
| *Ubx* | GMR32B03 | 49886 (Gal4), 69117 (DBD) |
| *Ubx* | VT042751 | 74891 (DBD) |
| **Additional** |  |  |
| *Vsx2* | GMR71G01 | 39599 (Gal4) |
| *cnc* | GMR40C01 | 50078 (Gal4), 69681 (AD) |
| *ab* | GMR75E01 | 39895 (Gal4), 75650 (DBD) |
| *beat-Vc* | GMR93D10 | 40653 (Gal4), 68248 (p65.AD) |

### Experimental preparation of behavioral subjects

Wild-type *w^1118^* flies were collected and kept on normal fly food for 5–8 days before experiment. Prior to the experiment, flies were assigned to starvation durations of 0, 24, or 48 hours; approximately 20 flies per group were transferred onto 2% agarose in water. The 48-hour starved groups were transferred again to a fresh vial of agarose, 24 hours after the start of starvation.

For the transgenic experiments, driver lines were crossed to virgin females of responder lines to produce appropriate test offspring. Genetic controls were produced by crossing the respective parents to *w^1118^* flies. Throughout the optogenetic experiments, all crosses and progenies were maintained in darkness using an aluminum foil vial sleeve. Two to three days prior to the optogenetic experiments, flies were transferred from normal fly food to food containing 0.5 mM all-*trans*-retinal (ATR, R2500, Sigma-Aldrich). In optogenetic experiments, controls and experimental animals were tested in the same sessions. Lights-off conditions were separately tested on different flies to provide an estimate for non-specific light effects.

### Immunohistochemistry

Fly brains were dissected in cold phosphate-buffered saline (PBS) and fixed in 4 % formaldehyde solution (w/v) in PBS for 30 min. The brains were washed three times in PBS before incubation in primary antibodies diluted with PBST [PBS with 0.2 % Triton X-100 (v/v) and 5 % BSA (w/v)] for at least 48 h at 4°C. The brains were then rinsed in PBST before incubation in relevant secondary antibodies diluted in PBST, for another 48 h at 4°C under foil wrap. The antibodies used were: rabbit anti-serotonin (1:1000 dilution; S5545, Sigma-Aldrich), mouse anti-Brp (1:50 dilution; nc82, Developmental Studies Hybridoma Bank), chicken anti-GFP (1:2000 dilution; ab13970, Abcam); goat anti-rabbit 568 nm (1:500 dilution; A-11011, Invitrogen), donkey anti-mouse 647 nm (1:200 dilution; 715-605-151, Jackson ImmunoResearch), and goat anti-chicken 488 nm (1:1000; A-11039, Invitrogen). Finally, brains were washed in PBST three times for 15 min each at room temperature, before mounting onto microscope slides in Vectashield® Vibrance® Antifade Mounting Media (H-1700, Vector Laboratories). Confocal images were acquired under a Zeiss LSM 710 or a Leica TCS SP8 microscope. Images were analyzed and quantified in Fiji (*75*). Maximum-intensity projections were generated from z-stacks spanning the full depth of the tissue.

For 5-HT quantification (**Figure 6C, D**), tissue samples were normalised for variation in antibody penetration and tissue thickness, 5-HT fluorescence intensity in each brain or VNC was expressed as a ratio relative to the mean α-*Brp* (nc82) signal measured in the same preparation (5-HT intensity / *Brp* intensity). Regions of interest were detected with a threshold in FIJI, and mean fluorescence intensities were extracted in this ROI for both channels. This normalisation corrects for global intensity differences between preparations while preserving the relative 5-HT signal. All images within a comparison (e.g., fed versus 48-h starved) were processed identically.

### Espresso: Hardware assembly

A custom data acquisition system, Espresso, was used to collect feeding data. In brief, flies were housed in custom-made plastic chips, each containing five feeding chambers. Two feed tubes, each holding a capillary, supplied food to each chamber. Each experiment was conducted on a single panel holding six Espresso chips. The entire panel was imaged at 30 frames per second (fps) using a videographic assembly consisting of a 4.4-11mm High Resolution Varifocal Lens (Edmund Optics) and a CCD camera (CM3-U3-13S2C-CS, Point Grey) connected to a data acquisition computer. The assay was backlit with a custom infrared LED (850 nm, OEM) panel so that the flies and IR dye and fly bodies appear black in images acquired.

### Espresso: Light administration for optogenetic experiments

Red-orange (617 nm, SP-05-E4, Luxeon Star LEDs, Quadica Developments Inc.) and green (530 nm, SP-05-G4, Luxeon Star LEDs, Quadica Developments Inc.) LEDs were mounted onto heatsinks (N50-25B, Luxeon Star LEDs, Quadica Developments Inc.) to achieve an illumination of approximately 20 µW/mm^2^ for red light and 15 µW/mm^2^ for green light. Light was administered continuously throughout the 2-hour experiment duration. The onset and offset of illumination were controlled by software (see below) via an assembly of breakout circuits consisting of I/O hub, BuckPuck and analog output modules (HUB0000, 3021-D-E-700 and OUT1001, Phidgets Inc.). Light-intensity measurements were performed with a photodiode (Thorlabs S130C) connected to power and energy-meter console (Thorlabs PM100D) to measure light intensity in a dark room, as previously described (*76*).

### Preparation for feeding experiments

Glass capillaries (1B100-3, 1/0.58mm OD/ID, WPI) were loaded with ∼1–2 mm of IR-Dye (*27*) followed by 30 mm of liquid food via capillary action. The capillaries were inserted into food channels of custom-made Espresso chips and secured with silicone adaptors. Chips loaded with food capillaries were then inserted into holders on the Espresso panel. Single flies were briefly anesthetized in a chilled empty food vial on ice and then transferred to the feeding chambers with a small paint brush, which were sealed with magnetic coverslips. Once the flies regained normal activity, the panel was positioned in the acquisition setup (**Figure 1A**) housed in an incubator (Mir-154, Panasonic). After the experiments, the capillaries were washed with 1% Tergazyme (Alconox Inc.) solution followed by deionized water in a vacuum-assisted wash chamber and air dried.

### Behavior tracking software

Video images were processed with the custom software package CRITTA (Control and Real-time Imaging Tool for Tracking Animals) written in LabView (*77*) in combination with Vision Acquisition and Vision Runtime software (2013, National Instruments Corp) to track both liquid food levels and fly positions.

### Tracking of food levels and fly positions

A dark band formed by the IR-absorbing dye marked the food meniscus in each capillary, allowing its position to be tracked over time to report the food level. The centroid of each individual fly was tracked in two-dimensional space, with X and Y coordinates recorded. Fly positions and food levels were tracked at 30 samples/second throughout the experiments.

### Detection of feed events

A virtual feed port ∼2mm from the end of the feeding capillary was demarcated in the software at the bottom end of each capillary, acting as a switch to detect feeding events. When feed ports were vacant, the reduction in liquid levels within the capillaries was used to estimate the rate of evaporation. Feeding events were recorded when a fly entered the feed port and a reduction in the food level (beyond the rate of evaporation) co-occurred. The reduction was considered significant and logged as feed using a moving average convergence/divergence (MACD) algorithm applied to the liquid level. The event ended when the food level stabilized and/or when the fly left the feed port. Real-time evaporation levels were subtracted from feed volumes recorded.

### Behavior analysis software

#### Data tabulation

Fly locomotion data and capillary levels were collected from all 30 feeding chambers simultaneously and recorded in real time. The timestamp, volume, duration of the feed and evaporation rate at the time of the feed were computed at the time of detection with a custom tracking software CRITTA. Fly X- and Y-positions and speed were recorded for each frame.

#### Definition of feed metrics

From continuous tracking data, further processed metrics were calculated in an analysis package (`esploco`), written in Python. Feeding metrics included:

1. Meal duration (in seconds)*
2. Meal size (in nL)*
3. Feed speed (rate of liquid consumption, in nL per second)*
4. Feed volume (total volume of the liquid food consumed during the assayed period, in nL)
5. Feed count (the total number of meals consumed during the assayed period)
6. Feed duration (total duration the fly spends feeding during the assayed period)

*\*Based on an individual meal*

Locomotive metrics included:

7. Latency (time spent before the first feed was initiated, in seconds)
8. Prefeed speed (speed during the 120 s before a feed, in mm/s)
9. Postfeed speed (speed during the 120 s after a feed, in mm/s)
10. Duringfeed speed (speed during the feed throughout the feed event)
11. Duringfeed speed ratio (ratio between duringfeed speed and prefeed speed)
12. Peri-feed speed ratio (ratio between postfeed speed and prefeed speed)
13. Speed (average speed in mm/s during the assayed period).
14. Elevation (y-position of the fly during the assayed period).
15. Food port occupancy (percent time a fly spends in a feeding port containing food).
16. Control port occupancy (percent time a fly spends in the feeding port where food is unavailable).
17. Falls (number of falls during the assayed period).

### Data visualization and statistics

Data were visualized with custom Python notebooks using the DABEST, espresso, esploco, numpy, scipy, pandas, seaborn, and matplotlib libraries (**Table S3**).

### Effect-size estimation computation and visualization

Effect sizes for comparisons were calculated as mean differences (Δ), Hedges’ *g*, ΔΔ, or Δ*g*, where appropriate. All estimation plots share common visualization elements: in the observed-data axes, individual values appear as dots in swarm plots, means and standard deviations are shown as vertical gapped lines (gaps indicating means); in the inference axes, bootstrapped differences between groups appear as half-violin distributions with black dots (mean effect sizes) and vertical bars (95% confidence intervals, which range between the mean ± the margin of error). For standardized effect sizes, g = 0.2 was considered a small effect size, 0.5 a medium effect size, and 0.8 a large effect size (*78*). In text, the mean differences and their 95% confidence intervals (95CIs) are written as: “meanΔ [95CI lower-bound, upper-bound].” Three types of estimation plots were used based on the different experimental designs, as follows.

Gardner-Altman plots (*79*) display two-group comparisons. Simple two-group estimation was performed as previously described (*80*). Briefly, the left panel shows individual data points as a swarm with their means (measures of central tendency) and standard deviations (measures of dispersion), while the right panel (the contrast or inference panel) shows the mean-difference estimate between the two groups as a point estimate, 95CI, and a bootstrapped distribution (visualization of estimate precision).

Cumming plots (*81*) accommodate multiple-group comparisons by showing observed-values swarm plots in the top panel with their means and standard deviations. Below these, contrast plots show bootstrapped difference distributions between specified groups, aligned with their corresponding observed values above.

Delta-delta (ΔΔ) plots visualize 2 × 2 experimental designs where two control groups (driver and responder flies) are pooled and used to calculate differences with the data from the test animals, across two conditions. This design was used to isolate the [light × genotype] effect, i.e. properly control for the two-way optogenetic effect. As in a Cumming plot, the top panel shows the observed data across all groups with their means and standard deviations. The bottom panel displays both between-group contrasts and the delta-delta effect (the difference between differences). For standardized two-way effect sizes, Δ*g* for a 2 × 2 experimental design with groups [A1, A2, B1, B2] was computed as follows:

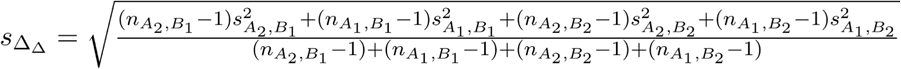

where *s* is the standard deviation and *n* is the sample size.

In a typical power calculation, the sample size of 45 per group would provide a power >0.8, where α = 0.05 and an effect size of ≥ 0.6 standard-deviation. However, in effect size estimation no alpha thresholds were set and no significance tests were conducted; two-tailed *P*-values were calculated in `SciPy` and reported *pro forma* only. The precision of estimates is reflected in the width of confidence intervals, where narrower intervals indicate more precise estimates. When 95% confidence intervals do not overlap zero, a null effect can be considered implausible, although this is not definitive (*81*).

### Construction of state-transition vectors and phenovectors

For each metric, Hedges’ *g* was computed for the difference between 0h starved and 24h starved groups, as well as between 0-h starved and 48-h starved groups. 95CI for Hedges’ g was computed by bootstrap resampling. Vectors of 17 dimensions were constructed for each of the following conditions 24h − 0h (24h starved), 48h − 0h (48h starved), 0h − 24h (24h reversed) and 0h − 48h (48h reversed) to represent the behavior changes of natural state transitions. To represent phenotypes in the same dimensionality, Δ*g* effects were computed for each metric in a 2 × 2 arrangement of 4 groups: [(test light on − test light off) − (control light on − control light off)] as mentioned in the section above for each of the 17 metrics. Such *phenovectors* were assembled to represent the multi-attribute phenotypes of each neurogenetic intervention.

### Phenovector analysis design rationale

The state-transition vector approach was designed around three principles. First, by using Hedges’ g to standardize effect sizes across metrics measured on different scales, we placed all behavioral dimensions on a common axis, allowing direct comparison of interventions with widely differing effect magnitudes. Second, by computing effect sizes against a reference group (fed, 0-h starvation), we constructed vectors that capture the direction and magnitude of behavioral change, not just the endpoint value. Third, by comparing optogenetic phenovectors to the independently-derived starvation reference vectors, we tested whether each manipulation moves the animal along the same behavioral axis as natural state changes—a conceptually distinct test from asking whether a manipulation changes feeding volume alone.

Linear regression and Pearson’s r were used as the primary alignment metric for its interpretability, though Spearman rank correlation and cosine distance analyses (**Figure S9 B, C**) confirm the key findings are robust to this choice. The vector approach is sensitive to the number and choice of metrics: adding or removing metrics can alter apparent similarity, as detailed in the sensitivity analyses (**Figure S9A**).

### Hierarchical clustering

Natural state transition vectors and phenovectors were computed and tabulated for each metric. The means of the bootstrap distribution was used to determine the hierarchical arrangement of each condition using the Python package **scipy.cluster.hierarchy**, using ‘average’ as the method setting and ‘correlation’ as the metric setting. To represent the precision of each effect size, each bootstrap distribution is color-coded by effect size and was visualized by ranking the colors in each cell in a spiral, from the middle of the cell outward. Linear regression was done with SciPy, to calculate the Pearson correlation coefficient (*r*), and slope (*β*).

### Definition of behavioral states

We define the natural hunger state as the coordinated profile of behavioral changes that reliably accompanies graded starvation durations (0, 24 and 48 h) in w^1118^ flies as discrete samples from a continuous hunger-satiety state space. An “authentic hunger-like state” (AHLS) refers to an optogenetically induced behavioral profile whose phenovector aligns with the natural starvation-transition vector (see section above). Since our definitions are grounded in the metrics collected by Espresso, we acknowledge the possibility of dissociation of our “AHLS” from unmeasured dimensions of natural behavior.

## Supporting information

Supplementary Table 3 Custom software and availability

Supplementary Table 4 Genotype and effect sizes of experiments

## Data and code availability

Software packages for analysis are provided as documented in **Table S3**.

Raw and processed data are available at the Zenodo data repository (DOI:10.5281/zenodo.205605821).

## Acknowledgements

The authors would like to thank Drs. Edward Kravitz (Harvard Medical School) and Olga Alekseyenko (Harvard Medical School) for the *Trhn-Gal4* line; Dr. Wes Grueber (Columbia University) for *Tub-frt-Gal80-frt; tsh-LexA*, *8*×*-LexAop2-FLPL/CyO-RFP-tb; UAS-10X-IVS-myr:GFP* flies; and Dr. Sherry Aw (Institute of Molecular and Cell Biology) for *tsh-Gal80* flies. The authors also thank Zoe Ang (National University of Singapore), Farhana Anisha (National University of Singapore), and Hazimah Yusoff (Institute of Molecular and Cell Biology) for technical assistance, Keshmarathy Sacadevan at SingHealth Advanced Bioimaging and Dr. Eloise Xiaoxiao Ma at IMCB Central Imaging Facility for assistance with confocal microscopy, and Dr. Jessica Tamanini (Insight Editing London) for critical reviews of the manuscript before submission.

## Author Contributions

**Conceptualization**: SX, XY, ACC; **Experiment design**: SX, XY, ACC; **Methodology**: SX, XY, JCS, DC, ZW; **Software**: SX, JH (Python), JCS (CRITTA, LabView); **Data Analysis:** SX, XY (Python); **Investigation**: SX (feeding assays, genetics, brain dissection, immunohistochemistry, and microscopy), XY (feeding assays); **Writing – Original Draft:** SX; **Writing – Revision:** SX, XY, ACC; **Visualization:** SX, XY, ACC, JH; **Supervision:** ACC; **Project Administration:** ACC; **Funding Acquisition:** ACC.

