## Supplementary Table 3 Custom software and availability for "Contextualized phenovectors reveal a Drosophila serotonergic circuit controlling satiety"

| Software | Platform | Version | Link |
|---|---|---|---|
| esploco | Python | 24.11.12 | https://github.com/sangyu/esploco |
| espresso | Python | 0.7.3 | https://github.com/ACCLAB/espresso |
| DABEST-Python | Python | 2025.10.20 | https://github.com/ACCLAB/DABEST-python |
| Figure Generation | Python Notebooks | 2026.06.08 | https://github.com/sangyu/Ethomics5HT |
