## Supplementary Table 4 Genotype and effect sizes of experiments for "Contextualized phenovectors reveal a Drosophila serotonergic circuit controlling satiety"

### Figure 1

#### Figure 1H Volume (nL)

| Genotype | control | test | control_N | test_N | effect_size | difference | bca_low | bca_high | pvalue_mann_whitney |
|---|---|---|---|---|---|---|---|---|---|
| w1118 | 0 | 24 | 150 | 120 | mean difference | 85.74 | 59.26 | 114.76 | 0.0 |
| w1118 | 0 | 48 | 150 | 90 | mean difference | 269.88 | 226.21 | 316.02 | 0.0 |

#### Figure 1N Food Port Occupancy

| Genotype | control | test | control_N | test_N | effect_size | difference | bca_low | bca_high | pvalue_mann_whitney |
|---|---|---|---|---|---|---|---|---|---|
| w1118 | 0 | 24 | 150 | 120 | mean difference | 0.33 | -0.16 | 2.27 | 0.04 |
| w1118 | 0 | 48 | 150 | 90 | mean difference | 0.81 | 0.44 | 1.38 | 0.0 |

#### Figure 1O Ctrl Port Occupancy

| Genotype | control | test | control_N | test_N | effect_size | difference | bca_low | bca_high | pvalue_mann_whitney |
|---|---|---|---|---|---|---|---|---|---|
| w1118 | 0 | 24 | 150 | 120 | mean difference | -0.37 | -0.54 | -0.21 | 0.04 |
| w1118 | 0 | 48 | 150 | 90 | mean difference | -0.36 | -0.53 | -0.21 | 0.19 |

#### Figure 1N Food Port Occupancy

| Genotype | control | test | control_N | test_N | effect_size | difference | bca_low | bca_high | pvalue_mann_whitney |
|---|---|---|---|---|---|---|---|---|---|
| w1118 | 0 | 24 | 150 | 120 | mean difference | 0.33 | -0.16 | 2.27 | 0.04 |
| w1118 | 0 | 48 | 150 | 90 | mean difference | 0.81 | 0.44 | 1.38 | 0.0 |

#### Figure 1O Ctrl Port Occupancy

| Genotype | control | test | control_N | test_N | effect_size | difference | bca_low | bca_high | pvalue_mann_whitney |
|---|---|---|---|---|---|---|---|---|---|
| w1118 | 0 | 24 | 150 | 120 | mean difference | -0.37 | -0.54 | -0.21 | 0.04 |
| w1118 | 0 | 48 | 150 | 90 | mean difference | -0.36 | -0.53 | -0.21 | 0.19 |

**Figure 2**

| | |
|---|---|
| **Figure 2A Trhn>Chr** | Figure2A_TrhnCsChrimson.lif |
| **Figure 2B R50H05>Chr** | Figure2B_R50H05CsChrimson.lif |

**Figure 2F Trhn>Chr**

| control genotype | test genotype | control condition | test condition | control_N | test_N | effect_size | difference | bca_low | bca_high | pvalue_permutation |
|---|---|---|---|---|---|---|---|---|---|---|
| w/Trhn-Gal4, w/UAS-Chrimson | Trhn-Gal4>UAS-Chrimson | Ctrl Red Light Off | Test Red Light Off | 81 | 39 | mean difference | 31.66 | -19.14 | 75.7 | 0.21 |
| w/Trhn-Gal4, w/UAS-Chrimson | Trhn-Gal4>UAS-Chrimson | Ctrl Red Light On | Test Red Light On | 121 | 59 | mean difference | -169.27 | -206.01 | -135.86 | 0.0 |
| | | Test Red Light Off - Ctrl Red Light Off | Test Red Light On - Ctrl Red Light On | 5000 | 5000 | Delta Delta | -200.93 | -259.88 | -139.77 | 0.0 |

**Figure 2G Trhn>ACR1**

| control genotype | test genotype | control condition | test condition | control_N | test_N | effect_size | difference | bca_low | bca_high | pvalue_permutation |
|---|---|---|---|---|---|---|---|---|---|---|
| w/Trhn-Gal4, w/UAS-ACR1 | Trhn-Gal4>UAS-ACR1 | Ctrl Green Light Off | Test Green Light Off | 81 | 39 | mean difference | 26.18 | -10.66 | 68.03 | 0.15 |
| w/Trhn-Gal4, w/UAS-ACR1 | Trhn-Gal4>UAS-ACR1 | Ctrl Green Light On | Test Green Light On | 80 | 40 | mean difference | 191.7 | 127.47 | 283.71 | 0.0 |
| | | Test Green Light Off - Ctrl Green Light Off | Test Green Light On - Ctrl Green Light On | 5000 | 5000 | Delta Delta | 165.51 | 85.66 | 252.93 | 0.0 |

**Figure 2J R50H05>Chr**

| control genotype | test genotype | control condition | test condition | control_N | test_N | effect_size | difference | bca_low | bca_high | pvalue_permutation |
|---|---|---|---|---|---|---|---|---|---|---|
| w/R50H05-Gal4, w/UAS-Chrimson | R50H05-Gal4>UAS-Chrimson | Ctrl Red Light Off | Test Red Light Off | 120 | 60 | mean difference | 1.43 | -22.79 | 20.83 | 0.91 |
| w/R50H05-Gal4, w/UAS-Chrimson | R50H05-Gal4>UAS-Chrimson | Ctrl Red Light On | Test Red Light On | 140 | 70 | mean difference | 249.13 | 211.21 | 290.46 | 0.0 |
| | | Test Red Light Off - Ctrl Red Light Off | Test Red Light On - Ctrl Red Light On | 5000 | 5000 | Delta Delta | 247.7 | 201.75 | 294.51 | 0.0 |

**Figure 2K R50H05>ACR1**

| control genotype | test genotype | control condition | test condition | control_N | test_N | effect_size | difference | bca_low | bca_high | pvalue_permutation |
|---|---|---|---|---|---|---|---|---|---|---|
| w/R50H05-Gal4, w/UAS-ACR1 | R50H05-Gal4>UAS-ACR1 | Ctrl Green Light Off | Test Green Light Off | 40 | 20 | mean difference | 51.41 | -38.45 | 133.9 | 0.25 |
| w/R50H05-Gal4, w/UAS-ACR1 | R50H05-Gal4>UAS-ACR1 | Ctrl Green Light On | Test Green Light On | 77 | 38 | mean difference | 23.1 | -60.33 | 109.59 | 0.63 |
| | | Test Green Light Off - Ctrl Green Light Off | Test Green Light On - Ctrl Green Light On | 5000 | 5000 | Delta Delta | -28.31 | -150.33 | 95.23 | 0.66 |

**Figure 2L Volume Forest**

| | Trhn>Chr | Trhn>ACR1 | R50H05>Chr | R50H05>ACR1 |
|---|---|---|---|---|
| mean | -200.93 | 165.51 | 247.7 | -28.31 |
| bca_low | -259.88 | 85.66 | 201.75 | -150.33 |
| bca_high | -139.77 | 252.93 | 294.51 | 95.23 |

**Figure 2O Count Forest**

| | Trhn>Chr | Trhn>ACR1 | R50H05>Chr | R50H05>ACR1 |
|---|---|---|---|---|
| mean | 4.86 | -2.98 | 15.95 | -0.12 |
| bca_low | -0.27 | -10.7 | 11.57 | -5.18 |
| bca_high | 10.33 | 4.32 | 20.69 | 4.73 |

**Figure 2N Meal Size Forest**

| | Trhn>Chr | Trhn>ACR1 | R50H05>Chr | R50H05>ACR1 |
|---|---|---|---|---|
| mean | -75.4 | 47.15 | 8.62 | 5.68 |
| bca_low | -106.19 | 30.07 | -0.43 | -45.48 |
| bca_high | -46.6 | 66.77 | 16.17 | 53.96 |

**Figure 2M Duration Forest**

| | Trhn>Chr | Trhn>ACR1 | R50H05>Chr | R50H05>ACR1 |
|---|---|---|---|---|
| mean | -0.31 | 0.57 | 1.38 | -0.08 |
| bca_low | -0.7 | 0.13 | 1.01 | -0.63 |
| bca_high | 0.12 | 1.03 | 1.75 | 0.45 |

### Figure S1

#### Figure S1A Trhn>Chrimson Female

| control genotype | test genotype | control condition | test condition | control_N | test_N | effect_size | difference | bca_low | bca_high | pvalue_permutation |
|---|---|---|---|---|---|---|---|---|---|---|
| w/Trhn-Gal4, w/UAS-Chrimson | Trhn-Gal4>UAS-Chrimson | Ctrl Red Light Off | Test Red Light Off | 35 | 25 | mean difference | 73.57 | -6.48 | 193.43 | 0.1 |
| w/Trhn-Gal4, w/UAS-Chrimson | Trhn-Gal4>UAS-Chrimson | Ctrl Red Light On | Test Red Light On | 18 | 12 | mean difference | -320.54 | -713.62 | -146.25 | 0.03 |
| | | Test Red Light Off - Ctrl Red Light Off | Test Red Light On - Ctrl Red Light On | 5000 | 5000 | Delta Delta | -394.11 | -714.71 | -179.0 | 0.01 |

#### Figure S1B Trhn>ACR1 Female

| control genotype | test genotype | control condition | test condition | control_N | test_N | effect_size | difference | bca_low | bca_high | pvalue_permutation |
|---|---|---|---|---|---|---|---|---|---|---|
| w/Trhn-Gal4, w/UAS-ACR1 | Trhn-Gal4>UAS-ACR1 | Ctrl Green Light Off | Test Green Light Off | 32 | 28 | mean difference | 13.79 | -38.52 | 62.63 | 0.6 |
| w/Trhn-Gal4, w/UAS-ACR1 | Trhn-Gal4>UAS-ACR1 | Ctrl Green Light On | Test Green Light On | 36 | 24 | mean difference | 102.58 | 36.24 | 177.48 | 0.0 |
| | | Test Green Light Off - Ctrl Green Light Off | Test Green Light On - Ctrl Green Light On | 5000 | 5000 | Delta Delta | 88.79 | 4.35 | 177.4 | 0.05 |

#### Figure S1C R50H05>Chr Female

| control genotype | test genotype | control condition | test condition | control_N | test_N | effect_size | difference | bca_low | bca_high | pvalue_permutation |
|---|---|---|---|---|---|---|---|---|---|---|
| w/R50H05-Gal4, w/UAS-Chrimson | R50H05-Gal4>UAS-Chrimson | Ctrl Red Light Off | Test Red Light Off | 17 | 13 | mean difference | 15.26 | -62.76 | 110.06 | 0.73 |
| w/R50H05-Gal4, w/UAS-Chrimson | R50H05-Gal4>UAS-Chrimson | Ctrl Red Light On | Test Red Light On | 33 | 27 | mean difference | 108.9 | 27.52 | 198.01 | 0.01 |
| | | Test Red Light Off - Ctrl Red Light Off | Test Red Light On - Ctrl Red Light On | 5000 | 5000 | Delta Delta | 93.64 | -22.78 | 209.94 | 0.13 |

### Figure S2

#### Figure S2A Trhn>Chr Separate controls

| control genotype | test genotype | control condition | test condition | control_N | test_N | effect_size | difference | bca_low | bca_high | pvalue_permutation |
|---|---|---|---|---|---|---|---|---|---|---|
| Chr/w | Trhn/w | Chr/w Red Light Off | Trhn/w Red Light Off | 40 | 41 | mean difference | 90.73134207 | 39.81461829 | 144.2835982 | 0.001 |
| Chr/w | Trhn>Chr | Chr/w Red Light Off | Trhn>Chr Red Light Off | 40 | 39 | mean difference | 77.58766603 | 18.22628269 | 131.0873019 | 0.0086 |
| Chr/w | Trhn/w | Chr/w Red Light On | Trhn/w Red Light On | 60 | 61 | mean difference | 131.8944514 | 77.58267732 | 194.4193981 | 0 |
| Chr/w | Trhn>Chr | Chr/w Red Light On | Trhn>Chr Red Light On | 60 | 59 | mean difference | -102.7745175 | -145.7865548 | -64.16819237 | 0 |

#### Figure S2B Trhn>ACR1 Separate controls

| control genotype | test genotype | control condition | test condition | control_N | test_N | effect_size | difference | bca_low | bca_high | pvalue_permutation |
|---|---|---|---|---|---|---|---|---|---|---|
| ACR1/w | Trhn/w | ACR1/w Green Light Off | Trhn/w Green Light Off | 41 | 40 | mean difference | 90.92534634 | 60.3107061 | 123.4856963 | 0 |
| ACR1/w | Trhn>ACR1 | ACR1/w Green Light Off | Trhn>ACR1 Green Light Off | 41 | 39 | mean difference | 71.08463352 | 34.41378924 | 112.4321401 | 0.0006 |
| ACR1/w | Trhn/w | ACR1/w Green Light On | Trhn/w Green Light On | 40 | 40 | mean difference | 70.697475 | 34.053575 | 113.747425 | 0.0004 |
| ACR1/w | Trhn>ACR1 | ACR1/w Green Light On | Trhn>ACR1 Green Light On | 40 | 40 | mean difference | 227.044475 | 163.245675 | 315.310775 | 0 |

#### Figure S2C R50H05>Chr Separate controls

| control genotype | test genotype | control condition | test condition | control_N | test_N | effect_size | difference | bca_low | bca_high | pvalue_permutation |
|---|---|---|---|---|---|---|---|---|---|---|
| Chr/w | R50H05/w | Chr/w Red Light Off | R50H05/w Red Light Off | 60 | 60 | mean difference | 4.574466667 | -37.8156 | 25.8089 | 0.8092 |
| Chr/w | R50H05>Chr | Chr/w Red Light Off | R50H05>Chr Red Light Off | 60 | 60 | mean difference | 3.7159 | -39.05115 | 26.24781667 | 0.8362 |
| Chr/w | R50H05/w | Chr/w Red Light On | R50H05/w Red Light On | 70 | 70 | mean difference | -7.291657143 | -27.54941429 | 14.56835714 | 0.4964 |
| Chr/w | R50H05>Chr | Chr/w Red Light On | R50H05>Chr Red Light On | 70 | 70 | mean difference | 245.4875429 | 205.0343 | 288.4537143 | 0 |

#### Figure S2D R50H05>ACR1 Separate controls

| control genotype | test genotype | control condition | test condition | control_N | test_N | effect_size | difference | bca_low | bca_high | pvalue_permutation |
|---|---|---|---|---|---|---|---|---|---|---|
| ACR1/w | R50H05/w | ACR1/w Green Light Off | R50H05/w Green Light Off | 20 | 20 | mean difference | 1.6484 | -94.362 | 93.6898 | 0.9708 |
| ACR1/w | R50H05>ACR1 | ACR1/w Green Light Off | R50H05>ACR1 Green Light Off | 20 | 20 | mean difference | 52.23605 | -40.41945 | 142.91685 | 0.271 |
| ACR1/w | R50H05/w | ACR1/w Green Light On | R50H05/w Green Light On | 38 | 39 | mean difference | 6.361115385 | -111.1295034 | 114.1459879 | 0.914 |
| ACR1/w | R50H05>ACR1 | ACR1/w Green Light On | R50H05>ACR1 Green Light On | 38 | 38 | mean difference | 26.32434211 | -89.15855263 | 130.9091053 | 0.636 |

### Figure 3

#### Figure 3E Elevation Forest

| | Trhn>Chr | Trhn>ACR1 | R50H05>Chr | R50H05>ACR1 |
|---|---|---|---|---|
| mean | -3.11 | 1.54 | -1.91 | 0.8 |
| bca_low | -4.32 | 0.45 | -3.05 | -0.94 |
| bca_high | -1.91 | 2.72 | -0.75 | 2.76 |

#### Figure 3F Food Port Occupancy

| | Trhn>Chr | Trhn>ACR1 | R50H05>Chr | R50H05>ACR1 |
|---|---|---|---|---|
| mean | -0.46 | 0.89 | -0.29 | -0.42 |
| bca_low | -2.92 | -0.53 | -2.36 | -1.98 |
| bca_high | 1.74 | 2.25 | 1.18 | 1.40 |

#### Figure 3F Latency Forest

| | Trhn>Chr | Trhn>ACR1 | R50H05>Chr | R50H05>ACR1 |
|---|---|---|---|---|
| mean | 50.21 | -34.38 | -59.03 | -0.27 |
| bca_low | 29.96 | -59.09 | -76.46 | -31.69 |
| bca_high | 70.59 | -9.75 | -41.34 | 29.82 |

#### Figure 3H Walking Speed Forest

| | Trhn>Chr | Trhn>ACR1 | R50H05>Chr | R50H05>ACR1 |
|---|---|---|---|---|
| mean | 0.48 | -0.38 | 1.18 | 0.02 |
| bca_low | 0.13 | -1.04 | 0.92 | -0.24 |
| bca_high | 0.84 | 0.28 | 1.44 | 0.34 |

#### Figure 3I Perifeed Speed Ratio Forest

| | Trhn>Chr | Trhn>ACR1 | R50H05>Chr | R50H05>ACR1 |
|---|---|---|---|---|
| mean | 0.41 | -0.14 | -0.08 | -0.13 |
| bca_low | 0.18 | -0.35 | -0.46 | -0.39 |
| bca_high | 0.67 | 0.06 | 0.24 | 0.1 |

**Figure S3**

**Figure S3A Early v Late Speed Controls**

| control genotype | test genotype | control condition | test condition | control_N | test_N | effect_size | difference | bca_low | bca_high | pvalue_permutation |
|---|---|---|---|---|---|---|---|---|---|---|
| Early | Late | Red Light Off E | Red Light On E | 81 | 121 | mean difference | -0.3481303006 | -0.7443259357 | 0.01486024621 | 0.06 |
| Early | Late | Red Light Off L | Red Light On L | 81 | 121 | mean difference | -0.2085230538 | -0.4364549105 | -0.03281634427 | 0.0288 |
| | | Red Light On E - Red Light Off E | Red Light On L - Red Light Off L | 5000 | 5000 | Delta Delta | 0.1396072468 | -0.2780001121 | 0.5712456256 | 0.281 |

**Figure S3B Early v Late Speed Test**

| control genotype | test genotype | control condition | test condition | control_N | test_N | effect_size | difference | bca_low | bca_high | pvalue_permutation |
|---|---|---|---|---|---|---|---|---|---|---|
| Early | Late | Red Light Off E | Red Light On E | 39 | 59 | mean difference | -0.1330632666 | -0.439784123 | 0.04590789205 | 0.2216 |
| Early | Late | Red Light Off L | Red Light On L | 39 | 59 | mean difference | 0.5430105788 | 0.2410194784 | 0.8048071355 | 0.0008 |
| | | Red Light On E - Red Light Off E | Red Light On L - Red Light Off L | 5000 | 5000 | Delta Delta | 0.6760738454 | 0.3304783326 | 1.045834282 | 0 |

**Figure S3C Early v Late Volume Controls**

| control genotype | test genotype | control condition | test condition | control_N | test_N | effect_size | difference | bca_low | bca_high | pvalue_permutation |
|---|---|---|---|---|---|---|---|---|---|---|
| Early | Late | Red Light Off E | Red Light On E | 81 | 121 | mean difference | 17.04593348 | -17.21209346 | 53.55251127 | 0.371 |
| Early | Late | Red Light Off L | Red Light On L | 81 | 121 | mean difference | -13.19393878 | -32.89111489 | 6.996187532 | 0.217 |
| | | Red Light On E - Red Light Off E | Red Light On L - Red Light Off L | 5000 | 5000 | Delta Delta | -30.23987226 | -72.60032374 | 8.903743802 | 0.1502 |

**Figure S3D Early v Late Volume Test**

| control genotype | test genotype | control condition | test condition | control_N | test_N | effect_size | difference | bca_low | bca_high | pvalue_permutation |
|---|---|---|---|---|---|---|---|---|---|---|
| Early | Late | Red Light Off E | Red Light On E | 39 | 59 | mean difference | -140.1925641 | -176.222598 | -102.0879235 | 0 |
| Early | Late | Red Light Off L | Red Light On L | 39 | 59 | mean difference | -56.88412777 | -91.5528731 | -25.09894481 | 0.0006 |
| | | Red Light On E - Red Light Off E | Red Light On L - Red Light Off L | 5000 | 5000 | Delta Delta | 83.30843633 | 33.46598479 | 134.4340348 | 0.0012 |

**Figure S3E Falls**

| | Trhn>Chr | Trhn>ACR1 | R50H05>Chr | R50H05>ACR1 |
|---|---|---|---|---|
| mean | -10.01 | -2.43 | -1.29 | -3.11 |
| bca_low | -40.33 | -45.45 | -19.99 | -17.2 |
| bca_high | 19.19 | 47.1 | 18.61 | 9.91 |

**Figure S3F R50H05>Chr xy Variance**

| control genotype | test genotype | control condition | test condition | control_N | test_N | effect_size | difference | bca_low | bca_high | pvalue_permutation |
|---|---|---|---|---|---|---|---|---|---|---|
| w/R50H05-Gal4, w/UAS-Chrimson | R50H05-Gal4>UAS-Chrimson | Ctrl Red Light Off | Test Red Light Off | 120 | 60 | mean difference | -0.24 | -0.75 | 0.27 | 0.37 |
| w/R50H05-Gal4, w/UAS-Chrimson | R50H05-Gal4>UAS-Chrimson | Ctrl Red Light On | Test Red Light On | 140 | 70 | mean difference | 1.77 | 1.39 | 2.16 | 0.0 |
| | | Test Red Light Off - Ctrl Red Light Off | Test Red Light On - Ctrl Red Light On | 5000 | 5000 | Delta Delta | 2.01 | 1.38 | 2.64 | 0.0 |

**Figure S3G Speed (mm/s)**

| control | test | control_N | test_N | effect_size | difference | bca_low | bca_high | pvalue_mann_whitney |
|---|---|---|---|---|---|---|---|---|
| 0 | 24 | 150 | 120 | mean difference | -0.21 | -0.33 | -0.12 | 0.12 |
| 0 | 48 | 150 | 90 | mean difference | -0.26 | -0.37 | -0.16 | 0.06 |

**Figure S3I R50H05>Chr Walking Speed No Food**

| control genotype | test genotype | control condition | test condition | control_N | test_N | effect_size | difference | bca_low | bca_high | pvalue_permutation |
|---|---|---|---|---|---|---|---|---|---|---|
| w/R50H05-Gal4, w/UAS-Chrimson | R50H05-Gal4>UAS-Chrimson | Ctrl 0 | Test 0 | 40 | 20 | mean difference | 1.2 | 0.94 | 1.54 | 0.0 |
| w/R50H05-Gal4, w/UAS-Chrimson | R50H05-Gal4>UAS-Chrimson | Ctrl 24 | Test 24 | 39 | 20 | mean difference | 1.23 | 0.82 | 1.63 | 0.0 |
| | | Test 0 - Ctrl 0 | Test 24 - Ctrl 24 | 5000 | 5000 | Delta Delta | 0.03 | -0.48 | 0.51 | 0.93 |

**Figure 4**

**Figure 4 Forest**

| | Trhn-NOT-R50H05>Chr | R50H05-NOT-Trhn>Chr | Trhn-NOT-tsh>Chr | Trhn-AND-tsh>Chr | SXVNC1>Chr | SXVNC2>Chr | Trhn-AND-tsh>ACR1 | SXVNC1>ACR1 | SXVNC2>ACR1 |
|---|---|---|---|---|---|---|---|---|---|
| mean | -117.43 | 24.03 | -36.33 | -227.97 | -250.53 | -181.38 | 33.52 | 50.58 | 35.88 |
| bca_low | -152.53 | -25.42 | -196.49 | -332.02 | -375.63 | -254.25 | -47.14 | -20.04 | -36.43 |
| bca_high | -78.92 | 66.68 | 126.91 | -128.32 | -126.98 | -106.32 | 114.89 | 127.01 | 111.12 |

### Figure S4

| Figure Panel | Genotype | Confocal File |
|---|---|---|
| Figure S4A | Trhn-Gal4; R50H05-LexA>LexAOp-Gal80; UAS-Chr ((Trhn-R50H05)>Chr). | FigureS4A_Trhn-NOT-R50H05-Chr_Brain.czi, FigureS4A_Trhn-NOT-R50H05-Chr_VNC.czi |
| Figure S4B | Trhn-LexA; R50H05-Gal4>LexAOp-Gal80; UAS-Chr ((R50H05 - Trhn)>Chr). | FigureS4B_R50H05-NOT-Trhn-Chr_Brain.lsm, FigureS4B_R50H05-NOT-Trhn-Chr.lif |
| Figure S4C | *Trhn-Gal4>tsh-Gal80; UAS-Chr ((Trhn NOT tsh)>Chr)* | FigureS4C_Trhn-NOT-tsh-Chr.lif |
| Figure S4D | Trhn-Gal4>Tub-frt-Gal80-frt; tsh-LexA, 8X-LexAop2-FLPL/CyO-RFP-tb; UAS-Chr ((Trhn AND *tsh*)>Chr) | FigureS4D_Trhn-AND-tsh-Chr.lif |
| Figure S4E | Trhn-p65.AD; VT042751-GAL4.DBD>UAS-ACR1 (SXVNC1>ACR1) | FigureS4E_VNC1-ACR1_BRAIN_VNC.lif |
| Figure S4F | Trhn-p65.AD; R75E01-GAL4.DBD>UAS-ACR1 (SXVNC2>ACR1) | VNC2-ACR1_BRAIN_VNC.lif |

### Figure S5

#### Figure S5A Trhn-NOT-R50H05>Chr Volume

| control genotype | test genotype | control condition | test condition | control_N | test_N | effect_size | difference | bca_low | bca_high | pvalue_permutation |
|---|---|---|---|---|---|---|---|---|---|---|
| Trhn-Gal4; R50H05-LexA/w \| LexAOp-Gal80; UAS-Chr/w | Trhn>Chr/R50H050>G80 | Ctrl | Test | 80 | 40 | mean difference | -117.43 | -152.53 | -78.92 | 0.0 |

#### Figure S5B R50H05-NOT-Trhn>Chr Volume

| control genotype | test genotype | control condition | test condition | control_N | test_N | effect_size | difference | bca_low | bca_high | pvalue_permutation |
|---|---|---|---|---|---|---|---|---|---|---|
| Trhn-LexA; R50H05-Gal4/w \| LexAOp-Gal80; UAS-Chr/w | Trhn>G80/R50H050>Chr | Ctrl | Test | 60 | 30 | mean difference | 24.03 | -25.42 | 66.68 | 0.42 |

#### Figure S5C Trhn-NOT-tsh>Chr Volume

| control genotype | test genotype | control condition | test condition | control_N | test_N | effect_size | difference | bca_low | bca_high | pvalue_permutation |
|---|---|---|---|---|---|---|---|---|---|---|
| Trhn-Gal4/w \| tsh-Gal80, UAS-Chr/w | Trhn>Chr/tsh>G80 | Ctrl Red Light Off | Test Red Light Off | 35 | 25 | mean difference | 9.7 | -130.8 | 161.26 | 0.91 |
| Trhn-Gal4/w \| tsh-Gal80, UAS-Chr/w | Trhn>Chr/tsh>G80 | Ctrl Red Light On | Test Red Light On | 113 | 67 | mean difference | -26.63 | -89.41 | 42.21 | 0.4 |
| | | Test Red Light Off - Ctrl Red Light Off | Test Red Light On - Ctrl Red Light On | 5000 | 5000 | Delta Delta | -36.33 | -196.49 | 126.91 | 0.66 |

#### Figure S5D Trhn-AND-tsh>Chr Volume

| control genotype | test genotype | control condition | test condition | control_N | test_N | effect_size | difference | bca_low | bca_high | pvalue_permutation |
|---|---|---|---|---|---|---|---|---|---|---|
| Trhn-Gal4/w \| UAS-VNC-Chr/w | Trhn-VNC>Chr | Ctrl Red Light Off | Test Red Light Off | 32 | 28 | mean difference | 111.45 | 23.93 | 210.93 | 0.02 |
| Trhn-Gal4/w \| UAS-VNC-Chr/w | Trhn-VNC>Chr | Ctrl Red Light On | Test Red Light On | 139 | 88 | mean difference | -116.52 | -155.42 | -73.22 | 0.0 |
| | | Test Red Light Off - Ctrl Red Light Off | Test Red Light On - Ctrl Red Light On | 5000 | 5000 | Delta Delta | -227.97 | -332.02 | -128.32 | 0.0 |

#### Figure S5E Trhn-AND-tsh>ACR1 Volume

| control genotype | test genotype | control condition | test condition | control_N | test_N | effect_size | difference | bca_low | bca_high | pvalue_permutation |
|---|---|---|---|---|---|---|---|---|---|---|
| Trhn-Gal4/w \| UAS-VNC-ACR1/w | Trhn-VNC>ACR1 | Ctrl Green Light Off | Test Green Light Off | 51 | 38 | mean difference | 8.57 | -65.48 | 80.78 | 0.83 |
| Trhn-Gal4/w \| UAS-VNC-ACR1/w | Trhn-VNC>ACR1 | Ctrl Green Light On | Test Green Light On | 52 | 38 | mean difference | 42.09 | 4.7 | 72.11 | 0.02 |
| | | Test Green Light Off - Ctrl Green Light Off | Test Green Light On - Ctrl Green Light On | 5000 | 5000 | Delta Delta | 33.52 | -47.14 | 114.89 | 0.43 |

#### Figure S5F Trhn-AND-tsh>ACR1 Meal Size

| control genotype | test genotype | control condition | test condition | control_N | test_N | effect_size | difference | bca_low | bca_high | pvalue_permutation |
|---|---|---|---|---|---|---|---|---|---|---|
| Trhn-Gal4/w \| UAS-VNC-ACR1/w | Trhn-VNC>ACR1 | Ctrl Green Light Off | Test Green Light Off | 51 | 38 | mean difference | -6.46 | -15.08 | -1.7 | 0.06 |
| Trhn-Gal4/w \| UAS-VNC-ACR1/w | Trhn-VNC>ACR1 | Ctrl Green Light On | Test Green Light On | 52 | 38 | mean difference | 16.25 | 8.81 | 27.88 | 0.0 |
| | | Test Green Light Off - Ctrl Green Light Off | Test Green Light On - Ctrl Green Light On | 5000 | 5000 | Delta Delta | 22.71 | 13.12 | 34.88 | 0.0 |

#### Figure S5F SXVNC1>Chr Volume

| control genotype | test genotype | control condition | test condition | control_N | test_N | effect_size | difference | bca_low | bca_high | pvalue_permutation |
|---|---|---|---|---|---|---|---|---|---|---|
| Trhn-AD;Chr/w \| VT042751-DBD/w | SXVNC1>Chr | Ctrl Red Light Off | Test Red Light Off | 16 | 14 | mean difference | 91.16277679 | 0.0456875 | 189.4878125 | 0.0806 |
| Trhn-AD;Chr/w \| VT042751-DBD/w | SXVNC1>Chr | Ctrl Red Light On | Test Red Light On | 68 | 52 | mean difference | -159.3638077 | -229.83119 | -81.94863235 | 0 |
| | | Test Red Light Off - Ctrl Red Light Off | Test Red Light On - Ctrl Red Light On | 5000 | 5000 | Delta Delta | -250.5265845 | -363.7000814 | -131.1731325 | 0 |

#### Figure S5G SXVNC1>ACR1 Volume

| control genotype | test genotype | control condition | test condition | control_N | test_N | effect_size | difference | bca_low | bca_high | pvalue_permutation |
|---|---|---|---|---|---|---|---|---|---|---|
| ACR1/w \| SXVNC1/w | SXVNC1>ACR1 | Ctrl Green Light Off | Test Green Light Off | 16 | 14 | mean difference | -167.8492946 | -245.93625 | -114.1067321 | 0 |
| ACR1/w \| SXVNC1/w | SXVNC1>ACR1 | Ctrl Green Light On | Test Green Light On | 32 | 28 | mean difference | -117.2687187 | -157.0147188 | -77.90541071 | 0 |
| | | Test Green Light Off - Ctrl Green Light Off | Test Green Light On - Ctrl Green Light On | 5000 | 5000 | Delta Delta | 50.58057589 | -22.40814732 | 125.5160714 | 0.3618 |

#### Figure S5H SXVNC2>Chr Volume

| control genotype | test genotype | control condition | test condition | control_N | test_N | effect_size | difference | bca_low | bca_high | pvalue_permutation |
|---|---|---|---|---|---|---|---|---|---|---|
| Chr/w \| SXVNC2/w | SXVNC2>Chr | Ctrl Red Light Off | Test Red Light Off | 48 | 42 | mean difference | 51.9985506 | -7.050654762 | 115.7266875 | 0.0946 |
| Chr/w \| SXVNC2/w | SXVNC2>Chr | Ctrl Red Light On | Test Red Light On | 82 | 68 | mean difference | -129.3857306 | -170.2541808 | -86.16744727 | 0 |
| | | Test Red Light Off - Ctrl Red Light Off | Test Red Light On - Ctrl Red Light On | 5000 | 5000 | Delta Delta | -181.3842812 | -256.7174315 | -107.6165114 | 0 |

#### Figure S5I SXVNC2>ACR1 Volume

| control genotype | test genotype | control condition | test condition | control_N | test_N | effect_size | difference | bca_low | bca_high | pvalue_permutation |
|---|---|---|---|---|---|---|---|---|---|---|
| ACR1/w \| SXVNC2/w | SXVNC2>ACR1 | Ctrl Green Light Off | Test Green Light Off | 16 | 14 | mean difference | -34.10799107 | -104.6213571 | 33.55538393 | 0.3612 |
| ACR1/w \| SXVNC2/w | SXVNC2>ACR1 | Ctrl Green Light On | Test Green Light On | 16 | 14 | mean difference | 1.771375 | -18.937625 | 41.72279464 | 0.9088 |
| | | Test Green Light Off - Ctrl Green Light Off | Test Green Light On - Ctrl Green Light On | 5000 | 5000 | Delta Delta | 35.87936607 | -36.35684821 | 110.5884375 | 0.3296 |

**Figure S6A: Separate Control Plots for Figure S6**

**Figure S6A Trhn-NOT-R50H05>Chr Volume**

| control genotype | test genotype | control condition | test condition | control_N | test_N | effect_size | difference | bca_low | bca_high | pvalue_permutation |
|---|---|---|---|---|---|---|---|---|---|---|
| LexAOp;Chr/w | Trhn-G;R50/w | LexAOp-Gal80; UAS-Chr/w RLTG | Trhn-G; R50H05-L/w RLTG | 40 | 40 | mean difference | -46.1322 | -108.031375 | 12.91005 | 0.1568 |
| LexAOp;Chr/w | Trhn-G;R50>LexAOp;Chr | LexAOp-Gal80; UAS-Chr/w RLTG | Trhn-G>Chr/R50H050-L>G80 RLTG | 40 | 40 | mean difference | -140.500425 | -197.507775 | -87.582475 | 0 |

**Figure S6B R50H05-NOT-Trhn>Chr Volume**

| control genotype | test genotype | control condition | test condition | control_N | test_N | effect_size | difference | bca_low | bca_high | pvalue_permutation |
|---|---|---|---|---|---|---|---|---|---|---|
| LexAOp;Chr/w | Trhn-L;R50/w | LexAOp-Gal80; UAS-Chr/w TLRG | Trhn-L; R50H05-G/w TLRG | 30 | 30 | mean difference | -61.53083333 | -163.4665667 | -15.48336667 | 0.0418 |
| LexAOp;Chr/w | Trhn-L;R50>LexAOp;Chr | LexAOp-Gal80; UAS-Chr/w TLRG | Trhn-L>G80/R50H050-G>Chr TLRG | 30 | 30 | mean difference | -6.7315 | -118.1302 | 42.07366667 | 0.8896 |

**Figure S6C Trhn-NOT-tsh>Chr Volume**

| control genotype | test genotype | control condition | test condition | control_N | test_N | effect_size | difference | bca_low | bca_high | pvalue_permutation |
|---|---|---|---|---|---|---|---|---|---|---|
| Chr/w | Trhn/w | Brain-Chr/w Red Light Off | Trhn/w Red Light Off | 17 | 18 | mean difference | 19.18243137 | -231.2331863 | 144.841451 | 0.8474 |
| Chr/w | Trhn>Brain-Chr | Brain-Chr/w Red Light Off | Trhn>Brain-Chr Red Light Off | 17 | 25 | mean difference | 19.56064471 | -246.5321976 | 174.6868141 | 0.8578 |
| Chr/w | Trhn/w | Brain-Chr/w Red Light On | Trhn/w Red Light On | 55 | 58 | mean difference | 3.837683072 | -71.2792047 | 79.97085737 | 0.9214 |
| Chr/w | Trhn>Brain-Chr | Brain-Chr/w Red Light On | Trhn>Brain-Chr Red Light On | 55 | 67 | mean difference | -24.66242578 | -108.8692336 | 60.80622198 | 0.5726 |

**Figure S6D Trhn-AND-tsh>Chr Volume**

| control genotype | test genotype | control condition | test condition | control_N | test_N | effect_size | difference | bca_low | bca_high | pvalue_permutation |
|---|---|---|---|---|---|---|---|---|---|---|
| Chr/w | Trhn/w | VNC-Chr/w Red Light Off | Trhn/w Red Light Off | 16 | 16 | mean difference | -62.3679375 | -192.2163125 | 38.6489375 | 0.313 |
| Chr/w | Trhn>VNC-Chr | VNC-Chr/w Red Light Off | Trhn>VNC-Chr Red Light Off | 16 | 28 | mean difference | 80.27048214 | -62.65097321 | 197.5616161 | 0.2252 |
| Chr/w | Trhn/w | VNC-Chr/w Red Light On | Trhn/w Red Light On | 62 | 77 | mean difference | 111.3626169 | 48.26512505 | 168.2346297 | 0.001 |
| Chr/w | Trhn>VNC-Chr | VNC-Chr/w Red Light On | Trhn>VNC-Chr Red Light On | 62 | 88 | mean difference | -54.82985227 | -107.932952 | -3.932808651 | 0.0324 |

**Figure S6E Trhn-AND-tsh> ACR1 Volume**

| control genotype | test genotype | control condition | test condition | control_N | test_N | effect_size | difference | bca_low | bca_high | pvalue_permutation |
|---|---|---|---|---|---|---|---|---|---|---|
| VNC-ACR1/w | Trhn/w | VNC-ACR1/w Green Light Off | Trhn/w Green Light Off | 21 | 30 | mean difference | -156.326119 | -282.1338095 | -66.53815238 | 0.0014 |
| VNC-ACR1/w | Trhn>VNC-ACR1 | VNC-ACR1/w Green Light Off | Trhn>VNC-ACR1 Green Light Off | 21 | 38 | mean difference | -83.38690852 | -217.1585677 | 14.29651754 | 0.132 |
| VNC-ACR1/w | Trhn/w | VNC-ACR1/w Green Light On | Trhn/w Green Light On | 26 | 26 | mean difference | 48.56676923 | -1.516846154 | 97.96857692 | 0.065 |
| VNC-ACR1/w | Trhn>VNC-ACR1 | VNC-ACR1/w Green Light On | Trhn>VNC-ACR1 Green Light On | 26 | 38 | mean difference | 66.37637854 | 19.36527328 | 96.9039251 | 0.0004 |

**Figure S6F SXVNC1>Chr Volume**

| control genotype | test genotype | control condition | test condition | control_N | test_N | effect_size | difference | bca_low | bca_high | pvalue_permutation |
|---|---|---|---|---|---|---|---|---|---|---|
| Trhn-AD;Chr/w | VT042751-DBD/w | ACR1/w Green Light Off | SXVNC1/w Green Light Off | 7 | 9 | mean difference | -33.35071429 | -191.619254 | 81.68928571 | 0.633 |
| Trhn-AD;Chr/w | SXVNC1>Chr | ACR1/w Green Light Off | SXVNC1>ACR1 Green Light Off | 7 | 14 | mean difference | -186.6090714 | -334.7140714 | -79.58571429 | 0.0002 |
| Trhn-AD;Chr/w | VT042751-DBD/w | ACR1/w Green Light On | SXVNC1/w Green Light On | 14 | 18 | mean difference | 10.65956349 | -61.03434921 | 87.38357143 | 0.7904 |
| Trhn-AD;Chr/w | SXVNC1>Chr | ACR1/w Green Light On | SXVNC1>ACR1 Green Light On | 14 | 28 | mean difference | -111.2727143 | -164.1853571 | -54.33307143 | 0 |

**Figure S6G SXVNC1>ACR1 Volume**

| control genotype | test genotype | control condition | test condition | control_N | test_N | effect_size | difference | bca_low | bca_high | pvalue_permutation |
|---|---|---|---|---|---|---|---|---|---|---|
| ACR1/w | SXVNC1/w | ACR1/w Green Light Off | SXVNC1/w Green Light Off | 7 | 9 | mean difference | -33.35071429 | -191.619254 | 81.68928571 | 0.633 |
| ACR1/w | SXVNC1>ACR1 | ACR1/w Green Light Off | SXVNC1>ACR1 Green Light Off | 7 | 14 | mean difference | -186.6090714 | -334.7140714 | -79.58571429 | 0.0002 |
| ACR1/w | SXVNC1/w | ACR1/w Green Light On | SXVNC1/w Green Light On | 14 | 18 | mean difference | 10.65956349 | -61.03434921 | 87.38357143 | 0.7904 |
| ACR1/w | SXVNC1>ACR1 | ACR1/w Green Light On | SXVNC1>ACR1 Green Light On | 14 | 28 | mean difference | -111.2727143 | -164.1853571 | -54.33307143 | 0 |

**Figure S6H SXVNC2>Chr Volume**

| control genotype | test genotype | control condition | test condition | control_N | test_N | effect_size | difference | bca_low | bca_high | pvalue_permutation |
|---|---|---|---|---|---|---|---|---|---|---|
| Chr/w | SXVNC2/w | Chr/w Red Light Off | SXVNC2/w Red Light Off | 24 | 24 | mean difference | 85.00745833 | 23.827875 | 157.7880833 | 0.0172 |
| Chr/w | SXVNC2>Chr | Chr/w Red Light Off | SXVNC2>Chr Red Light Off | 24 | 42 | mean difference | 94.50227976 | 23.93966667 | 171.9557857 | 0.0216 |
| Chr/w | SXVNC2/w | Chr/w Red Light On | SXVNC2/w Red Light On | 41 | 41 | mean difference | 54.454 | -14.0212439 | 121.7067073 | 0.1242 |
| Chr/w | SXVNC2>Chr | Chr/w Red Light On | SXVNC2>Chr Red Light On | 41 | 68 | mean difference | -102.1587306 | -158.1093257 | -50.31224103 | 0.0004 |

**Figure S6I SXVNC2>ACR1 Volume**

| control genotype | test genotype | control condition | test condition | control_N | test_N | effect_size | difference | bca_low | bca_high | pvalue_permutation |
|---|---|---|---|---|---|---|---|---|---|---|
| ACR1/w | SXVNC2/w | ACR1/w Green Light Off | SXVNC2/w Green Light Off | 8 | 8 | mean difference | 3.482625 | -118.00825 | 94.53725 | 0.9446 |
| ACR1/w | SXVNC2>ACR1 | ACR1/w Green Light Off | SXVNC2>ACR1 Green Light Off | 8 | 14 | mean difference | -32.36667857 | -158.3811786 | 55.37291071 | 0.5338 |
| ACR1/w | SXVNC2/w | ACR1/w Green Light On | SXVNC2/w Green Light On | 8 | 8 | mean difference | -7.776 | -36.80925 | 22.593875 | 0.6324 |
| ACR1/w | SXVNC2>ACR1 | ACR1/w Green Light On | SXVNC2>ACR1 Green Light On | 8 | 14 | mean difference | -2.116625 | -32.37041071 | 31.10498214 | 0.929 |

### Figure 5

#### Figure 5A

| | Volume | Duration | Meal Duration | Feed Speed | Meal Size | Food Port Occupancy | Height | Count | Duringfeed Speed Ratio | Falls | Prefeed Speed | Ctrl Port Occupancy | Speed | Postfeed Speed | Perifeed Speed Ratio | Latency | Duringfeed Speed |
|---|---|---|---|---|---|---|---|---|---|---|---|---|---|---|---|---|---|
| mean 24 | 0.74 | 0.67 | -0.37 | -0.84 | -0.86 | 0.84 | 0.78 | -0.09 | 0.5 | -0.34 | 0.16 | -0.46 | -0.45 | 0.18 | 0.11 | -0.5 | 0.65 |
| bca_low 24 | 0.44 | 0.4 | -0.66 | -1.08 | -1.16 | 0.56 | 0.53 | -0.31 | 0.24 | -0.58 | -0.12 | -0.86 | -0.62 | -0.07 | -0.14 | -0.66 | 0.44 |
| bca_high 24 | 0.93 | 0.94 | -0.04 | -0.54 | -0.57 | 1.04 | 1.01 | 0.14 | 0.75 | -0.1 | 0.41 | -0.1 | -0.27 | 0.41 | 0.32 | -0.23 | 0.86 |
| mean 48 | 1.25 | 1.26 | -0.37 | -1.0 | -0.7 | 1.28 | 1.91 | 0.29 | 1.3 | -0.85 | 0.13 | -0.73 | -0.52 | 0.32 | 0.51 | -0.49 | 0.05 |
| bca_low 48 | 0.93 | 0.93 | -0.69 | -1.24 | -1.03 | 0.96 | 1.56 | 0.02 | 1.0 | -1.13 | -0.19 | -1.05 | -0.68 | 0.04 | 0.24 | -0.63 | -0.21 |
| bca_high 48 | 1.54 | 1.59 | -0.03 | -0.74 | -0.37 | 1.56 | 2.25 | 0.57 | 1.62 | -0.57 | 0.42 | -0.37 | -0.34 | 0.59 | 0.76 | -0.31 | 0.33 |

#### Figure 5B Trhn>Chr

| | Volume | Duration | Meal Duration | Feed Speed | Meal Size | Food Port Occupancy | Height | Count | Duringfeed Speed Ratio | Falls | Prefeed Speed | Ctrl Port Occupancy | Speed | Postfeed Speed | Perifeed Speed Ratio | Latency | Duringfeed Speed |
|---|---|---|---|---|---|---|---|---|---|---|---|---|---|---|---|---|---|
| mean 24 | -1.44 | -0.33 | -1.33 | -1.74 | -1.44 | -0.07 | -1.12 | 0.38 | 0.14 | -0.17 | 0.02 | 0.01 | 0.6 | 0.27 | 0.83 | 1.15 | 0.49 |
| bca_low 24 | -1.87 | -0.78 | -1.93 | -2.25 | -2.05 | -0.42 | -1.55 | -0.03 | -0.31 | -0.69 | -0.47 | -0.35 | 0.16 | -0.19 | 0.38 | 0.7 | -0.07 |
| bca_high 24 | -1.03 | 0.11 | -0.77 | -1.24 | -0.89 | 0.26 | -0.7 | 0.79 | 0.65 | 0.32 | 0.48 | 0.29 | 1.04 | 0.71 | 1.36 | 1.62 | 0.96 |

#### Figure 5B Trhn>ACR1

| | Volume | Duration | Meal Duration | Feed Speed | Meal Size | Food Port Occupancy | Height | Count | Duringfeed Speed Ratio | Falls | Prefeed Speed | Ctrl Port Occupancy | Speed | Postfeed Speed | Perifeed Speed Ratio | Latency | Duringfeed Speed |
|---|---|---|---|---|---|---|---|---|---|---|---|---|---|---|---|---|---|
| mean 24 | 1.3 | 0.75 | 1.56 | 1.21 | 1.76 | 0.36 | 0.65 | -0.22 | -0.61 | -0.03 | -0.53 | 0.68 | -0.35 | -0.61 | -0.46 | -0.69 | -0.9 |
| bca_low 24 | 0.68 | 0.17 | 0.99 | 0.67 | 1.12 | -0.2 | 0.15 | -0.75 | -1.33 | -0.57 | -1.12 | 0.1 | -0.95 | -1.23 | -1.15 | -1.2 | -1.5 |
| bca_high 24 | 2.0 | 1.37 | 2.22 | 1.77 | 2.49 | 0.91 | 1.15 | 0.34 | 0.06 | 0.58 | 0.08 | 1.59 | 0.28 | 0.04 | 0.19 | -0.22 | -0.32 |

#### Figure 5C R50H05>Chr

| | Volume | Duration | Meal Duration | Feed Speed | Meal Size | Food Port Occupancy | Height | Count | Duringfeed Speed Ratio | Falls | Prefeed Speed | Ctrl Port Occupancy | Speed | Postfeed Speed | Perifeed Speed Ratio | Latency | Duringfeed Speed |
|---|---|---|---|---|---|---|---|---|---|---|---|---|---|---|---|---|---|
| mean 24 | 2.61 | 1.67 | 0.39 | 1.13 | 0.53 | -0.07 | -0.67 | 1.57 | -0.96 | -0.03 | 0.98 | -0.36 | 1.83 | 1.14 | -0.14 | -1.3 | 0.55 |
| bca_low 24 | 2.16 | 1.21 | -0.12 | 0.67 | -0.02 | -0.56 | -1.07 | 1.14 | -1.81 | -0.41 | 0.5 | -0.71 | 1.44 | 0.67 | -0.82 | -1.68 | 0.09 |
| bca_high 24 | 3.11 | 2.1 | 0.87 | 1.57 | 0.98 | 0.31 | -0.27 | 2.01 | -0.34 | 0.38 | 1.45 | -0.12 | 2.25 | 1.61 | 0.44 | -0.92 | 0.99 |

#### Figure 5C R50H05>ACR1

| | Volume | Duration | Meal Duration | Feed Speed | Meal Size | Food Port Occupancy | Height | Count | Duringfeed Speed Ratio | Falls | Prefeed Speed | Ctrl Port Occupancy | Speed | Postfeed Speed | Perifeed Speed Ratio | Latency | Duringfeed Speed |
|---|---|---|---|---|---|---|---|---|---|---|---|---|---|---|---|---|---|
| mean 24 | -0.13 | -0.09 | 0.01 | 0.07 | 0.08 | -0.12 | 0.29 | -0.02 | -0.1 | -0.16 | -0.03 | 0.01 | 0.06 | 0.09 | -0.38 | -0.01 | -0.1 |
| bca_low 24 | -0.7 | -0.65 | -0.77 | -0.57 | -0.66 | -0.57 | -0.36 | -0.67 | -0.66 | -0.9 | -0.72 | -0.41 | -0.57 | -0.6 | -1.13 | -0.65 | -0.79 |
| bca_high 24 | 0.45 | 0.49 | 0.74 | 0.7 | 0.74 | 0.39 | 0.97 | 0.62 | 0.46 | 0.51 | 0.67 | 0.54 | 0.76 | 0.8 | 0.29 | 0.6 | 0.7 |

#### Figure 5D

| | Volume | Feed Speed | Meal Size | Meal Duration | Duration | Count | Height | Food Port Occupancy | Ctrl Port Occupancy | Latency | Speed | Prefeed Speed | Duringfeed Speed | Postfeed Speed | Duringfeed Speed Ratio | Perifeed Speed Ratio | Falls |
|---|---|---|---|---|---|---|---|---|---|---|---|---|---|---|---|---|---|
| Trhn>Chr | -1.44 | -1.74 | -1.44 | -1.33 | -0.33 | 0.38 | -1.12 | -0.07 | 0.01 | 1.15 | 0.6 | 0.02 | 0.49 | 0.27 | 0.14 | 0.83 | -0.17 |
| R50H05>Chr | 2.61 | 1.13 | 0.53 | 0.39 | 1.67 | 1.57 | -0.67 | -0.07 | -0.36 | -1.3 | 1.83 | 0.98 | 0.55 | 1.14 | -0.96 | -0.14 | -0.03 |
| Trhn>ACR1 | 1.3 | 1.21 | 1.76 | 1.56 | 0.75 | -0.22 | 0.65 | 0.36 | 0.68 | -0.69 | -0.35 | -0.53 | -0.9 | -0.61 | -0.61 | -0.46 | -0.03 |
| R50H05>ACR1 | -0.13 | 0.07 | 0.08 | 0.01 | -0.09 | -0.02 | 0.29 | -0.12 | 0.01 | -0.01 | 0.06 | -0.03 | -0.1 | 0.09 | -0.1 | -0.38 | -0.16 |
| Trhn-AND-tsh>Chr | -1.35 | -0.64 | -0.61 | -0.33 | -0.64 | -1.12 | -0.16 | -0.17 | -0.79 | -0.15 | -1.05 | 0.01 | 0.03 | -0.14 | -0.05 | -0.24 | 0.25 |
| Trhn-AND-tsh>ACR1 | 0.27 | 1.08 | 1.28 | 1.27 | -0.01 | -0.4 | -0.07 | -0.57 | -0.49 | -0.87 | -0.93 | 0.0 | -0.51 | 0.09 | -0.13 | 0.44 | -0.41 |
| SXVNC1>Chr | -1.29 | -1.23 | -0.54 | -0.55 | -0.8 | -0.67 | -0.62 | -0.06 | -0.14 | 0.88 | -0.24 | 0.71 | 0.73 | -0.1 | 0.01 | -0.56 | -0.47 |
| SXVNC2>Chr | -1.29 | -0.62 | -0.58 | -0.64 | -0.92 | -0.45 | 0.06 | -0.5 | -0.22 | 0.64 | -0.39 | -0.17 | 0.37 | -0.32 | 0.36 | -0.1 | 0.25 |
| SXVNC2>ACR1 | 0.5 | 0.25 | 0.43 | 0.36 | 0.47 | 0.53 | 0.33 | 0.59 | 0.67 | -0.27 | 0.09 | -0.52 | -1.46 | -0.09 | -1.01 | -0.38 | -0.11 |
| Trhn-NOT-R50H05>Chr | -0.94 | -0.97 | -0.67 | -0.63 | -0.46 | -0.08 | -1.34 | 0.24 | -0.17 | 0.79 | -0.29 | -0.46 | 0.66 | -0.12 | 0.38 | 0.42 | -0.02 |
| R50H05-NOT-Trhn>Chr | 0.2 | 0.25 | 0.03 | 0.5 | 0.32 | -0.16 | 0.25 | -0.14 | -0.61 | 0.0 | -0.35 | -1.01 | -0.78 | -0.94 | -0.06 | -0.07 | 0.11 |
| Trhn-NOT-tsh>Chr | -0.16 | -0.95 | -0.76 | -0.54 | 0.94 | 1.13 | -0.29 | 1.31 | 0.18 | 0.35 | 0.44 | -1.3 | -0.52 | -0.79 | 0.41 | 0.28 | -0.03 |
| NPF>Chr | 1.29 | 0.35 | 0.51 | 1.47 | 1.96 | 1.56 | -0.43 | 1.06 | -0.09 | -0.81 | 2.06 | 1.01 | -0.39 | 1.21 | -1.06 | -0.02 | 0.31 |
| NPF>ACR1 | -0.06 | -0.47 | -0.38 | -0.16 | 0.39 | 0.21 | 0.42 | -0.12 | -0.74 | -0.24 | 0.52 | 0.65 | 0.08 | 0.2 | 0.13 | -0.4 | -0.58 |
| Akh>Chr | -0.58 | -0.64 | -0.12 | -0.47 | -0.36 | -0.0 | -0.5 | 0.05 | 0.3 | 0.43 | 0.67 | 0.47 | 1.11 | 0.74 | 0.49 | 0.82 | 0.47 |
| Akh>ACR1 | 0.29 | 0.02 | 0.11 | 0.28 | 0.37 | 0.28 | -0.06 | 0.48 | -0.04 | 0.02 | 0.22 | -0.79 | -0.86 | -0.83 | 0.51 | -0.26 | 0.04 |
| Ilp2>Chr | 1.25 | 0.05 | -0.17 | -0.31 | 1.13 | 1.05 | -0.02 | 0.42 | -0.12 | -0.52 | 0.51 | 0.08 | -0.01 | -0.21 | 0.24 | -0.25 | -0.44 |
| Ilp2>ACR1 | 0.59 | 0.08 | 0.52 | 0.34 | 0.44 | 0.0 | 0.31 | 0.6 | 0.11 | 0.19 | -0.14 | 0.11 | -0.34 | -0.13 | 0.52 | -0.2 | -0.0 |
| 24 h starved | 0.78 | 0.67 | 0.74 | 0.84 | 0.5 | -0.09 | 0.18 | 0.11 | -0.5 | -0.34 | -0.45 | -0.37 | -0.84 | -0.86 | 0.16 | -0.46 | 0.65 |
| 24 h reversed | -0.78 | -0.67 | -0.74 | -0.84 | -0.5 | 0.09 | -0.18 | -0.11 | 0.5 | 0.34 | 0.45 | 0.37 | 0.84 | 0.86 | -0.16 | 0.46 | -0.65 |
| 48 h starved | 1.91 | 1.26 | 1.25 | 1.28 | 1.3 | 0.29 | 0.32 | 0.51 | -0.49 | -0.85 | -0.52 | -0.37 | -1.0 | -0.7 | 0.13 | -0.73 | 0.05 |
| 48 h reversed | -1.91 | -1.26 | -1.25 | -1.28 | -1.3 | -0.29 | -0.32 | -0.51 | 0.49 | 0.85 | 0.52 | 0.37 | 1.0 | 0.7 | -0.13 | 0.73 | -0.05 |

**Figure S7**

**Figure S7A**

| Metric | Volume | Duration | Feed Speed | Count | Meal Size | Meal Duration | Food Port Occupancy | Height | Speed | Ctrl Port Occupancy | Falls | Postfeed Speed | Prefeed Speed | Perifeed Speed Ratio | Duringfeed Speed Ratio | Duringfeed Speed | Latency |
|---|---|---|---|---|---|---|---|---|---|---|---|---|---|---|---|---|---|
| Volume | 1.0 | 0.87 | 0.61 | 0.59 | 0.44 | 0.42 | 0.33 | 0.22 | 0.18 | 0.12 | 0.03 | -0.01 | -0.03 | -0.08 | -0.09 | -0.13 | -0.64 |
| Duration | 0.87 | 1.0 | 0.44 | 0.83 | 0.24 | 0.3 | 0.52 | 0.27 | 0.32 | 0.28 | 0.06 | 0.04 | -0.07 | -0.02 | -0.08 | -0.11 | -0.66 |
| Feed Speed | 0.61 | 0.44 | 1.0 | 0.21 | 0.68 | 0.67 | 0.1 | 0.19 | 0.09 | -0.0 | -0.0 | -0.24 | 0.06 | -0.26 | -0.12 | -0.4 | -0.72 |
| Count | 0.59 | 0.83 | 0.21 | 1.0 | -0.09 | -0.03 | 0.45 | 0.2 | 0.6 | 0.53 | -0.0 | 0.37 | 0.11 | 0.2 | -0.11 | 0.23 | -0.58 |
| Meal Size | 0.44 | 0.24 | 0.68 | -0.09 | 1.0 | 0.96 | 0.02 | 0.09 | -0.11 | -0.13 | -0.02 | -0.32 | 0.01 | -0.32 | -0.04 | -0.35 | -0.31 |
| Meal Duration | 0.42 | 0.3 | 0.67 | -0.03 | 0.96 | 1.0 | 0.09 | 0.13 | -0.09 | -0.11 | 0.03 | -0.38 | -0.07 | -0.34 | -0.01 | -0.41 | -0.37 |
| Food Port Occupancy | 0.33 | 0.52 | 0.1 | 0.45 | 0.02 | 0.09 | 1.0 | 0.2 | 0.24 | 0.23 | -0.0 | 0.04 | -0.06 | 0.06 | 0.09 | -0.01 | -0.3 |
| Height | 0.22 | 0.27 | 0.19 | 0.2 | 0.09 | 0.13 | 0.2 | 1.0 | -0.01 | 0.05 | -0.05 | -0.25 | -0.27 | -0.07 | 0.31 | -0.11 | -0.28 |
| Speed | 0.18 | 0.32 | 0.09 | 0.6 | -0.11 | -0.09 | 0.24 | -0.01 | 1.0 | 0.77 | -0.11 | 0.68 | 0.47 | 0.24 | -0.23 | 0.55 | -0.41 |
| Ctrl Port Occupancy | 0.12 | 0.28 | -0.0 | 0.53 | -0.13 | -0.11 | 0.23 | 0.05 | 0.77 | 1.0 | -0.12 | 0.54 | 0.34 | 0.2 | -0.17 | 0.44 | -0.31 |
| Falls | 0.03 | 0.06 | -0.0 | -0.0 | -0.02 | 0.03 | -0.0 | -0.05 | -0.11 | -0.12 | 1.0 | -0.13 | -0.11 | -0.09 | -0.04 | -0.15 | 0.02 |
| Postfeed Speed | -0.01 | 0.04 | -0.24 | 0.37 | -0.32 | -0.38 | 0.04 | -0.25 | 0.68 | 0.54 | -0.13 | 1.0 | 0.63 | 0.51 | -0.31 | 0.59 | -0.23 |
| Prefeed Speed | -0.03 | -0.07 | 0.06 | 0.11 | 0.01 | -0.07 | -0.06 | -0.27 | 0.47 | 0.34 | -0.11 | 0.63 | 1.0 | -0.13 | -0.56 | 0.39 | -0.14 |
| Perifeed Speed Ratio | -0.08 | -0.02 | -0.26 | 0.2 | -0.32 | -0.34 | 0.06 | -0.07 | 0.24 | 0.2 | -0.09 | 0.51 | -0.13 | 1.0 | 0.34 | 0.24 | -0.1 |
| Duringfeed Speed Ratio | -0.09 | -0.08 | -0.12 | -0.11 | -0.04 | -0.01 | 0.09 | 0.31 | -0.23 | -0.17 | -0.04 | -0.31 | -0.56 | 0.34 | 1.0 | 0.05 | 0.14 |
| Duringfeed Speed | -0.13 | -0.11 | -0.4 | 0.23 | -0.35 | -0.41 | -0.01 | -0.11 | 0.55 | 0.44 | -0.15 | 0.59 | 0.39 | 0.24 | 0.05 | 1.0 | -0.12 |
| Latency | -0.64 | -0.66 | -0.72 | -0.58 | -0.31 | -0.37 | -0.3 | -0.28 | -0.41 | -0.31 | 0.02 | -0.23 | -0.14 | -0.1 | 0.14 | -0.12 | 1.0 |

**Figure S7H**

| Metric | Volume | Duration | Count | Speed | Food Port Occupancy | Feed Speed | Ctrl Port Occupancy | Meal Duration | Postfeed Speed | Meal Size | Prefeed Speed | Duringfeed Speed | Height | Perifeed Speed Ratio | Duringfeed Speed Ratio | Falls | Latency |
|---|---|---|---|---|---|---|---|---|---|---|---|---|---|---|---|---|---|
| Volume | 1.0 | 0.84 | 0.79 | 0.67 | 0.67 | 0.59 | 0.59 | 0.41 | 0.37 | 0.35 | 0.33 | 0.23 | 0.09 | 0.06 | -0.16 | -0.3 | -0.69 |
| Duration | 0.84 | 1.0 | 0.96 | 0.74 | 0.9 | 0.36 | 0.7 | 0.3 | 0.34 | 0.13 | 0.21 | 0.1 | 0.17 | 0.15 | -0.18 | -0.25 | -0.69 |
| Count | 0.79 | 0.96 | 1.0 | 0.75 | 0.88 | 0.28 | 0.71 | 0.16 | 0.43 | 0.01 | 0.24 | 0.24 | 0.2 | 0.23 | -0.14 | -0.24 | -0.66 |
| Speed | 0.67 | 0.74 | 0.75 | 1.0 | 0.81 | 0.29 | 0.8 | 0.18 | 0.69 | 0.05 | 0.61 | 0.43 | 0.09 | 0.13 | -0.25 | -0.26 | -0.61 |
| Food Port Occupancy | 0.67 | 0.9 | 0.88 | 0.81 | 1.0 | 0.24 | 0.79 | 0.21 | 0.44 | 0.04 | 0.28 | 0.16 | 0.18 | 0.16 | -0.2 | -0.22 | -0.61 |
| Feed Speed | 0.59 | 0.36 | 0.28 | 0.29 | 0.24 | 1.0 | 0.17 | 0.71 | -0.23 | 0.67 | -0.04 | -0.26 | 0.11 | -0.22 | -0.02 | -0.36 | -0.7 |
| Ctrl Port Occupancy | 0.59 | 0.7 | 0.71 | 0.8 | 0.79 | 0.17 | 1.0 | 0.11 | 0.55 | -0.0 | 0.43 | 0.37 | 0.12 | 0.15 | -0.18 | -0.21 | -0.52 |
| Meal Duration | 0.41 | 0.3 | 0.16 | 0.18 | 0.21 | 0.71 | 0.11 | 1.0 | -0.31 | 0.94 | 0.1 | -0.29 | 0.01 | -0.39 | -0.12 | -0.28 | -0.44 |
| Postfeed Speed | 0.37 | 0.34 | 0.43 | 0.69 | 0.44 | -0.23 | 0.55 | -0.31 | 1.0 | -0.26 | 0.74 | 0.58 | -0.05 | 0.41 | -0.3 | -0.09 | -0.14 |
| Meal Size | 0.35 | 0.13 | 0.01 | 0.05 | 0.04 | 0.67 | -0.0 | 0.94 | -0.26 | 1.0 | 0.16 | -0.19 | -0.05 | -0.38 | -0.11 | -0.23 | -0.27 |
| Prefeed Speed | 0.33 | 0.21 | 0.24 | 0.61 | 0.28 | -0.04 | 0.43 | 0.1 | 0.74 | 0.16 | 1.0 | 0.59 | -0.26 | -0.14 | -0.44 | -0.14 | -0.02 |
| Duringfeed Speed | 0.23 | 0.1 | 0.24 | 0.43 | 0.16 | -0.26 | 0.37 | -0.29 | 0.58 | -0.19 | 0.59 | 1.0 | -0.12 | 0.01 | 0.11 | -0.2 | -0.29 |
| Height | 0.09 | 0.17 | 0.2 | 0.09 | 0.18 | 0.11 | 0.12 | 0.01 | -0.05 | -0.05 | -0.26 | -0.12 | 1.0 | 0.27 | 0.13 | -0.04 | -0.26 |
| Perifeed Speed Ratio | 0.06 | 0.15 | 0.23 | 0.13 | 0.16 | -0.22 | 0.15 | -0.39 | 0.41 | -0.38 | -0.14 | 0.01 | 0.27 | 1.0 | 0.43 | -0.0 | -0.22 |
| Duringfeed Speed Ratio | -0.16 | -0.18 | -0.14 | -0.25 | -0.2 | -0.02 | -0.18 | -0.12 | -0.3 | -0.11 | -0.44 | 0.11 | 0.13 | 0.43 | 1.0 | -0.11 | -0.2 |
| Falls | -0.3 | -0.25 | -0.24 | -0.26 | -0.22 | -0.36 | -0.21 | -0.28 | -0.09 | -0.23 | -0.14 | -0.2 | -0.04 | -0.0 | -0.11 | 1.0 | 0.42 |
| Latency | -0.69 | -0.69 | -0.66 | -0.61 | -0.61 | -0.7 | -0.52 | -0.44 | -0.14 | -0.27 | -0.02 | -0.29 | -0.26 | -0.22 | -0.2 | 0.42 | 1.0 |

**Figure S7I**

| Metric | Volume | Duration | Count | Speed | Feed Speed | Ctrl Port Occupancy | Falls | Meal Size | Meal Duration | Food Port Occupancy | Postfeed Speed | Height | Prefeed Speed | Perifeed Speed Ratio | Duringfeed Speed | Duringfeed Speed Ratio | Latency |
|---|---|---|---|---|---|---|---|---|---|---|---|---|---|---|---|---|---|
| Volume | 1.0 | 0.91 | 0.79 | 0.59 | 0.54 | 0.26 | 0.24 | 0.24 | 0.24 | 0.22 | 0.19 | 0.13 | 0.06 | 0.04 | 0.04 | -0.17 | -0.67 |
| Duration | 0.91 | 1.0 | 0.91 | 0.57 | 0.42 | 0.31 | 0.23 | 0.11 | 0.18 | 0.41 | 0.15 | 0.2 | -0.03 | 0.03 | -0.0 | -0.14 | -0.68 |
| Count | 0.79 | 0.91 | 1.0 | 0.64 | 0.27 | 0.31 | 0.26 | -0.11 | -0.07 | 0.29 | 0.34 | 0.13 | 0.03 | 0.15 | 0.17 | -0.14 | -0.59 |
| Speed | 0.59 | 0.57 | 0.64 | 1.0 | 0.3 | 0.44 | 0.19 | -0.02 | -0.03 | 0.05 | 0.62 | -0.09 | 0.36 | 0.16 | 0.4 | -0.34 | -0.52 |
| Feed Speed | 0.54 | 0.42 | 0.27 | 0.3 | 1.0 | 0.11 | 0.18 | 0.66 | 0.65 | 0.01 | 0.02 | 0.15 | 0.36 | -0.14 | -0.4 | -0.23 | -0.7 |
| Ctrl Port Occupancy | 0.26 | 0.31 | 0.31 | 0.44 | 0.11 | 1.0 | 0.06 | -0.04 | -0.02 | 0.04 | 0.27 | 0.08 | 0.14 | 0.05 | 0.11 | -0.16 | -0.26 |
| Falls | 0.24 | 0.23 | 0.26 | 0.19 | 0.18 | 0.06 | 1.0 | 0.01 | 0.05 | 0.03 | 0.07 | -0.13 | -0.07 | 0.02 | 0.02 | -0.08 | -0.19 |
| Meal Size | 0.24 | 0.11 | -0.11 | -0.02 | 0.66 | -0.04 | 0.01 | 1.0 | 0.95 | -0.02 | -0.21 | 0.06 | 0.28 | -0.19 | -0.36 | -0.11 | -0.28 |
| Meal Duration | 0.24 | 0.18 | -0.07 | -0.03 | 0.65 | -0.02 | 0.05 | 0.95 | 1.0 | 0.05 | -0.3 | 0.09 | 0.13 | -0.19 | -0.41 | -0.06 | -0.33 |
| Food Port Occupancy | 0.22 | 0.41 | 0.29 | 0.05 | 0.01 | 0.04 | 0.03 | -0.02 | 0.05 | 1.0 | -0.06 | 0.18 | -0.13 | 0.05 | -0.08 | 0.14 | -0.17 |
| Postfeed Speed | 0.19 | 0.15 | 0.34 | 0.62 | 0.02 | 0.27 | 0.07 | -0.21 | -0.3 | -0.06 | 1.0 | -0.37 | 0.41 | 0.53 | 0.42 | -0.26 | -0.38 |
| Height | 0.13 | 0.2 | 0.13 | -0.09 | 0.15 | 0.08 | -0.13 | 0.06 | 0.09 | 0.18 | -0.37 | 1.0 | -0.14 | -0.26 | -0.06 | 0.26 | -0.18 |
| Prefeed Speed | 0.06 | -0.03 | 0.03 | 0.36 | 0.36 | 0.14 | -0.07 | 0.28 | 0.13 | -0.13 | 0.41 | -0.14 | 1.0 | -0.29 | 0.12 | -0.58 | -0.18 |
| Perifeed Speed Ratio | 0.04 | 0.03 | 0.15 | 0.16 | -0.14 | 0.05 | 0.02 | -0.19 | -0.19 | 0.05 | 0.53 | -0.26 | -0.29 | 1.0 | 0.24 | 0.42 | -0.08 |
| Duringfeed Speed | 0.04 | -0.0 | 0.17 | 0.4 | -0.4 | 0.11 | 0.02 | -0.36 | -0.41 | -0.08 | 0.42 | -0.06 | 0.12 | 0.24 | 1.0 | 0.14 | -0.03 |
| Duringfeed Speed Ratio | -0.17 | -0.14 | -0.14 | -0.34 | -0.23 | -0.16 | -0.08 | -0.11 | -0.06 | 0.14 | -0.26 | 0.26 | -0.58 | 0.42 | 0.14 | 1.0 | 0.26 |
| Latency | -0.67 | -0.68 | -0.59 | -0.52 | -0.7 | -0.26 | -0.19 | -0.28 | -0.33 | -0.17 | -0.38 | -0.18 | -0.18 | -0.08 | -0.03 | 0.26 | 1.0 |

**Figure S7J**

| Metric | Volume | Duration | Food Port Occupancy | Count | Speed | Feed Speed | Ctrl Port Occupancy | Height | Duringfeed Speed | Meal Size | Meal Duration | Postfeed Speed | Perifeed Speed Ratio | Prefeed Speed | Falls | Duringfeed Speed Ratio | Latency |
|---|---|---|---|---|---|---|---|---|---|---|---|---|---|---|---|---|---|
| Volume | 1.0 | 0.81 | 0.63 | 0.61 | 0.51 | 0.48 | 0.36 | 0.3 | 0.24 | 0.22 | 0.18 | 0.13 | 0.12 | -0.02 | -0.05 | -0.14 | -0.64 |
| Duration | 0.81 | 1.0 | 0.9 | 0.86 | 0.39 | 0.16 | 0.34 | 0.36 | 0.05 | -0.03 | 0.02 | 0.03 | 0.09 | -0.14 | 0.13 | -0.04 | -0.57 |
| Food Port Occupancy | 0.63 | 0.9 | 1.0 | 0.77 | 0.25 | 0.02 | 0.29 | 0.35 | 0.05 | -0.09 | -0.04 | -0.0 | 0.1 | -0.16 | 0.06 | 0.07 | -0.41 |
| Count | 0.61 | 0.86 | 0.77 | 1.0 | 0.55 | -0.02 | 0.49 | 0.24 | 0.21 | -0.33 | -0.29 | 0.24 | 0.25 | 0.0 | 0.07 | -0.07 | -0.43 |
| Speed | 0.51 | 0.39 | 0.25 | 0.55 | 1.0 | 0.14 | 0.82 | -0.07 | 0.56 | -0.1 | -0.13 | 0.74 | 0.37 | 0.52 | -0.13 | -0.36 | -0.39 |
| Feed Speed | 0.48 | 0.16 | 0.02 | -0.02 | 0.14 | 1.0 | -0.01 | 0.21 | -0.19 | 0.6 | 0.55 | -0.14 | -0.23 | 0.07 | -0.14 | -0.17 | -0.62 |
| Ctrl Port Occupancy | 0.36 | 0.34 | 0.29 | 0.49 | 0.82 | -0.01 | 1.0 | 0.01 | 0.46 | -0.16 | -0.17 | 0.62 | 0.34 | 0.4 | -0.1 | -0.28 | -0.29 |
| Height | 0.3 | 0.36 | 0.35 | 0.24 | -0.07 | 0.21 | 0.01 | 1.0 | 0.02 | 0.08 | 0.13 | -0.28 | 0.08 | -0.39 | 0.03 | 0.5 | -0.37 |
| Duringfeed Speed | 0.24 | 0.05 | 0.05 | 0.21 | 0.56 | -0.19 | 0.46 | 0.02 | 1.0 | -0.23 | -0.31 | 0.57 | 0.34 | 0.35 | -0.07 | 0.0 | -0.28 |
| Meal Size | 0.22 | -0.03 | -0.09 | -0.33 | -0.1 | 0.6 | -0.16 | 0.08 | -0.23 | 1.0 | 0.97 | -0.27 | -0.35 | -0.1 | -0.05 | -0.0 | -0.11 |
| Meal Duration | 0.18 | 0.02 | -0.04 | -0.29 | -0.13 | 0.55 | -0.17 | 0.13 | -0.31 | 0.97 | 1.0 | -0.33 | -0.4 | -0.15 | 0.03 | 0.02 | -0.14 |
| Postfeed Speed | 0.13 | 0.03 | -0.0 | 0.24 | 0.74 | -0.14 | 0.62 | -0.28 | 0.57 | -0.27 | -0.33 | 1.0 | 0.54 | 0.7 | -0.11 | -0.44 | -0.33 |
| Perifeed Speed Ratio | 0.12 | 0.09 | 0.1 | 0.25 | 0.37 | -0.23 | 0.34 | 0.08 | 0.34 | -0.35 | -0.4 | 0.54 | 1.0 | -0.03 | -0.18 | 0.25 | -0.16 |
| Prefeed Speed | -0.02 | -0.14 | -0.16 | 0.0 | 0.52 | 0.07 | 0.4 | -0.39 | 0.35 | -0.1 | -0.15 | 0.7 | -0.03 | 1.0 | -0.07 | -0.63 | -0.3 |
| Falls | -0.05 | 0.13 | 0.06 | 0.07 | -0.13 | -0.14 | -0.1 | 0.03 | -0.07 | -0.05 | 0.03 | -0.11 | -0.18 | -0.07 | 1.0 | -0.04 | -0.13 |
| Duringfeed Speed Ratio | -0.14 | -0.04 | 0.07 | -0.07 | -0.36 | -0.17 | -0.28 | 0.5 | 0.0 | -0.0 | 0.02 | -0.44 | 0.25 | -0.63 | -0.04 | 1.0 | 0.21 |
| Latency | -0.64 | -0.57 | -0.41 | -0.43 | -0.39 | -0.62 | -0.29 | -0.37 | -0.28 | -0.11 | -0.14 | -0.33 | -0.16 | -0.3 | -0.13 | 0.21 | 1.0 |

**Figure S7K Correlation Coeffient Contrast**

| control | test | control_N | test_N | effect_size | difference | bca_low | bca_high | pvalue_mann_whitney |
|---|---|---|---|---|---|---|---|---|
| 0 | 24 | 136 | 136 | mean difference | -0.09 | -0.14 | -0.04 | 0.0 |
| 0 | 48 | 136 | 136 | mean difference | -0.06 | -0.12 | -0.0 | 0.03 |

**Figure S7L Explained Variance**

| | |
|---|---|
| 1.0 | 0.28 |
| 2.0 | 0.44 |
| 3.0 | 0.58 |
| 4.0 | 0.65 |
| 5.0 | 0.72 |
| 6.0 | 0.78 |
| 7.0 | 0.82 |
| 8.0 | 0.86 |
| 9.0 | 0.9 |
| 10.0 | 0.93 |
| 11.0 | 0.95 |
| 12.0 | 0.97 |
| 13.0 | 0.98 |
| 14.0 | 0.99 |
| 15.0 | 1.0 |
| 16.0 | 1.0 |
| 17.0 | 1.0 |

**Figure S8**

**Figure S8A, B**

| Slope | r |
|---|---|
| -0.83 | -0.86 |
| 0.50 | 0.43 |
| 0.82 | 0.86 |
| 0.02 | 0.13 |
| -0.23 | -0.46 |
| 0.47 | 0.62 |
| -0.52 | -0.76 |
| -0.42 | -0.77 |
| 0.38 | 0.59 |
| -0.45 | -0.68 |
| 0.35 | 0.70 |
| -0.01 | -0.02 |
| 0.41 | 0.39 |
| -0.04 | -0.09 |
| -0.53 | -0.89 |
| 0.29 | 0.60 |
| 0.29 | 0.48 |
| 0.24 | 0.73 |
| 0.58 | 0.91 |
| -0.58 | -0.91 |
| 1.00 | 1.00 |
| -1.00 | -1.00 |

**Figure S8C-F**

| L24 | L48 | H24 | H48 | M24 | M48 | Metric |
|---|---|---|---|---|---|---|
| 1.01 | 2.25 | 0.53 | 1.56 | 0.78 | 1.91 | Volume |
| 0.75 | 1.62 | 0.24 | 1.0 | 0.5 | 1.3 | Duration |
| 1.04 | 1.56 | 0.56 | 0.96 | 0.84 | 1.28 | Meal Duration |
| 0.94 | 1.59 | 0.4 | 0.93 | 0.67 | 1.26 | Feed Speed |
| 0.93 | 1.54 | 0.44 | 0.93 | 0.74 | 1.25 | Meal Size |
| 0.32 | 0.76 | -0.14 | 0.24 | 0.11 | 0.51 | Food Port Occupancy |
| 0.41 | 0.59 | -0.07 | 0.04 | 0.18 | 0.32 | Height |
| 0.14 | 0.57 | -0.31 | 0.02 | -0.09 | 0.29 | Count |
| 0.41 | 0.42 | -0.12 | -0.19 | 0.16 | 0.13 | Duringfeed Speed Ratio |
| 0.86 | 0.33 | 0.44 | -0.21 | 0.65 | 0.05 | Falls |
| -0.04 | -0.03 | -0.66 | -0.69 | -0.37 | -0.37 | Prefeed Speed |
| -0.23 | -0.31 | -0.66 | -0.63 | -0.5 | -0.49 | Ctrl Port Occupancy |
| -0.27 | -0.34 | -0.62 | -0.68 | -0.45 | -0.52 | Speed |
| -0.57 | -0.37 | -1.16 | -1.03 | -0.86 | -0.7 | Postfeed Speed |
| -0.1 | -0.37 | -0.86 | -1.05 | -0.46 | -0.73 | Perifeed Speed Ratio |
| -0.1 | -0.57 | -0.58 | -1.13 | -0.34 | -0.85 | Latency |
| -0.54 | -0.74 | -1.08 | -1.24 | -0.84 | -1.0 | Duringfeed Speed |

**Figure S8G NPF>Chr**

| | Volume | Duration | Meal Duration | Feed Speed | Meal Size | Food Port Occupancy | Height | Count | Duringfeed Speed Ratio | Falls | Prefeed Speed | Ctrl Port Occupancy | Speed | Postfeed Speed | Perifeed Speed Ratio | Latency | Duringfeed Speed |
|---|---|---|---|---|---|---|---|---|---|---|---|---|---|---|---|---|---|
| mean 24 | 1.29 | 1.96 | 1.47 | 0.35 | 0.51 | 1.06 | -0.43 | 1.56 | -1.06 | 0.31 | 1.01 | -0.09 | 2.06 | 1.21 | -0.02 | -0.81 | -0.39 |
| bca_low 24 | 0.68 | 1.37 | 0.89 | -0.26 | -0.1 | 0.44 | -1.02 | 0.92 | -2.23 | -0.28 | 0.27 | -0.95 | 1.38 | 0.52 | -0.8 | -1.41 | -1.25 |
| bca_high 24 | 1.85 | 2.56 | 2.02 | 0.87 | 1.06 | 1.55 | 0.2 | 2.18 | -0.38 | 1.03 | 1.75 | 0.41 | 2.73 | 1.93 | 0.8 | -0.2 | 0.39 |

**Figure S8G NPF>ACR1**

| | Volume | Duration | Meal Duration | Feed Speed | Meal Size | Food Port Occupancy | Height | Count | Duringfeed Speed Ratio | Falls | Prefeed Speed | Ctrl Port Occupancy | Speed | Postfeed Speed | Perifeed Speed Ratio | Latency | Duringfeed Speed |
|---|---|---|---|---|---|---|---|---|---|---|---|---|---|---|---|---|---|
| mean 24 | -0.06 | 0.39 | -0.16 | -0.47 | -0.38 | -0.12 | 0.42 | 0.21 | 0.13 | -0.58 | 0.65 | -0.74 | 0.52 | 0.2 | -0.4 | -0.24 | 0.08 |
| bca_low 24 | -1.04 | -0.4 | -1.21 | -1.43 | -1.46 | -1.05 | -0.6 | -0.37 | -0.79 | -1.52 | 0.09 | -1.96 | -0.01 | -0.44 | -1.38 | -1.14 | -0.76 |
| bca_high 24 | 0.89 | 1.21 | 0.99 | 0.52 | 0.7 | 0.7 | 1.26 | 0.96 | 1.15 | 0.35 | 1.31 | -0.01 | 1.15 | 0.82 | 0.39 | 0.63 | 0.89 |

**Figure S8F AKH>Chr**

| | Volume | Duration | Meal Duration | Feed Speed | Meal Size | Food Port Occupancy | Height | Count | Duringfeed Speed Ratio | Falls | Prefeed Speed | Ctrl Port Occupancy | Speed | Postfeed Speed | Perifeed Speed Ratio | Latency | Duringfeed Speed |
|---|---|---|---|---|---|---|---|---|---|---|---|---|---|---|---|---|---|
| mean 24 | -0.58 | -0.36 | -0.47 | -0.64 | -0.12 | 0.05 | -0.5 | -0.0 | 0.49 | 0.47 | 0.47 | 0.3 | 0.67 | 0.74 | 0.82 | 0.43 | 1.11 |
| bca_low 24 | -1.32 | -1.08 | -1.01 | -1.21 | -0.58 | -0.57 | -1.1 | -0.77 | -0.5 | -0.16 | -0.45 | -0.54 | -0.09 | -0.26 | -0.29 | -0.26 | 0.19 |
| bca_high 24 | 0.14 | 0.35 | 0.19 | -0.01 | 0.6 | 0.62 | 0.14 | 0.73 | 2.04 | 1.13 | 1.42 | 1.02 | 1.44 | 1.69 | 2.01 | 1.1 | 2.14 |

**Figure S8F AKH>ACR1**

| | Volume | Duration | Meal Duration | Feed Speed | Meal Size | Food Port Occupancy | Height | Count | Duringfeed Speed Ratio | Falls | Prefeed Speed | Ctrl Port Occupancy | Speed | Postfeed Speed | Perifeed Speed Ratio | Latency | Duringfeed Speed |
|---|---|---|---|---|---|---|---|---|---|---|---|---|---|---|---|---|---|
| mean 24 | 0.29 | 0.37 | 0.28 | 0.02 | 0.11 | 0.48 | -0.06 | 0.28 | 0.51 | 0.04 | -0.79 | -0.04 | 0.22 | -0.83 | -0.26 | 0.02 | -0.86 |
| bca_low 24 | -0.54 | -0.21 | -0.28 | -0.64 | -0.48 | -0.0 | -0.69 | -0.58 | -0.39 | -0.55 | -1.78 | -0.64 | -0.77 | -1.85 | -1.07 | -0.69 | -2.34 |
| bca_high 24 | 1.04 | 1.05 | 0.88 | 0.88 | 0.83 | 1.09 | 0.56 | 1.03 | 1.44 | 0.59 | 0.28 | 0.41 | 1.05 | 0.27 | 0.54 | 0.69 | 0.3 |

**Figure S8H Dilp2>Chr**

| | Volume | Duration | Meal Duration | Feed Speed | Meal Size | Food Port Occupancy | Height | Count | Duringfeed Speed Ratio | Falls | Prefeed Speed | Ctrl Port Occupancy | Speed | Postfeed Speed | Perifeed Speed Ratio | Latency | Duringfeed Speed |
|---|---|---|---|---|---|---|---|---|---|---|---|---|---|---|---|---|---|
| mean 24 | 1.25 | 1.13 | -0.31 | 0.05 | -0.17 | 0.42 | -0.02 | 1.05 | 0.24 | -0.44 | 0.08 | -0.12 | 0.51 | -0.21 | -0.25 | -0.52 | -0.01 |
| bca_low 24 | 0.57 | 0.44 | -1.04 | -0.71 | -0.89 | -0.45 | -0.72 | 0.35 | -0.4 | -1.23 | -0.66 | -0.99 | -0.2 | -0.99 | -0.96 | -1.22 | -0.77 |
| bca_high 24 | 1.97 | 1.88 | 0.41 | 0.72 | 0.51 | 1.09 | 0.69 | 1.78 | 1.08 | 0.28 | 0.81 | 0.54 | 1.21 | 0.52 | 0.49 | 0.22 | 0.73 |

**Figure S8H**
**Dilp2>ACR1**

| | Volume | Duration | Meal Duration | Feed Speed | Meal Size | Food Port Occupancy | Height | Count | Duringfeed Speed Ratio | Falls | Prefeed Speed | Ctrl Port Occupancy | Speed | Postfeed Speed | Perifeed Speed Ratio | Latency | Duringfeed Speed |
|---|---|---|---|---|---|---|---|---|---|---|---|---|---|---|---|---|---|
| mean 24 | 0.59 | 0.44 | 0.34 | 0.08 | 0.52 | 0.6 | 0.31 | 0.0 | 0.52 | -0.0 | 0.11 | 0.11 | -0.14 | -0.13 | -0.2 | 0.19 | -0.34 |
| bca_low 24 | -0.25 | -0.55 | -0.47 | -0.72 | -0.23 | -0.52 | -0.62 | -0.89 | -0.39 | -1.12 | -0.73 | -0.48 | -0.82 | -0.83 | -1.43 | -0.47 | -1.18 |
| bca_high 24 | 1.43 | 1.44 | 1.37 | 1.05 | 1.59 | 1.69 | 1.19 | 0.88 | 1.55 | 0.96 | 0.99 | 1.11 | 0.67 | 0.62 | 0.77 | 1.06 | 0.39 |

**Figure S8E**
**Trhn-AND-tsh>Chr**

| | Volume | Duration | Meal Duration | Feed Speed | Meal Size | Food Port Occupancy | Height | Count | Duringfeed Speed Ratio | Falls | Prefeed Speed | Ctrl Port Occupancy | Speed | Postfeed Speed | Perifeed Speed Ratio | Latency | Duringfeed Speed |
|---|---|---|---|---|---|---|---|---|---|---|---|---|---|---|---|---|---|
| mean 24 | -1.35 | -0.64 | -0.33 | -0.64 | -0.61 | -0.17 | -0.16 | -1.12 | -0.05 | 0.25 | 0.01 | -0.79 | -1.05 | -0.14 | -0.24 | -0.15 | 0.03 |
| bca_low 24 | -1.94 | -1.15 | -1.05 | -1.13 | -1.46 | -0.66 | -0.66 | -1.97 | -0.49 | -0.44 | -0.56 | -1.5 | -1.88 | -0.74 | -0.71 | -0.47 | -0.45 |
| bca_high 24 | -0.76 | -0.13 | 0.14 | -0.17 | -0.11 | 0.35 | 0.35 | -0.33 | 0.52 | 1.05 | 0.6 | -0.12 | -0.25 | 0.51 | 0.26 | 0.18 | 0.55 |

**Figure S8E**
**Trhn-AND-tsh>ACR1**

| | Volume | Duration | Meal Duration | Feed Speed | Meal Size | Food Port Occupancy | Height | Count | Duringfeed Speed Ratio | Falls | Prefeed Speed | Ctrl Port Occupancy | Speed | Postfeed Speed | Perifeed Speed Ratio | Latency | Duringfeed Speed |
|---|---|---|---|---|---|---|---|---|---|---|---|---|---|---|---|---|---|
| mean 24 | 0.27 | -0.01 | 1.27 | 1.08 | 1.28 | -0.57 | -0.07 | -0.4 | -0.13 | -0.41 | 0.0 | -0.49 | -0.93 | 0.09 | 0.44 | -0.87 | -0.51 |
| bca_low 24 | -0.32 | -0.57 | 0.72 | 0.5 | 0.71 | -1.18 | -0.64 | -1.0 | -0.8 | -1.02 | -0.64 | -1.1 | -1.55 | -0.56 | -0.15 | -1.46 | -1.24 |
| bca_high 24 | 0.84 | 0.56 | 1.94 | 1.68 | 1.96 | -0.04 | 0.53 | 0.18 | 0.49 | 0.16 | 0.64 | 0.11 | -0.31 | 0.75 | 1.13 | -0.32 | 0.17 |

### Figure 6

#### Figure 6A Brain Normalized 5HT

| control genotype | test genotype | control condition | test condition | control_N | test_N | effect_size | difference | bca_low | bca_high | pvalue_permutation |
|---|---|---|---|---|---|---|---|---|---|---|
| Fed W1118 | Starved W1118 | Fed | Starved | 10 | 11 | mean difference | 0.74 | 0.42 | 1.03 | 0.0 |

#### Figure 6B VNC Normalized 5HT

| control genotype | test genotype | control condition | test condition | control_N | test_N | effect_size | difference | bca_low | bca_high | pvalue_permutation |
|---|---|---|---|---|---|---|---|---|---|---|
| Fed W1118 | Starved W1118 | Fed | Starved | 10 | 11 | mean difference | 0.63 | 0.37 | 0.96 | 0.0 |

#### Figure 6E Trhi

| control | test | control_N | test_N | effect_size | difference | bca_low | bca_high | pvalue_mann_whitney |
|---|---|---|---|---|---|---|---|---|
| Ctrl_TrhTrhi_Combined Controls | Test_TrhTrhi_Trh-Gal4>Trhi | 60 | 29 | mean difference | 85.29 | 54.83 | 117.56 | 0.0 |
| Ctrl_TrhVTrhi_Combined Controls | Test_TrhVTrhi_Trh-AND-tsh-Gal4>Trhi | 53 | 35 | mean difference | 68.83 | 13.77 | 119.0 | 0.01 |

#### Figure 6F Gluti Sut2i

| control | test | control_N | test_N | effect_size | difference | bca_low | bca_high | pvalue_mann_whitney |
|---|---|---|---|---|---|---|---|---|
| Ctrl_TrhGluti_Combined Controls | Test_TrhGluti_Trh-Gal4>Glut1i | 54 | 35 | mean difference | -23.04 | -64.74 | 28.21 | 0.23 |
| Ctrl_TrhVGluti_Combined Controls | Test_TrhVGluti_Trh-AND-tsh-Gal4>Glut1i | 31 | 28 | mean difference | 40.99 | -10.92 | 96.67 | 0.08 |
| Ctrl_TrhSut2i_Combined Controls | Test_TrhSut2i_Trh-Gal4>sut2i | 31 | 28 | mean difference | -57.61 | -154.83 | 27.19 | 0.98 |
| Ctrl_TrhVSut2i_Combined Controls | Test_TrhVSut2i_Trh-AND-tsh-Gal4>sut2i | 82 | 53 | mean difference | 56.97 | 29.69 | 83.47 | 0.0 |

### Figure S10

#### Figure S10A Fed/Starved/Refed Brain Normalized 5HT

| control genotype | test genotype | control condition | test condition | control_N | test_N | effect_size | difference | bca_low | bca_high | pvalue_permutation |
|---|---|---|---|---|---|---|---|---|---|---|
| w1118 fed brain | w1118 starved brain | Fed | Starved | 7 | 8 | mean difference | 0.2130358795 | -0.00620133131 | 0.5418938438 | 0.1926 |
| w1118 fed brain | w1118 refed brain | Fed | Refed | 7 | 7 | mean difference | 0.02921209364 | -0.209629119 | 0.2807703873 | 0.8262 |

#### Figure S10B Fed/Starved/Refed VNC Normalized 5HT

| control genotype | test genotype | control condition | test condition | control_N | test_N | effect_size | difference | bca_low | bca_high | pvalue_permutation |
|---|---|---|---|---|---|---|---|---|---|---|
| w1118 fed vnc | w1118 starved vnc | Fed | Starved | 6 | 7 | mean difference | 0.34695408 | -0.09400835124 | 0.9018183469 | 0.258 |
| w1118 fed vnc | w1118 refed brain | Fed | Refed | 6 | 5 | mean difference | 0.03239131212 | -0.3305545389 | 0.4016859178 | 0.8866 |

#### Figure S10C Trh>Trhi Separate Controls

| control genotype | test genotype | control condition | test condition | control_N | test_N | effect_size | difference | bca_low | bca_high | pvalue_permutation |
|---|---|---|---|---|---|---|---|---|---|---|
| Trhi/W | Trh-Gal4/W | RNAi/W | Trh-Gal4/W | 30 | 30 | mean difference | 75.30433333 | 37.9705 | 108.2328333 | 0 |
| Trhi/W | Trh-Gal4>Trhi | RNAi/W | Trh-Gal4>RNAi | 30 | 29 | mean difference | 122.9385816 | 89.07505287 | 154.0116506 | 0 |

#### Figure S10D TrhV>Trhi Separate Controls

| control genotype | test genotype | control condition | test condition | control_N | test_N | effect_size | difference | bca_low | bca_high | pvalue_permutation |
|---|---|---|---|---|---|---|---|---|---|---|
| Trhi/W | TrhV-Gal4/W | RNAi/W | Trh-AND-tsh-Ga | 29 | 24 | mean difference | 91.84683477 | 34.71646121 | 167.8001609 | 0.0054 |
| Trhi/W | TrhV-Gal4>Trhi | RNAi/W | Trh-AND-tsh-Ga | 29 | 35 | mean difference | 110.4218788 | 59.00497833 | 160.0885685 | 0.0002 |

#### Figure S10E Trh>Gluti Separate Controls

| control genotype | test genotype | control condition | test condition | control_N | test_N | effect_size | difference | bca_low | bca_high | pvalue_permutation |
|---|---|---|---|---|---|---|---|---|---|---|
| Glut1i/W | Trh-Gal4/W | RNAi/W | Trh-Gal4/W | 25 | 29 | mean difference | 122.6049766 | 77.91926207 | 167.0865628 | 0 |
| Glut1i/W | Trh-Gal4>Trhi | RNAi/W | Trh-Gal4>RNAi | 25 | 35 | mean difference | 42.79928 | -3.885108571 | 91.62305714 | 0.1098 |

#### Figure S10F TrhV>Gluti Separate Controls

| control genotype | test genotype | control condition | test condition | control_N | test_N | effect_size | difference | bca_low | bca_high | pvalue_permutation |
|---|---|---|---|---|---|---|---|---|---|---|
| Glut1i/W | TrhV-Gal4/W | RNAi/W | Trh-AND-tsh-Ga | 16 | 15 | mean difference | 116.4641708 | 55.01127083 | 178.5515958 | 0.0008 |
| Glut1i/W | TrhV-Gal4>Glut1i | RNAi/W | Trh-AND-tsh-Ga | 16 | 28 | mean difference | 97.3459375 | 47.75407143 | 144.0803571 | 0.0008 |

#### Figure S10G Trh>Sut2i Separate Controls

| control genotype | test genotype | control condition | test condition | control_N | test_N | effect_size | difference | bca_low | bca_high | pvalue_permutation |
|---|---|---|---|---|---|---|---|---|---|---|
| sut2i/W | Trh-Gal4/W | RNAi/W | Trh-Gal4/W | 15 | 16 | mean difference | 335.4671583 | 248.8878292 | 449.1937625 | 0 |
| sut2i/W | Trh-Gal4>Trhi | RNAi/W | Trh-Gal4>RNAi | 15 | 28 | mean difference | 115.5308548 | 73.32935238 | 171.854169 | 0.0002 |

#### Figure S10H TrhV>Sut2i Separate Controls

| control genotype | test genotype | control condition | test condition | control_N | test_N | effect_size | difference | bca_low | bca_high | pvalue_permutation |
|---|---|---|---|---|---|---|---|---|---|---|
| sut2i/W | TrhV-Gal4/W | RNAi/W | Trh-AND-tsh-Ga | 40 | 42 | mean difference | 79.11949881 | 46.2926619 | 105.9037393 | 0 |
| sut2i/W | TrhV-Gal4>sut2i | RNAi/W | Trh-AND-tsh-Ga | 40 | 53 | mean difference | 97.49482406 | 63.61688821 | 123.9037788 | 0 |
